# Targeted Protein Degradation through Recruitment of the CUL4A Complex Adaptor Protein DDB1

**DOI:** 10.1101/2023.08.11.553046

**Authors:** Margot Meyers, Sabine Cismoski, Anoohya Panidapu, Barbara Chie-Leon, Daniel K. Nomura

**Affiliations:** Department of Chemistry, University of California, Berkeley, Berkeley, CA 94720 USA; Novartis-Berkeley Translational Chemical Biology Institute; Innovative Genomics Institute, Berkeley, CA 94720 USA; Novartis Institutes for BioMedical Research, Emeryville, CA 94608 USA; Department of Molecular and Cell Biology, University of California, Berkeley, Berkeley, CA 94720 USA

**Keywords:** activity-based protein profiling, targeted protein degradation, DDB1, CUL4A, BRD4, androgen receptor, AR, covalent, chemoproteomics

## Abstract

Targeted protein degradation has arisen as a powerful therapeutic modality for eliminating proteins. Thus far, most heterobifunctional Proteolysis Targeting Chimeras (PROTACs) have utilized recruiters against substrate receptors of Cullin RING E3 ubiquitin ligases, such as cereblon and VHL. However, previous studies have surprisingly uncovered molecular glue degraders that exploit a CUL4A adaptor protein DDB1 to degrade neosubstrate proteins. Here, we sought to investigate whether DDB1 recruiters can be discovered that can be exploited for PROTAC applications. We utilized activity-based protein profiling and cysteine chemoproteomic screening to identify a covalent recruiter that targets C173 on DDB1 and exploited this recruiter to develop PROTACs against BRD4 and androgen receptor (AR). We demonstrated that the BRD4 PROTAC results in selective degradation of the short BRD4 isoform over the long isoform in a proteasome, NEDDylation, and DDB1-dependent manner. We also demonstrated degradation of AR with the AR PROTAC in prostate cancer cells. Our study demonstrated that covalent chemoproteomic approaches can be used to discover recruiters against Cullin RING adapter proteins and that these recruiters can be used for PROTAC applications to degrade neo-substrates.

## Introduction

Targeted protein degradation (TPD) has arisen as a powerful approach for eliminating disease-causing proteins ^1^. Currently there are two major approaches for TPD: Proteolysis Targeting Chimeras (PROTACs) and molecular glue degraders. These approaches utilize either heterobifunctional or monovalent small-molecules to induce the proximity of E3 ubiquitin ligases with neo-substrates, resulting in their ubiquitination and subsequent degradation through the proteasome ^2–5^. While there are >600 E3 ligases in the human proteome, most degraders have exploited only a small number of E3 ligases. To tackle the full scope of undrugged therapeutic targets,there is a need for the discovery of additional recruiters of the ubiquitin-proteasome system (UPS) that can be used in TPD applications ^6^.

Currently most PROTACs have exploited substrate receptors of Cullin RING E3 ubiquitin ligases, including most notably cereblon and VHL, and more recently DCAF1, DCAF11, DCAF16, KEAP1, and FEM1B ^7–14^. There have also been several recruiters that have been discovered against RING E3 ligases, including RNF4 and RNF114 ^15–17^. Many of these E3 ligases that have been exploited to date are not essential to cell viability. Cancer cells treated with degraders utilizing non-essential E3 ligases may select for the loss of the genes encoding those ligases, rendering the degrader ineffective over time. Developing recruiters against core and essential components of the UPS may prevent or slow these potential resistance mechanisms. Towards this goal, recent studies have uncovered novel recruiters against essential E2 ubiquitin-conjugating enzymes and even the proteasome itself which can be used in heterobifunctional degraders, but it has not yet been shown whether core adaptor proteins of Cullin RING E3 ubiquitin ligases, such as DDB1, SKP1, or ELOB/ELOC in CUL1-CUL7 E3 ligases are also exploitable for PROTAC applications^18,19^.

Intriguingly, Slabicki and Ebert *et al* recently demonstrated that the cyclin-dependent kinase inhibitor CR8 acts as a molecular glue between CDK12-cyclin K and the CUL4 adaptor protein DDB1^20^. This molecule resulted in the degradation of cyclin K even in the absence of a substrate receptor, suggesting that DDB1 could be used to drive TPD of neo-substrates. In this study, we further show the utility of DDB1 for TPD through the discovery of a novel DDB1 ligand and its elaboration into PROTACs targeting BRD4 and AR.

## Results

### Covalent Ligand Screening Against DDB1 to Identify a DDB1 Recruiter

We used activity-based protein profiling (ABPP) to screen a library of cysteine-reactive small-molecules to identify covalent ligands that would displace cysteine-reactive fluorescent probe labeling of pure human DDB1 protein **(Figure S1; Table S1)**. From this screen, we identified a chloroacetamide hit with a pyrazoline core, EN223, that potently bound to DDB1 in a dose-responsive manner **(Figure S2A).** While EN223 was a promising hit, we had observed EN223 as a hit across previous screens, including an E3 ligase RNF114 ^17^, suggesting this compound may be promiscuous. Interestingly, we had previously identified another pyrazoline-based chloroacetamide EN219 as the primary hit against RNF114, but this compound was not a hit against DDB1 **(Figure S1)** ^17^. Based on the EN223 hit, we tested five additional pyrazoline-based covalent ligands against DDB1 and identified MM-02-57 as a promising alternate binder that showed comparable binding to DDB1 as EN223 **(Figure S2B, Figure 1A-1B).** Importantly, MM-02-57 did not bind to RNF114 **(Figure 1C)**. We further synthesized an alkyne-functionalized probe of this hit, MM-02-54, and demonstrated that this probe also bound to DDB1 comparably to MM-02-057, indicating that extending off the phenyl substituent may be a favorable exit vector for generating a PROTAC **(Figure 1D-1E).** We also demonstrated that MM-02-54 directly covalently labels pure DDB1 protein and that this labeling was dose-responsively displaced by MM-02-57 *in vitro* **(Figure 1F).** We mapped the site of modification of MM-02-57 to cysteine C173 in DDB1 by liquid chromatography-mass spectrometry (LC-MS/MS) analysis of MM-02-57 modified DDB1 tryptic peptides **(Figure 1G)**. We next sought to confirm that MM-02-57 does not disrupt overall ubiquitination mediated by the CUL4A-DDB1 complex—a requisite for use of this ligand as a DDB1 recruiter. We reconstituted the CUL4A complex with CRBN and thalidomide-mediated ubiquitination of IKZF1 and showed that MM-02-57 does not impair IKZF1 ubiquitination **(Figure 1H)**.

**Figure 1.**
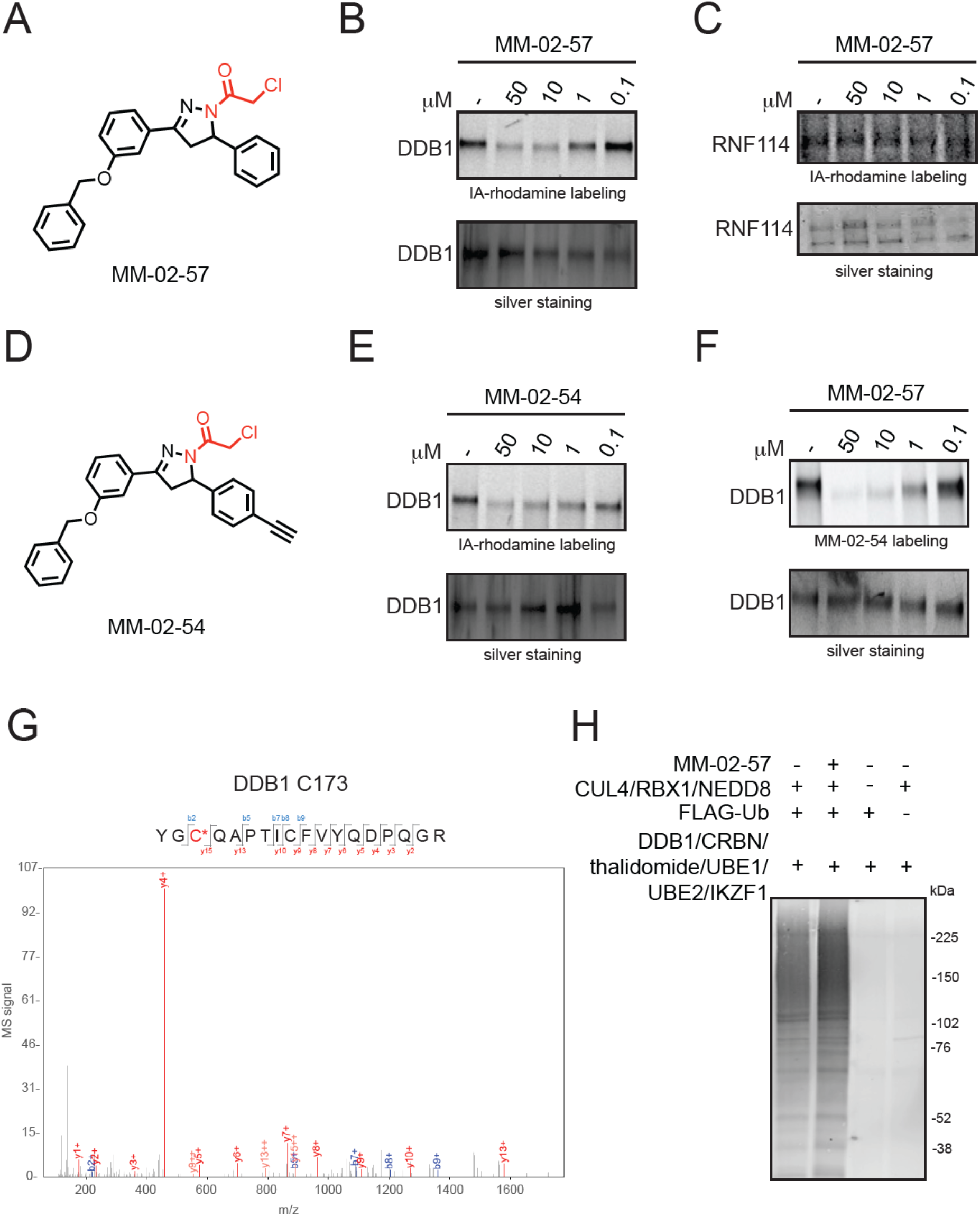
Characterization of DDB1 covalent recruiter MM-02-57. **(A)** Structure of DDB1 covalent ligand hit MM-02-57. **(B)** Gel-based ABPP of MM-02-57 against rhodamine-functionalized cysteine-reactive iodoacetamide probe (IA-rhodamine) labeling of pure DDB1 protein. **(C)** Gel-based ABPP of MM-02-57 against IA-rhodamine labeling of pure RNF114. **(D)** Structure of alkyne-functionalized probe MM-02-54. **(E)** Gel-based ABPP of MM-02-57 against IA-rhodamine labeling of pure DDB1 protein. For gels in **(B, C, E)**, DDB1 or RNF114 was pre-incubated with DMSO or the covalent ligand for 30 min prior to IA-rhodamine labeling (100 nM for **(B, E)** and 10 μM for **(C)**) for 30 min after which proteins were resolved by SDS/PAGE and IA-rhodamine labeling was visualized by in-gel fluorescence and loading was assessed by silver staining. **(F)** Gel-based ABPP of MM-02-57 against MM-02-54 labeling of pure DDB1 protein. Probe-labeled proteins were subjected to CuAAC-mediated appendage of a rhodamine-azide after which proteins were resolved on SDS/PAGE and probe labeling was assessed by in-gel fluorescence and loading was assessed by silver staining. **(G)** LC-MS/MS analysis of MM-02-57 modification on DDB1. DDB1 was labeled with MM-02-57 (50 μM) for 30 min and subsequently tryptically digested for LC-MS/MS analysis. Shown are the MS/MS data for the MM-02-57-modified C173-containing DDB1 peptide. **(H)** Reconstituted CUL4A ubiquitination assay with MM-02-057. Various components of the complex were pre-incubated with DMSO vehicle or MM-02-57 (50 μM) for 30 min, after which FLAG-Ubiquitinated proteins were detected by Western blotting. Gels shown in **(B, C, E, F, H)** are representative gels from n=3 biologically independent replicates/group.

### Assessing DDB1 Engagement and Overall Selectivity of DDB1 Recruiter in Cells

We next determined whether MM-02-57 or MM-02-54 engaged DDB1 in cells and assessed the proteome-wide cysteine reactivity of MM-02-57. Using the MM-02-54 probe in cells, we showed that we could pulldown DDB1, but not unrelated targets such as GAPDH, from cells upon *ex situ* capture of MM-02-54 labeled DDB1 through copper-catalyzed azide-alkyne cycloaddition (CuAAC) with an azide-functionalized biotin and subsequent avidin capture and elution **(Figure 2A)**. These data showed that MM-02-54 directly engaged DDB1 in cells. We next performed cysteine chemoproteomic profiling using the alkyne-functionalized iodoacetamide probe, the isotopically labeled desthiobiotin azide tags, and activity-based protein profiling (isoDTB-ABPP) ^21–23^ to map the proteome-wide cysteine-reactivity of MM-02-057. Out of 8974 cysteines quantified, we identified only 24 targets that showed control versus treated log_2_ ratios of at least 3 where in DDB1 C173 showed an absolute ratio of 8.4, indicating 88 % engagement of DDB1 **(Figure 2B; Table S2)**. These data also confirmed that C173 was the primary site engaged by MM-02-057 in cells. Among the 24 off-targets, we identified two additional E3 ligase targets that are involved in ubiquitin-mediated degradation—KEAP1 C288 and ZNRF2 C236 **(Figure 2B; Table S2)**. Given that DDB1 is part of the CUL4 E3 ligase class, KEAP1 is a member of the CUL3 family, and ZNRF2 is not part of any Cullin family E3 ligase, we conjectured that we could mechanistically tease apart DDB1-mediated degradation events resulting from subsequent experiments with MM-02-057-based PROTACs. Furthermore, we also confirmed in our isoDTB-ABPP dataset that C8 of RNF114 was not engaged by MM-02-057, showing a ratio of 1.18 and not statistically significant, given the structural similarities between our previously discovered RNF114 recruiter EN219 and MM-02-057 **(Table S2).**

**Figure 2.**
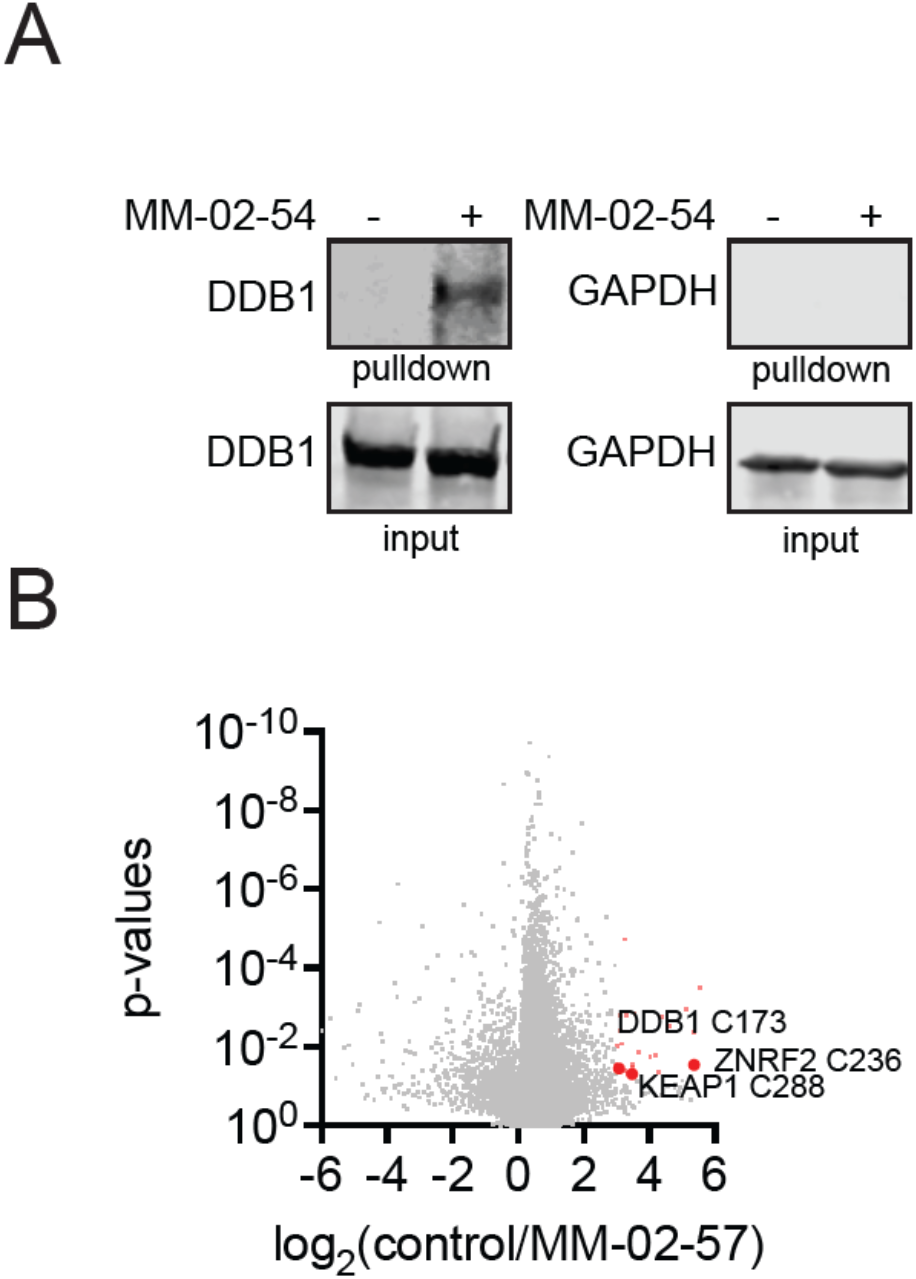
DDB1 target engagement and selectivity of MM-02-057 in cells. **(A)** MM-02-54 target engagement of DDB1 in cells. HEK293T cells were treated with DMSO vehicle or MM-02-54 (xxx μM) for xxx hours, after which resulting cell lysates were subjected to CuAAC-mediated appendage of an azide-functionalized biotin handle onto probe-labeled proteins. Probe-labeled proteins were avidin-enriched, eluted, and resolved on SDS/PAGE and DDB1 and an unrelated protein GAPDH input and pulldown were detected by Western blotting. **(B)** IsoDTB-ABPP analysis of MM-02-57 in HEK293T cells. HEK293T cells were treated with DMSO vehicle or MM-02-57 (50 μM) for 4 hours after which resulting control and treated lysates were labeled with an alkyne-functionalized iodoacetamide probe (IA-alkyne) for 1 h. Probe-labeled proteins were subjected to CuAAC to append either an isotopically light (for control) or heavy (for treated) azide-functionalized desthiobiotin handle, after which probe-labeled proteins were avidin-enriched, digested with trypsin, and probe-modified peptides were eluted and analyzed by LC-MS/MS. Control/treated probe-modified peptides were quantified and plotted. The points shown in red were targets that showed ratios of >8 with adjusted p-values <0.05. The full dataset can be found in **Table S2.** Data in **(A, B)** are from n=3 biologically independent replicates per group.

### BRD4 Degradation using MM-02-057-Based PROTACs

Having shown that MM-02-057 engages DDB1 in cells in a non-inhibitory manner, we next assessed whether this covalent DDB1 binder could be used as a recruiter in PROTAC applications. We synthesized six PROTACs linking our DDB1 recruiter MM-02-057 onto the BET bromodomain inhibitor JQ1 through either a C2, C3, C4, C5, C6, or C7 alkyl linker **(Figure 3A).** All 6 PROTACs degraded BRD4, but each degrader showed differential potency in degrading the target **(Figure 3B)**. Surprisingly, we observed that all six PROTACs only degraded the short isoform of BRD4 without affecting the long isoform. The selective degradation of only the short isoform of BRD4 is intriguing given previous studies that showed tumor suppressive roles of the long isoform of BRD4 and the oncogenic roles of the short BRD4 isoform ^24^. Also interestingly, only MM-03-42 bearing the C4 alkyl linker showed hook effects at the highest concentration tested **(Figure 3B)**. The most potent degrader was MM-02-08 possessing a C5 alkyl linker **(Figure 3B)**. Quantitative proteomic profiling of MM-02-08 did not show appreciable >2-fold loss of total BRD4 levels, given that we only showed loss of the less abundant short BRD4 isoform (data not shown).

**Figure 3.**
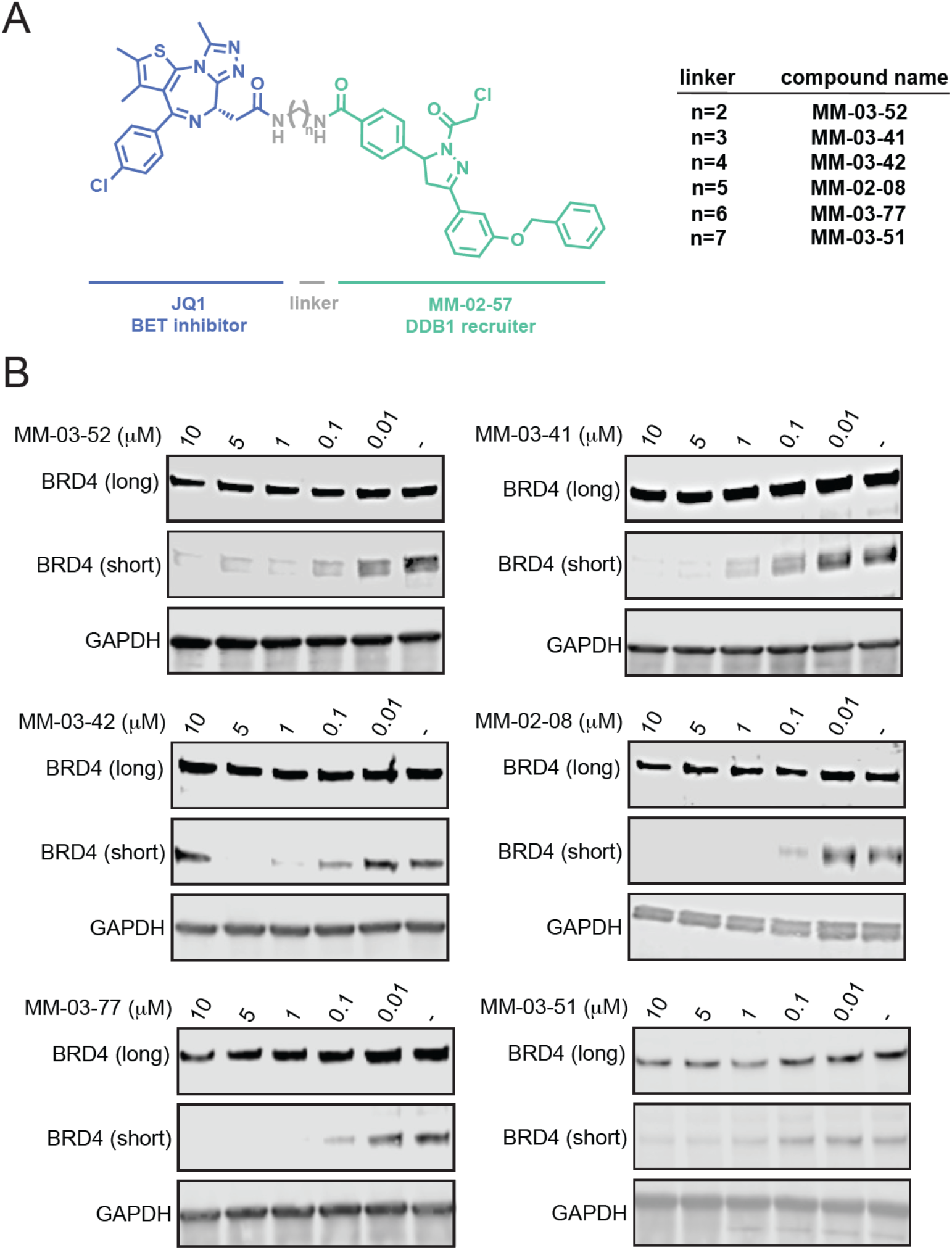
DDB1-based BRD4 PROTACs. **(A)** Structures of MM-02-057-based BRD4 PROTACs with varying linkers. **(B)** BRD4 degradation by MM-02-057 based BRD4 PROTACs. HEK293T cells were treated with DMSO vehicle or PROTAC for 24 h after which the long and short BRD4 isoforms and loading control GAPDH were assessed by Western blotting. All blots are representative of n=3 biologically independent replicates/group.

The loss of the short-BRD4 isoform from MM-02-08 treatment was attenuated by pre-treatment with both the proteasome inhibitor bortezomib (BTZ) and the NEDDylation inhibitor MLN4924 **(Figure 4A).** Confirming the contribution of DDB1 in the observed BRD4 degradation, we showed complete attenuation of BRD4 degradation upon DDB1 knockdown **(Figure 4B)**. We further demonstrated enhanced BRD4 ubiquitination only in the presence of the NEDDylated CUL4A/DDB1/UBE1/UBE2/RBX1 complex treated with MM-02-08 compared to vehicle-treated controls. BRD4 ubiquitination was not observed without addition of DDB1 or without the CUL4A complex **(Figure 4C)**. Interestingly, we observed this enhanced ubiquitination without the addition of any E3 ligase substrate receptor, indicating that these DDB1-based PROTACs may bypass the necessity for an E3 ligase substrate receptor, analogous to what was observed with the DDB1-based cyclin K molecular glue degrader CR8 ^20^.

**Figure 4.**
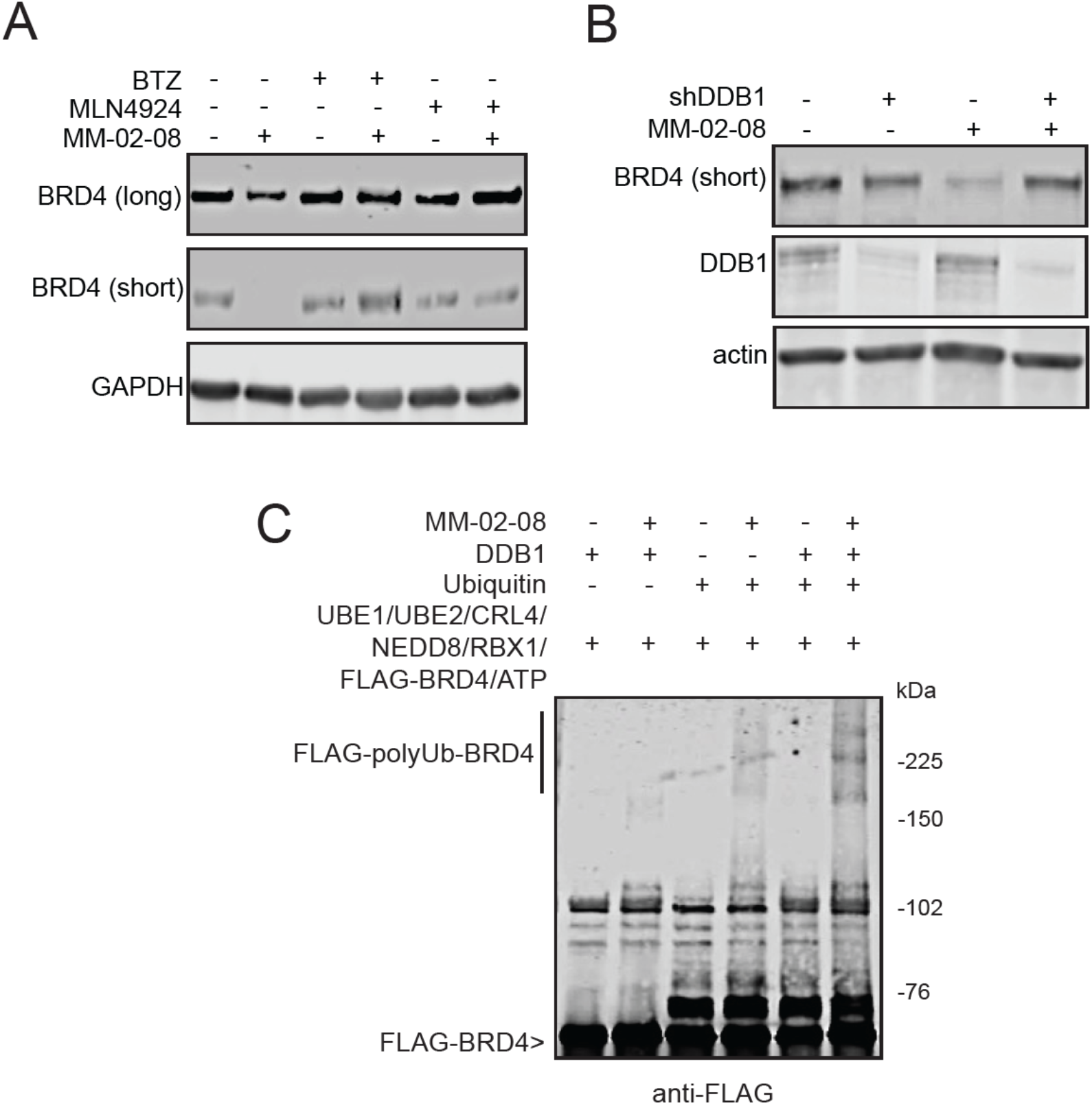
DDB1-dependence of BRD4 degradation. **(A)** Attenuation of MM-02-08-mediated BRD4 degradation with proteasome and NEDDylation inhibitors. HEK293T cells were treated with DMSO or BTZ (1 μM) or MLN4924 (1 μM) for 1 h prior to MM-02-08 (1 μM) treatment for 24 hours. BRD4 and loading control GAPDH were assessed by Western blotting. All blots are representative of n=3 biologically independent replicates/group. **(B)** DDB1 knockdown attenuates MM-02-08-mediated BRD4 degradation. Stable short-hairpin mediated control and DDB1 knockdown HEK293T cells were treated with DMSO vehicle or MM-02-08 (1 μM) treatment for 24 hours and BRD4, DDB1, and loading control actin were assessed by Western blotting. **(C)** Reconstitution of various components of the CUL4A complex with ubiquitin and BRD4 treated with DMSO vehicle or MM-02-08 (1 μM) treatment for 1 hour. FLAG-BRD4 levels, including higher molecular weight ubiquitinated FLAG-BRD4 levels were assessed by Western blotting. All blots are representative of n=3 biologically independent replicates/group.

As a negative control, the non-reactive analogs of MM-02-08, AP-01-104 expectedly did not show DDB1 binding and did not degrade BRD4 in cells **(Figure 5A-5C).** The chloroacetamide cysteine-reactive warhead, while suitable for cellular studies and generating tool compounds, is likely to be metabolically unstable and thus intractable for future drug discovery applications ^25^. As such, we also tested whether a more metabolically suitable warhead could be accommodated. While warhead swapping often compromises the activity of covalent ligands ^26^, we found that the acrylamide warhead-bearing counterpart of MM-02-08, MM-04-09, still bound to DDB1 and degraded BRD4 in cells **(Figure 5D-5F).**

**Figure 5.**
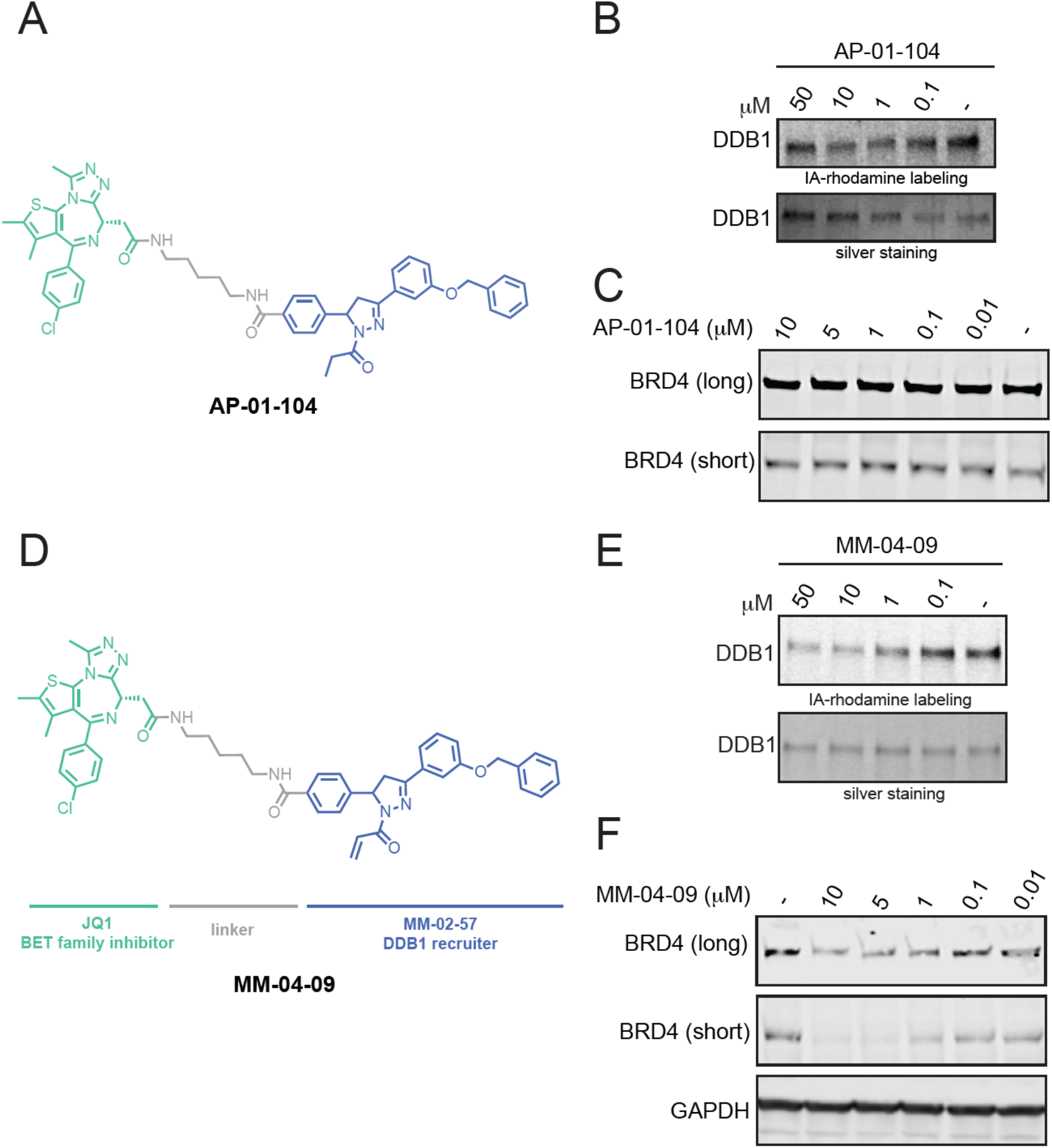
Analogs of BRD4 PROTACs. **(A)** Structure of a non-reactive BRD4 PROTAC, AP-01-104. **(B)** Gel-based ABPP of AP-01-104 against IA-rhodamine labeling of DDB1 pure protein. **(C)** BRD4 levels in HEK293T cells treated with DMSO vehicle or AP-01-104 for 24 h, assessed by Western blotting. **(D)** Acrylamide-bearing BRD4 PROTAC MM-04-09. **(E)** Gel-based ABPP of MM-04-09 against IA-rhodamine labeling of DDB1 pure protein. **(F)** BRD4 and loading control GAPDH levels in HEK293T cells treated with DMSO vehicle or MM-04-09 for 24 h, assessed by Western blotting. Gels and blots are representative of n=3 biologically independent replicates per group.

### Androgen Receptor (AR) Degradation using MM-02-057-Based PROTACs

While we demonstrated that MM-02-057 could be used to degrade BRD4, BRD4 is also one of the easiest targets to degrade with the PROTAC modality. We thus also tested our MM-02-057 handle could be used to degrade AR. We synthesized four PROTACs linking our MM-02-57 DDB1 recruiter to the AR-targeting moiety of the Arvinas AR PROTAC ARV-110^27^ with either no linker, or via a C2, C4, or C5 alkyl linker **(Figure 6A)**. While all four PROTACs showed some degree of AR degradation, the best degradation was observed with MM-03-73, utilizing a C2 alkyl linker **(Figure 6B).** While this degradation was modest compared to the efficacy of ARV-110, which exploits a cereblon recruiter, our data shows that DDB1 can be used to degrade neo-substrates beyond BRD4.

**Figure 6.**
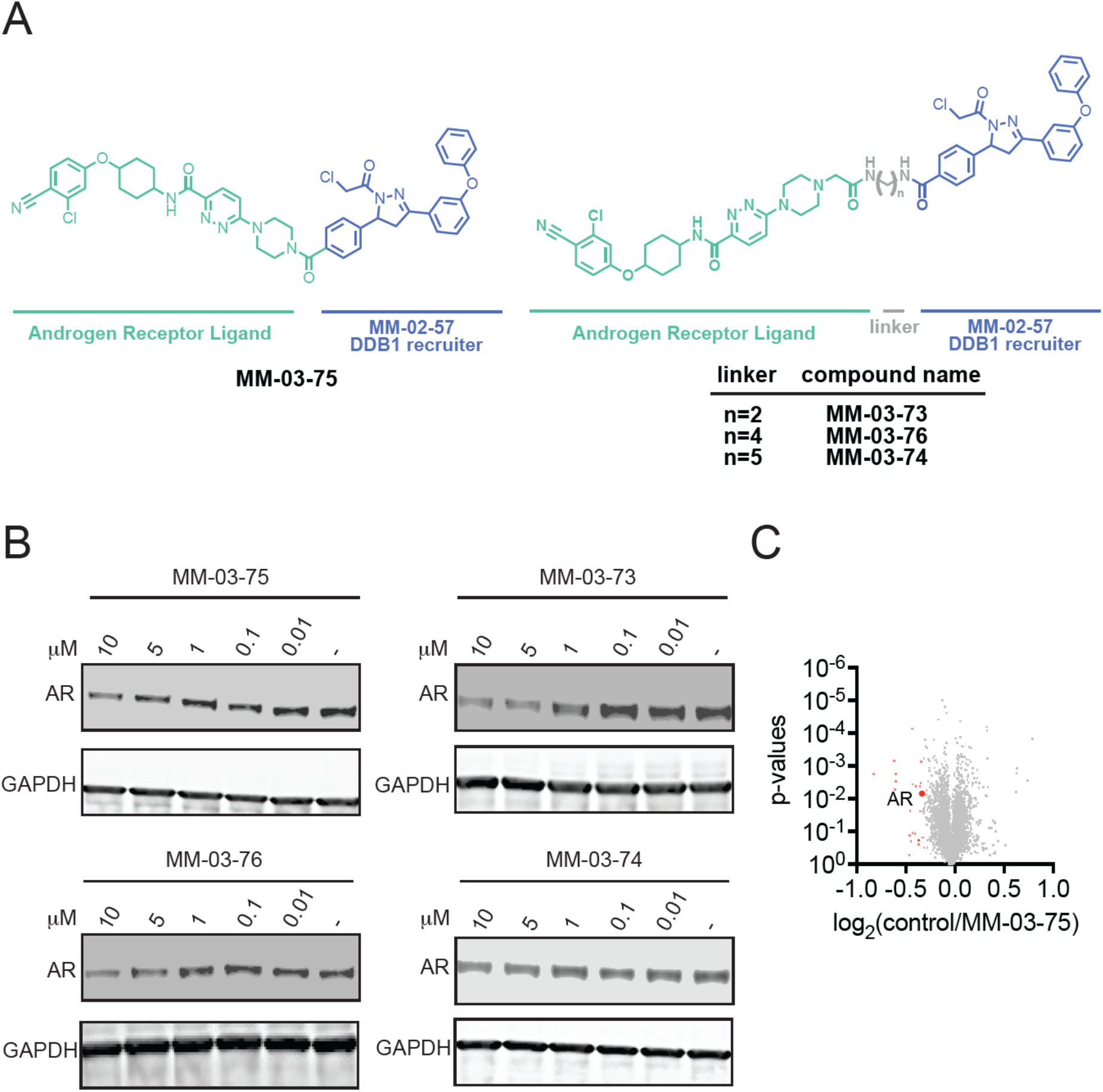
DDB1-based androgen receptor PROTACs. **(A)** Structures of MM-02-057-based AR PROTACs with different linker lengths. **(B)** AR levels in LNCaP prostate cancer cells. LNCaP cells were treated with DMSO vehicle or PROTACs for 24 h after which AR and loading control GAPDH levels were assessed by Western blotting. **(C)** TMT-based quantitative proteomic profiling of MM-03-75 in LNCaP cells. LNCaP cells were treated with MM-03-75 (10 μM) for 24 h. Red points denote statistically significant downregulated proteins including AR. Blots and proteomic data are representative of n=3 biologically independent replicates per group.

## Conclusions

Our study revealed that covalent DDB1 recruiters can be utilized in heterobifunctional PROTACs to degrade neo-substrates including BRD4 and AR. We demonstrated that our covalent handle MM-02-57 robustly engaged DDB1 C173 in cells with a relatively high degree of selectivity. We showed that the degradation observed was driven through DDB1, NEDDylation, and proteasome-mediated degradation. However, exploiting DDB1, at least through targeting C173, may be less permissive in substrate scope compared to existing CRBN and VHL recruiters given that we were only able to observe degradation of the short BRD4 isoform and we did not observe complete loss of AR. Interestingly, as was observed with the DDB1-based molecular glue degrader CR8, based on our reconstitution studies, our PROTACs showed enhanced BRD4 ubiquitination in the presence of the CUL4A complex without the necessity for a substrate adaptor protein. Targeted degradation may potentially be improved through further medicinal chemistry efforts to optimize the potency of our DDB1 recruiter. Nonetheless, our study revealed that core components of the ubiquitin-proteasome machinery that are essential for cell viability such as the CUL4A adaptor protein DDB1 can be covalently targeted for recruitment in heterobifunctional PROTAC applications and opens up the possibility for recruitment of other Cullin family adapter proteins for degradation of neo-substrates.

## Acknowledgement

We thank the members of the Nomura Research Group and Novartis Institutes for BioMedical Research for critical reading of the manuscript. This work was supported by Novartis Institutes for BioMedical Research and the Novartis-Berkeley Translational Chemical Biology Institute (NB-TCBI) for all listed authors. This work was also supported by the Nomura Research Group and the Mark Foundation for Cancer Research ASPIRE Award for DKN, NF. This work was also supported by grants from the National Institutes of Health (R01CA240981 and R35CA263814 for DKN) and the National Science Foundation Molecular Foundations for Biotechnology Award (2127788). We also thank Drs. Hasan Celik, Alicia Lund, and UC Berkeley’s NMR facility in the College of Chemistry (CoC-NMR) for spectroscopic assistance. Instruments in the College of Chemistry NMR facility are supported in part by NIH S10OD024998.

## Author Contributions

MM, DKN conceived of the project idea, designed experiments, performed experiments, analyzed and interpreted the data, and wrote the paper. MM, SC, AP, DKN performed experiments, analyzed and interpreted data, and provided intellectual contributions. BCL generated protein reagents.

## Methods

### Gel-Based ABPP

DDB1 (0.05 μg/25 μL in phosphate-buffered saline (PBS)) was treated with either DMSO vehicle or covalent ligand at 37 °C for 30 min and subsequently treated with 0.1 μM IA-rhodamine (Setareh Biotech) for 1 h at RT in the dark. The reaction was stopped by the addition of 4 × reducing Laemmli SDS sample loading buffer (Alfa Aesar). After boiling at 95 °C for 5 min, the samples were separated on precast 4–20% Criterion TGX gels (Bio-Rad). Probe-labeled proteins were analyzed by in-gel fluorescence using a ChemiDoc MP (Bio-Rad).

### Cell Culture

HEK293T cells were obtained from the UC Berkeley Cell Culture Facility and were cultured in Dulbecco’s modified Eagle medium (DMEM) containing 10% (v/v) fetal bovine serum (FBS) and maintained at 37 °C with 5% CO_2_. MDA-MB-231 cells were obtained from the UC Berkeley Cell Culture Facility and were cultured in Dulbecco’s modified Eagle medium (DMEM) containing 10% (v/v) FBS and maintained at 37 °C with 5% CO_2_. LNCaP cells were obtained from the UC Berkeley Cell Culture Facility and were cultured in RPMI-1640 medium containing 10% (v/v) FBS and maintained at 37 °C with 5% CO_2_. Unless otherwise specified, all cell culture materials were purchased from Gibco. It is not known whether HEK293T cells are from male or female origin.

### Preparation of Cell Lysates

For western blot analysis, cells were washed once with cold PBS before they were lysed in RIPA buffer (supplemented with protease inhibitor cocktail, ThermoFisher, A32963) for 30 minutes on ice. The plates were scraped and the lysate was clarified by centrifugation (6500*g*, 5 min, 4 °C). For all other experiments, cells were resuspended in PBS (supplemented with protease inhibitor cocktail ThermoFisher, A32963) and sonicated. Cells were clarified by centrifugation (6500*g*, 5 min, 4 °C), and the lysate was transferred to new low-adhesion microcentrifuge tubes. Proteome concentrations were determined using the Pierce BCA assay kit (Thermo Scientific, 23225), and the lysate was diluted to appropriate working concentrations.

### Western Blotting

Proteins were resolved by sodium dodecyl sulfate poly(acrylamide) gel electrophoresis (SDS/PAGE) and transferred to nitrocellulose membranes using the Trans-Blot Turbo transfer system (Bio-Rad). Membranes were blocked with 5% BSA in Tris-buffered saline containing Tween 20 (TBST) solution for overnight at 4 °C and then washed three times with TBST. The blot was incubated with primary antibody diluted in 5% BSA in TBST for two hours at room temperature. After three washes with TBST, the membranes were incubated in the dark with IR680-or IR800-conjugated secondary antibodies at 1:10,000 dilution in 5% BSA in TBST for 1 h at RT. After three additional washes with TBST, blots were visualized using an Odyssey Li-Cor fluorescent scanner. The membranes were stripped using ReBlot Plus Strong Antibody Stripping Solution (EMD Millipore) when additional primary antibody incubations were performed. Antibodies used in this study were GAPDH (Cell Signaling Technology, 14C10 or Proteintech, 60004-1-IG-150L), BRD4 (Abcam, ab128874), DDB1 (ABCAM, ab124672), androgen receptor (Cell Signaling Technology, 5153S), and Anti-DDDDK tag (Abcam, ab205606).

### Expression and Purification of DDB1

Human DDB1 (residues 1-1140) tagged with N-terminal His-TEV was synthesized with codons optimized for Spodoptera frugiperda expression. Bacmid DNA encoding N-terminal His-TEV tagged DDB1 was expressed in ESF921 cultured Sf21 cells. Frozen cells were lysed by homogenization at pH 7.5. The soluble fraction was purified with histidine-affinity and His-TEV tag removed with 100U/mg TEV protease. Protein was polished with size-exclusion chromatography. Protein was concentrated to ∼20 mg/mL in 20 mM HEPES, pH 7.0, 150mM NaCl, 2mM TCEP.

### IKAROS In Vitro Ubiquitination Assay

DDB1 complex with CRBN (0.6 μL, 14 μM, Boston Biochem. Inc., E3-500-025) was diluted in TBS (24.4 μL) and incubated with 0.5 μL of DMSO vehicle or MM-02-57 (50 μM final concentration) for 30 min at 37 °C. Subsequently, a 25 uL mixture of CUL4A/NEDD8/RBX1 (0.4 uL, 9 μL, Boston Biochem. Inc., E3-441-025), UBE1 (0.4 μL, 5 μM, Boston Biochem. Inc., E-305-025), UBE2D1 (5.2 μL, 3.6 μM, Boston Biochem. Inc., E2-616-100), recombinant IKAROS (3 μL, 8 μM, ABCAM, ab169877-5), (+/-)-thalidomide (1 μL, 17.5 μM, Sigma-Aldrich, T144-100MG), FLAG-ubiquitin (0.5 μL, 20 mg/mL, R&D Systems, U12001M), MgCl_2_ (1.25 μL, 200 mM), DTT (1.25 μL, 200 mM), and ATP (0.3 μL μL, 216 mM) was added to achieve a total final volume of 50 μL. The mixture was incubated at 37 °C for 1.5 h. Then, 20 μL of Laemmli SDS sample loading buffer (Alfa Aesar) was added to quench the reaction and proteins were analyzed by Western Blot. All dilutions were made using 50 mM TBS.

### DDB1 In Vitro Ubiquitination Assay

DDB1 (3.3 uL, 8 μM) was mixed with CUL4A/NEDD8/RBX1 (0.2 uL, 9 μM, Boston Biochem. Inc., E3-441-025), UBE1 (0.2 μL, 5 μM, Boston Biochem. Inc., E-305-025), UBE2D1 (2.6 μL, 3.6 μM, Boston Biochem. Inc., E2-616-100), His10-FLAG-BRD4 (0.2 μL, 7 μM, R&D Systems, SP600100), ubiquitin (10 μL, 0.5 mg/mL, ABCAM. Inc., ab269109), MgCl_2_ (1.25 μL, 200 mM), DTT (1.25 μL, 200 mM), and ATP (0.3 μL μL, 216 mM), and DMSO vehicle or MM-02-08 (50 μM final concentration). 50 μM TBS was added to achieve a total final volume of 50 μL. The mixture was incubated at 37 °C for 1.5 h. Then, 20 μL of Laemmli SDS sample loading buffer (Alfa Aesar) was added to quench the reaction and proteins were analyzed by Western Blot.

### Mapping of MM-02-57 Site of Modification on DDB1/CRBN by LC-MS/MS

DDB1/CRBN (40 μg, Boston Biochem. Inc., E3-500-025) was diluted in PBS (100 uL) and pre-incubated with MM-02-57 (50 μM final concentration) for 30 min at 37 °C. The protein was precipitated by the addition of 25 μL of TCA (100% w/v) and incubation at −80 °C overnight. The sample was then spun at 20,000*g* for 10 min at 4 °C. The supernatant was carefully removed, and the sample was washed three times with 200 μL of ice-cold 0.01 M HCl/90% acetone solution, with spinning at 20,000 for 5 min at 4 °C between washes. The sample was then resuspended in 30 μL of 8M urea in PBS and 30 μL of ProteaseMax surfactant (20 μg/mL in 100 mM ammonium bicarbonate, Promega, V2071) with vortexing. Ammonium bicarbonate (40 μL, 100 mM) was then added for a final volume of 100 μL. The sample was reduced with 10 μL of TCEP (10 mM final concentration) for 30 min at 60 °C and alkylated with 10 μL of iodoacetamide (12.5 mM final concentration) for 30 min at 37 °C. The sample was then diluted with 120 μL of PBS before 1.2 μL of ProteaseMax surfactant (0.1 mg/mL in 100 mM ammonium bicarbonate, Promega, V2071) and sequencing grade trypsin (10 μL, 0.5 mg/mL in 50 mM ammonium bicarbonate, Promega, V5111) were added for overnight incubation at 37 °C. The next day, the sample was acidified with formic acid (5% final concentration) and fractionated using high pH reversed-phase peptide fractionation kits (ThermoFisher, 84868) according to Vinogradova and coworkers (Cell, **2020**, 182, 1009-1026).

### Pulldown of DDB1 from HEK293T Cells with an MM-02-54 Probe

HEK293T cells were treated at 70% confluency with DMSO MM-02-54 (75 uM) for 2 hr. Cells were harvested, lysed via sonication, and the resulting lysate normalized to 6 mg/mL per sample. 25 uL of lysate was removed for Western Blot analysis of the input. 500 μL of each lysate sample was incubated for 1 h at RT with 10 μL of 10 mM biotin picolyl azide (in DMSO) (Sigma Aldrich 900912), 10 μL of 50 mM TCEP (in H_2_O), 30 μL of TBTA ligand (0.9 mg/mL in 1:4 DMSO/*t*BuOH), and 10 μL of 50 mM CuSO_4_. Proteins were precipitated, washed three times with cold MeOH, resolubilized in 200 μL of 1.2% SDS/PBS (w/v), and heated for 5 min at 90 °C. The sample was centrifuged (5 min, 10,000*g*) to remove any insoluble components. The supernatant was transferred into a tube containing prewashed high-capacity streptavidin beads (50 μL, Thermo Scientific, 20359) in PBS (1 mL). The proteins were incubated with the beads at 4 °C overnight on a rotator. The following day, the samples were warmed to RT and washed with 0.2% SDS and further washed three times with 500 μL of PBS and three times with 500 μL of H_2_O to remove nonprobe-labeled proteins. The washed beads were resuspended in 30 μL of Laemmli SDS sample loading buffer (Alfa Aesar), heated to 95 °C for 5 min, and analyzed by Western Blot.

### DDB1 Lentiviral Knockdown Studies

In separate 15 mL conicals, 1 μg of expression clone cDNA (Origene NM_001923) or control cDNA (Origene PS100093) was mixed with packaging plasmids MD2G (1 μg, <u>Addgene</u> 12,259) and PSPAX2 (1 μg, Addgene 12,260) in 600 μL per plate OPTIMEM and Lipofectamine 2000 transfection reagent (Invitrogen 11668027) was incubated with an equal volume of OPTIMEM (1:30 v/v) for 5 min prior to tubes being combined and incubated for 40 min at room temperature. The DNA-Lipofectamine mix was diluted with 8 mL of DMEM and added to HEK293T cells at 40% confluency in 10 cm plates. The next day, media was replaced with 6 mL fresh DMEM for 24 h.

For each control or knock-down clone, media was removed from HEK293T cells, filtered through a 0.45-micron syringe filter, mixed with 10 μL polybrene transfection reagent, and added to new HEK293T cells at 50% confluency. The original HEK293T media was replaced with 6 mL fresh DMEM for 24 h and the infection process was repeated. 24 h after the second infection, the new HEK293T infection media was removed, and cells were seeded for Western blot analysis.

### IsoDTB-ABPP Cysteine Chemoproteomic Profiling of MM-02-57

IsoDTB-ABPP cysteine chemoproteomic profiling was performed as described previously ^21^. HEK293T cells were treated with either MM-02-57 (50 μM) or DMSO for 4 h before cell collection and lysis. The proteome concentrations were determined using BCA assay and adjusted to 2 mg/mL. For each biological replicate, 2 aliquots of 1 mL of 2 mg/mL were used (i.e. 4 mg per condition). Each aliquot was treated with 20 μL of IA-alkyne (26.6 mg/mL in DMSO, 200 μM final concentration) for 1 h at RT. Two master mixes of the click reagents were prepared in the meanwhile, each containing 1020 μL TBTA (0.9 mg/mL in 4:1 tBuOH/DMSO), 330 μL CuSO4 (12.5 mg/mL in H2O), 330 μL TCEP (14.0 mg/mL in H2O) and 160 μL of either heavy or light isoDTB tags (4 mg in DMSO, Click Chemistry Tools, 1565). The samples were then treated with 120 μL of the heavy (DMSO treated) or light (compound treated) master mix for 1 h at RT. After incubation, one light and one heavy-labeled samples were combined and acetone-precipitated overnight at -20 °C. The samples were then centrifuged at 3,500 rpm for 10 min, acetone was removed and they were resuspended in cold MeOH by sonication. They were centrifuged at 3,500 rpm for 10 min and MeOH was removed (repeated 3× in total). The pellets were dissolved in 600 μL urea (8M in 0.1 M TEAB) by sonication and the urea concentration was then adjusted to 2M by adding 1800 μL of TEAB (0.1 M). Two tubes containing solubilized proteins were combined, further diluted with 2400 μL 0.2% NP40 in PBS and bound to high-capacity streptavidin agarose beads (200 μL/sample, ThermoFisher, 20357) for 1 h at RT with mixing. The beads were then centrifuged for 1 min at 1,000 g, the supernatant was removed and the beads were washed 3 times with 0.1% NP40 in PBS, 3 times with PBS and 3 times with H2O. They were then resuspended in 8M urea (600 μL in 0.1 M TEAB) and treated with DTT (30 μL, 31 mg/mL in H2O) for 45 min at 37 °C. They were then reacted with iodoacetamide (30 μL, 74 mg/mL in H2O) for 30 min at RT, followed by DTT (30 μL, 31 mg/mL in H2O) for 30 min at RT. The samples were diluted with 1800 μL TEAB (0.1 M), centrifuged for 1 min at 1,000 g and the supernatant was removed. The beads were resuspended in 400 μL urea (2M in 0.1 M TEAB), trypsin (8 μL, 0.5 mg/mL) was added and incubated for 20 h at 37 °C. The samples were then diluted with 800 μL 0.1% NP40 in PBS and the beads were washed 3 times with 0.1% NP40 in PBS, 3 times with PBS and 3 times with H2O. Peptides were then eluted by incubating beads with 0.1% formic acid in 50% acetonitrile (500 μL) for 10 min before eluting via centrifugation. The elution step was repeated 2 more times and the three elution fractions were combined. The samples were then dried by using a vacuum concentrator at 30 °C, resuspended in 300 μL in 5% acetonitrile and 0.1% formic acid in water, and fractionated using high pH reversed-phase peptide fractionation kits (ThermoFisher, 84868) according to Vinogradova and coworkers ^28^.

### IsoDTB-ABPP Mass Spectrometry Analysis

Mass spectrometry analysis was performed on an Orbitrap Eclipse Tribrid Mass Spectrometer with a High Field Asymmetric Waveform Ion Mobility (FAIMS Pro) Interface (Thermo Scientific) with an UltiMate 3000 Nano Flow Rapid Separation LCnano System (Thermo Scientific). Off-line fractionated samples (5 μl aliquot of 15 μl sample) were injected via an autosampler (Thermo Scientific) onto a 5 μl sample loop which was subsequently eluted onto an Acclaim PepMap 100 C18 HPLC column (75 μm x 50 cm, nanoViper). Peptides were separated at a flow rate of 0.3 μl/min using the following gradient: 2 % buffer B (100 % acetonitrile with 0.1 % formic acid) in buffer A (95:5 water:acetonitrile, 0.1 % formic acid) for 5 min, followed by a gradient from 2 to 40 % buffer B from 5 to 159 min, 40 to 95 % buffer B from 159 to 160 minutes, holding at 95 % B from 160-179 min, 95 % to 2 % buffer B from 179 to 180 min, and then 2 % buffer B from 180 to 200 min. Voltage applied to the nano-LC electrospray ionization source was 2.1 kV. Data was acquired through an MS1 master scan (Orbitrap analysis, resolution 120,000, 400-1800 m/z, RF lens 30 %, heated capillary temperature 250 °C) with dynamic exclusion enabled (repeat count 1, duration 60 s). Data-dependent data acquisition comprised a full MS1 scan followed by sequential MS2 scans based on 2 s cycle times. FAIMS compensation voltages (CV) of -35, -45, and -55 were applied. MS2 analysis consisted of quadrupole isolation window of 0.7 m/z of precursor ion followed by higher energy collision dissociation (HCD) energy of 38 % with a orbitrap resolution of 50,000. Data were extracted in the form of MS1 and MS2 files using Raw Converter (Scripps Research Institute) and searched against the Uniprot human database using ProLuCID search methodology in IP2 v.3-1 v.5 (Integrated Proteomics Applications, Inc.). Cysteine residues were searched with a static modification for carboxyaminomethylation (+57.02146) and up to two differential modifications for methionine oxidation and either the light or heavy isoDTB tags (+561.33872 or +567.34621, respectively). Peptides were required to be fully tryptic peptides and to contain the TEV modification. ProLUCID data were filtered through DTASelect to achieve a peptide false-positive rate below 5%. Only those probe-modified peptides that were evident across two out of three biological replicates were interpreted for their isotopic light to heavy ratios. Light versus heavy isotopic probe-modified peptide ratios are calculated by taking the mean of the ratios of each replicate paired light versus heavy precursor abundance for all peptide-spectral matches associated with a peptide. The paired abundances were also used to calculate a paired sample *t*-test *P* value in an effort to estimate constancy in paired abundances and significance in change between treatment and control. *P* values were corrected using the Benjamini–Hochberg method.

### TMT-based quantitative proteomic profiling

LNCaP cells were treated at 70% confluency with either DMSO vehicle or MM-03-75 (10 μM) for 24 h and lysate was prepared as described above. Briefly, 25-100 μg protein from each sample was reduced, alkylated and tryptically digested overnight. Individual samples were then labeled with isobaric tags using commercially available TMTsixplex (ThermoFisher, 90061) kits, in accordance with the manufacturer’s protocols. Tagged samples (20 μg per sample) were combined, dried with SpeedVac, with resuspended in 300 μL in 5% acetonitrile and 0.1% formic acid in water, and fractionated using high pH reversed-phase peptide fractionation kits (ThermoFisher, 84868) according to Vinogradova and coworkers ^28^. Fractions were dried with SpeedVac, resuspended with 50 μL 0.1% FA in H2O, and analyzed by LC-MS/MS as described below.

Mass spectrometry analysis of resulting TMT peptides were performed as described above. Trypsin cleavage specificity (cleavage at K, R except if followed by P) allowed for up to 2 missed cleavages.

Carbamidomethylation of cysteine was set as a fixed modification, methionine oxidation, and TMT-modification of *N*-termini and lysine residues were set as variable modifications. Reporter ion ratio calculations were performed using summed abundances with most confident centroid selected from 20 ppm window. Only peptide-to-spectrum matches that are unique assignments to a given identified protein within the total dataset are considered for protein quantitation. High confidence protein identifications were reported with a <1% false discovery rate (FDR) cut-off. Differential abundance significance was estimated using ANOVA with Benjamini-Hochberg correction to determine p-values.

