## Supporting Information for "Targeted Protein Degradation through Recruitment of the CUL4A Complex Adaptor Protein DDB1"

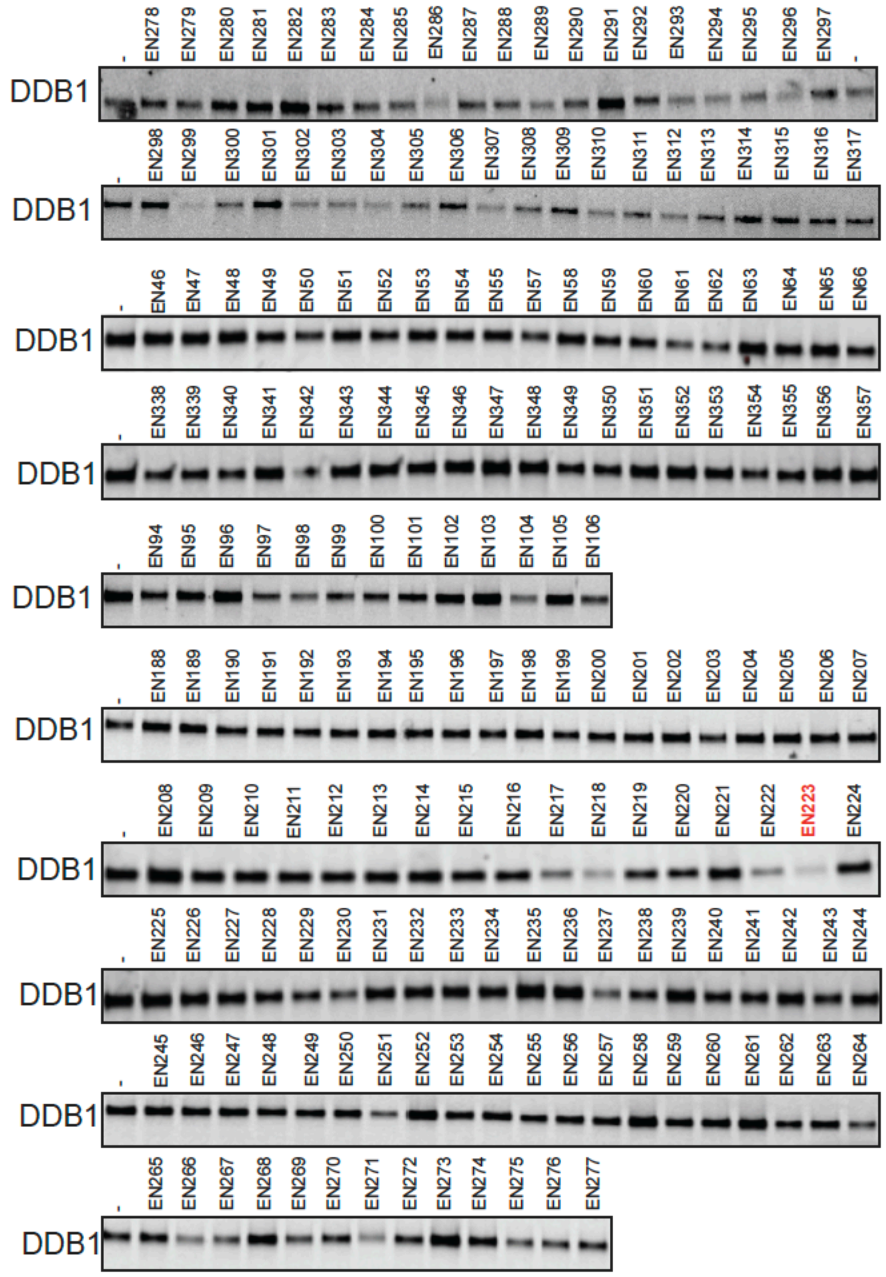


**Figure S1. Covalent ligand screening against DDB1.** Gel-based ABPP screening of covalent ligands against IA-rhodamine labeling of pure DDB1 protein. DDB1 was pre-incubated with DMSO or the covalent ligand (50 μM) for 30 min prior to IA-rhodamine labeling (100 nM) for 60 min after which proteins were resolved by SDS/PAGE and IA-rhodamine labeling was visualized by in-gel fluorescence and loading was assessed by silver staining.

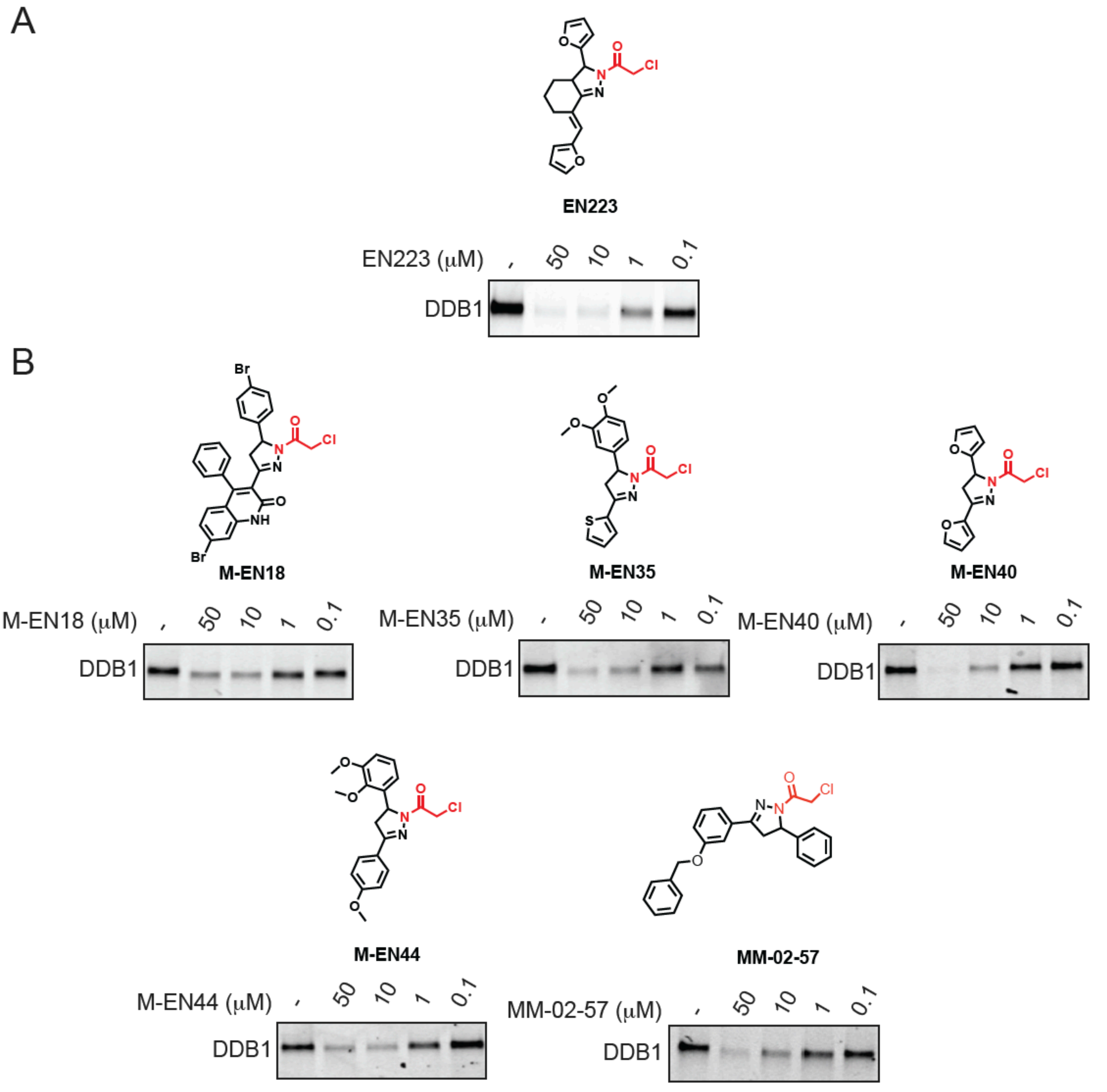


**Figure S2. Further testing of DDB1 hit ligands and analogs. (A, B)** Gel-based ABPP screening of covalent ligands against IA-rhodamine labeling of pure DDB1 protein. DDB1 was pre-incubated with DMSO or the covalent ligand for 30 min prior to IA-rhodamine labeling (100 nM) for 60 min after which proteins were resolved by SDS/PAGE and IA-rhodamine labeling was visualized by in-gel fluorescence and loading was assessed by silver staining. Gels are representative of n=3 biologically independent replicates/group.

**Supporting Tables.**

**Table S1.** Structures and names of cysteine-reactive covalent ligand screened against DDB1.

**Table S2.** IsoDTB-ABPP analysis of MM-02-57 in HEK293T cells. HEK293T cells were treated with DMSO vehicle or MM-02-57 (50 μM) for 4 hours after which resulting control and treated lysates were labeled with an alkyne-functionalized iodoacetamide probe (IA-alkyne) for 1 h. Probe-labeled proteins were subjected to CuAAC to append either an isotopically light (for control) or heavy (for treated) azide-functionalized desthiobiotin handle, after which probe-labeled proteins were avidin-enriched, digested with trypsin, and probe-modified peptides were eluted and analyzed by LC-MS/MS. Control/treated probe-modified peptides were quantified.

**Synthetic Methods and Characterization**

Reagents were purchased at the highest commercial quality and used without further purification, unless otherwise stated. Room temperature is defined as between 21-25 °C. Reactions were stirred magnetically and monitored by thin layer chromatography (TLC) using TLC plates pre- coated with silica gel 60 F254 on aluminium (Merck KGaA). Detection was by UV (254 nm and 365 nm) or chemical stain (ninhydrin, iodine). Solvents were removed *in vacuo* using a Buchi R-300 Rotavapor (equipped with an I-300 Pro Interface, B-300 Base Heating Bath, Welch 2037B-01 DryFast pump, and VWR AD15R-40-V11B Circulating Bath) or a Biotage V-10 Touch. Solvents for silica gel chromatography were used as supplied by Sigma- Aldrich. Automated flash chromatography was performed on a Biotage Isolera instrument, equipped with a UV detector. Chromatograms were recorded at 254 and 280 nm. High-resolution mass spectra (HRMS) obtained on a Q Exactive Plus mass spectrometer (ThermoFisher Scientific). Nuclear Magnetic Resonance (NMR) spectra were recorded on BRUKER AV spectrometers operating at 600 MHz or 400 MHz for $^{1}$H NMR and 126 or 151 MHz $^{13}$C NMR. Measurements were carried out at ambient temperature. Chemical shifts (δ) are reported in ppm with the residual solvent signal as internal standard (chloroform at 7.26 and 77.2 ppm for $^{1}$H NMR and $^{13}$C NMR, respectively, acetone at 2.05 and 29.8, respectively). The multiplicity of each signal is indicated as s = singlet, d = doublet, t = triplet, q = quartet, quin = quintet, m = multiplet (i.e. complex peak obtained due to overlap), app = apparent or a combination of these. Coupling constants (J) are reported in Hertz (Hz). $^{13}$C NMR spectra were recorded with broadband $^{1}$H decoupling.

**1-(3-(benzyloxy)phenyl)ethan-1-one (1)**

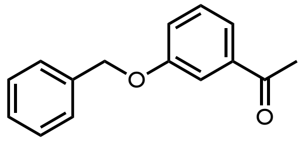

3-hydroxyacetophenone (500 mg, 3.7 mmol) was dissolved in ethanol (70 mL). The solution was heated to 70°C. Benzyl bromide and potassium carbonate in ethanol (5 mL) were added to the reaction mixture. The reaction was stirred overnight. The mixture was concentrated and then the product was purified by column chromatography (gradient elution 0-50% ethyl acetate in hexanes). The target product was a clear oil (760 mg, 3.4 mmol, 92% yield).
$^{1}$H NMR (400 MHz, $CDCl_3$) δ 7.56 – 7.46 (m, 2H), 7.41 (dd, *J* = 2.7, 1.6 Hz, 2H), 7.32 (t, *J* = 7.9 Hz, 2H), 7.03 (ddd, *J* = 8.1, 2.6, 1.0 Hz, 2H), 5.09 (s, 2H), 1.54 (s, 3H).

**(*E*)-1-(3-(benzyloxy)phenyl)-3-phenylprop-2-en-1-one (2)**

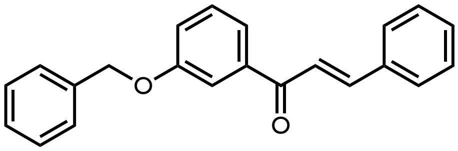

To a solution of **1** (700 mg, 3.1 mmol) in ethanol (20 mL), benzaldehyde (360 mg, 3.0 mmol) and potassium hydroxide, (260.4 mg, 4.6 mmol) each dissolved in ethanol, were added. The reaction was stirred at room temperature overnight. The product precipitated as a white solid and was isolated via filtration. The product was redissolved in DCM (5 mL) and purified by column chromatography (gradient elution 0-50% ethyl acetate in hexanes). The target product was a white solid (440 mg, 1.4 mmol, 45%).
$^{1}$H NMR (400 MHz, $CDCl_3$) δ 7.81 (d, *J* = 15.8 Hz, 1H), 7.69 – 7.59 (m, 4H), 7.54 – 7.29 (m, 10H), 7.21 (ddd, *J* = 8.2, 2.6, 1.1 Hz, 1H), 5.15 (s, 2H).

***tert*-butyl 3-(3-(benzyloxy)phenyl)-5-phenyl-4,5-dihydro-1*H*-pyrazole-1-carboxylate (3)**

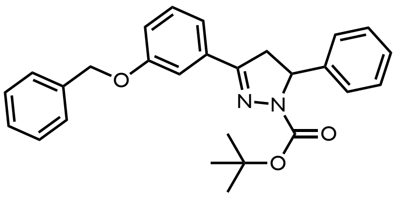

A round bottomed flask containing **2** (420 mg, 1.4 mmol) was purged with $N_2$ before anhydrous acetonitrile (15 mL) was added. The solution was heated to 60°C and stirred for 5 min to dissolve **2**. Once there was a uniform solution, tert-butyl carbazate (212 mg, 1.6 mmol) and the catalyst triazabicyclodecene (37 mg, 0.27 mmol) were each added in anhydrous acetonitrile (5 mL). The reaction was heated overnight at 60°C under $N_2$. The next day the reaction was concentrated and

purified by column chromatography (gradient elution 0-50% ethyl acetate in hexanes). The target product was a white solid (244 mg, 0.57 mmol, 41% yield).
$^{1}H$ NMR (400 MHz, $CDCl_3$) δ 7.52 (s, 1H), 7.48 – 7.20 (m, 11H), 7.01 (ddd, *J* = 7.6, 2.6, 1.8 Hz, 1H), 5.40 – 5.31 (m, 1H), 3.74 (dd, *J* = 17.5, 12.1 Hz, 1H), 3.16 (dd, *J* = 17.5, 5.4 Hz, 1H), 1.31 (s, 9H).

**1-(3-(3-(benzyloxy)phenyl)-5-phenyl-4,5-dihydro-1*H*-pyrazol-1-yl)-2-chloroethan-1-one (4)**

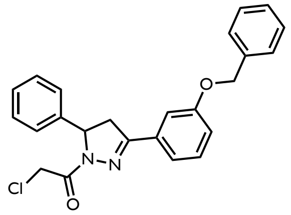


To **3** (244 mg, 0.57 mmol) in dichloromethane (2 mL) was added TFA (1 mL). The reaction mixture was stirred for 1 hr at room temperature before it was concentrated *in vacuo*. The reaction mixture was redissolved in dichloromethane (2 mL) and cooled to -10°C. Chloroacetyl chloride was added dropwise (68.1 μL, 0.86 mmol) before triethylamine was added (238 μL, 1.7 mmol). The reaction was brought to room temperature and then stirred for 30 mins. The reaction was concentrated and then purified by column chromatography (gradient elution 0-50% ethyl acetate in hexanes). The target product was a yellow solid (116.3 mg, 0.29 mmol, 51% yield).
$^{1}H$ NMR (600 MHz, Acetone) δ 7.54 – 7.22 (m, 12H), 7.14 (ddd, *J* = 8.0, 2.6, 1.2 Hz, 1H), 5.65 – 5.57 (m, 1H), 5.19 (s, 2H), 4.78 – 4.53 (m, 2H), 3.93 (dd, *J* = 18.0, 11.8 Hz, 1H), 3.29 – 3.18 (m, 1H). $^{13}C$ NMR (151 MHz, Acetone) δ 163.28, 159.11, 155.19, 142.00, 137.24, 132.65, 129.88, 128.69, 128.46, 127.88, 127.69, 127.46, 125.69, 119.58, 117.16, 113.03, 69.77, 60.51, 42.22, 42.12; HRMS (ES+) calcd for $C_{24}H_{21}ClN_2O_2Na$ $[M+Na]^+$ 427.1292 found 427.1175

**(*E*)-1-(3-(benzyloxy)phenyl)-3-(4-ethynylphenyl)prop-2-en-1-one (5)**

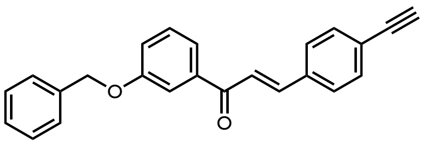


To a solution of **1** (300 mg, 1.3 mmol) in ethanol (10 mL), ethynyl-benzaldehyde (190 mg, 1.5 mmol) and potassium hydroxide, (112 mg, 2.0 mmol) each dissolved in ethanol, were added. The reaction was stirred at room temperature overnight. The product precipitated as a white solid and was isolated via filtration. The product was redissolved in DCM (5 mL) and purified by column chromatography (gradient elution 0-50% ethyl acetate in hexanes). The target product was a white solid (208 mg, 0.62 mmol, 48%).
$^{1}H$ NMR (400 MHz, $CDCl_3$) δ 7.81 (d, *J* = 15.7 Hz, 1H), 7.70 – 7.35 (m, 14H), 7.26 (ddd, *J* = 8.2, 2.5, 1.1 Hz, 1H), 5.19 (s, 2H), 3.25 (s, 1H).

***tert*-butyl 3-(3-(benzyloxy)phenyl)-5-(4-(methoxycarbonyl)phenyl)-4,5-dihydro-1*H*-pyrazole-1-carboxylate (6)**

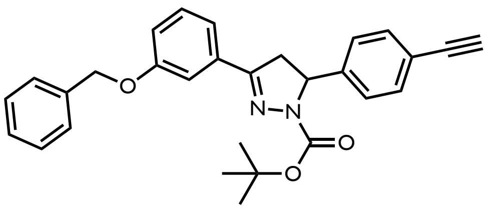


A round bottomed-flask containing **5** (208.3 mg, 0.62 mmol) was purged with $N_2$ before anhydrous acetonitrile (10 mL) was added. The solution was heated to 60°C and stirred for 5 min to dissolve **6**. Once there was a uniform solution tert-butyl carbazate (97.7 mg, 0.74 mmol) and the catalyst triazabicyclodecene (17.1 mg, 0.12 mmol) were each added in anhydrous acetonitrile (3 mL). The reaction was heated overnight at 60°C under $N_2$. The next day the reaction was concentrated and purified by column chromatography (gradient elution 0-50% ethyl acetate in hexanes). The target product was a white solid (126.5 mg, 0.28 mmol, 45% yield).
$^{1}H$ NMR (400 MHz, $CDCl_3$) δ 7.60 – 7.32 (m, 11H), 7.25 (d, *J* = 8.2 Hz, 2H), 7.10 – 7.03 (m, 1H), 5.34 (s, 0H), 5.15 (s, 2H), 3.79 (dd, *J* = 17.5, 12.3 Hz, 1H), 3.12 (s, 1H), 1.37 (s, 12H).

**1-(3-(3-(benzyloxy)phenyl)-5-(4-ethynylphenyl)-4,5-dihydro-1*H*-pyrazol-1-yl)-2-chloroethan-1-one (7)**

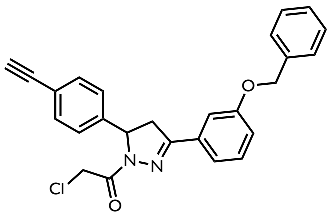

To **6** (116 mg, 0.26 mmol) in dichloromethane (2 mL) was added TFA (1 mL). The reaction mixture was stirred for 1 hr at room temperature before it was concentrated *in vacuo*. The reaction mixture was redissolved in dichloromethane (2 mL) and cooled to -10$^o$C. Chloroacetyl chloride was added dropwise (30.6 $\mu$L, 0.38 mmol) before triethylamine was added (107 $\mu$L, 0.77 mmol). The reaction was brought to room temperature and then stirred for 30 mins. The reaction was concentrated and then purified by column chromatography (gradient elution 0-50% ethyl acetate in hexanes). The target product was a yellow solid (59.4 mg, 0.13 mmol, 51% yield).
$^1$H NMR (600 MHz, Acetone) δ 7.55 – 7.49 (m, 3H), 7.50 – 7.46 (m, 2H), 7.46 – 7.38 (m, 4H), 7.38 – 7.33 (m, 1H), 7.34 – 7.29 (m, 2H), 7.16 (ddd, *J* = 7.9, 2.6, 1.3 Hz, 1H), 5.69 – 5.61 (m, 1H), 5.20 (s, 2H), 4.72 (d, *J* = 13.6 Hz, 1H), 4.64 (d, *J* = 13.6 Hz, 1H), 4.01 – 3.90 (m, 1H), 3.31 – 3.22 (m, 1H).
$^{13}$C NMR (151 MHz, Acetone) δ 163.40, 159.11, 155.24, 142.73, 137.22, 132.54, 132.29, 129.90, 128.47, 127.89, 127.69, 126.06, 121.52, 119.62, 117.20, 113.08, 83.02, 78.42, 69.78, 60.28, 42.06 (d, *J* = 8.0 Hz); HRMS (ES+) calcd for $C_{26}H_{21}ClN_2O_2Na$ [M+Na]$^+$ 451.12916, found 451.11725

**methyl (*E*)-4-(3-(3-(benzyloxy)phenyl)-3-oxoprop-1-en-1-yl)benzoate (8)**

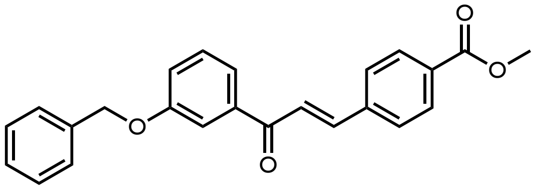

To a solution of **1** (1.5 g, 6.2 mmol) in ethanol (30 mL), methyl 4-formylbenzoate (1.0 g, 6.2 mmol) and potassium hydroxide, (518 mg, 9.2 mmol) each dissolved in ethanol, were added. The reaction was stirred at room temperature overnight. The product precipitated as a white solid and was isolated via filtration. The product was redissolved in DCM (5 mL) and purified by column chromatography (gradient elution 0-50% ethyl acetate in hexanes). The target product was a white solid (1.2 g, 3.2 mmol, 52% yield).
$^1$H NMR (500 MHz, $CDCl_3$) δ 8.13 – 8.05 (m, 2H), 7.80 (d, *J* = 15.7 Hz, 1H), 7.72 – 7.65 (m, 2H), 7.65 – 7.60 (m, 2H), 7.59 – 7.53 (m, 1H), 7.50 – 7.38 (m, 5H), 7.36 (d, *J* = 7.2 Hz, 1H), 7.23 (d, *J* = 1.9 Hz, 1H), 5.15 (s, 2H), 3.95 (s, 3H).

***tert*-butyl 3-(3-(benzyloxy)phenyl)-5-(4-ethynylphenyl)-4,5-dihydro-1*H*-pyrazole-1-carboxylate (9)**

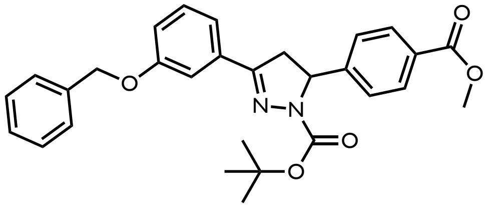

A round bottomed-flask containing **8** (1.2 g, 3.1 mmol) was purged with $N_2$ before anhydrous acetonitrile (30 mL) was added. The solution was heated to 60$^o$C and stirred for 5 min to dissolve **9**. Once there was a uniform solution tert-butyl carbazate (490.0 mg, 3.7 mmol) and the catalyst triazabicyclodecene (86.0 mg, 0.62 mmol) were each added in anhydrous acetonitrile (5 mL). The reaction was heated overnight at 60$^o$C under $N_2$. The next day the reaction was concentrated and purified by column chromatography (gradient elution 0-50% ethyl acetate in hexanes). The target product was a white solid (836 mg, 1.7 mmol, 55% yield).
$^1$H NMR (500 MHz, $CDCl_{3z}$) δ 7.92 – 7.87 (m, 2H), 7.38 (s, 1H), 7.35 – 7.11 (m, 10H), 6.91 (dtd, *J* = 8.3, 4.4, 2.5 Hz, 1H), 5.27 (dd, *J* = 12.1, 5.7 Hz, 1H), 4.98 (s, 2H), 3.79 (s, 3H), 3.65 (dd, *J* = 17.6, 12.2 Hz, 1H), 3.01 (dd, *J* = 17.4, 5.6 Hz, 1H), 1.37 – 1.03 (m, 9H).

**General Procedure A: Preparation of linkers coupled to (+)-JQ1**

*i.* To a 20 mL scintillation vial (+)-JQ1 (1 eq) was dissolved in DCM (0.05 M). A N-boc diamino alkane (1 eq.), HATU (2 eq.) and DIPEA (5 eq.) were added stepwise. The resulting reaction mixture was stirred for 20 min at room temperature. The reaction was concentrated *in vacuo* and purified via column chromatography (gradient elution, 0-10% Methanol in DCM).

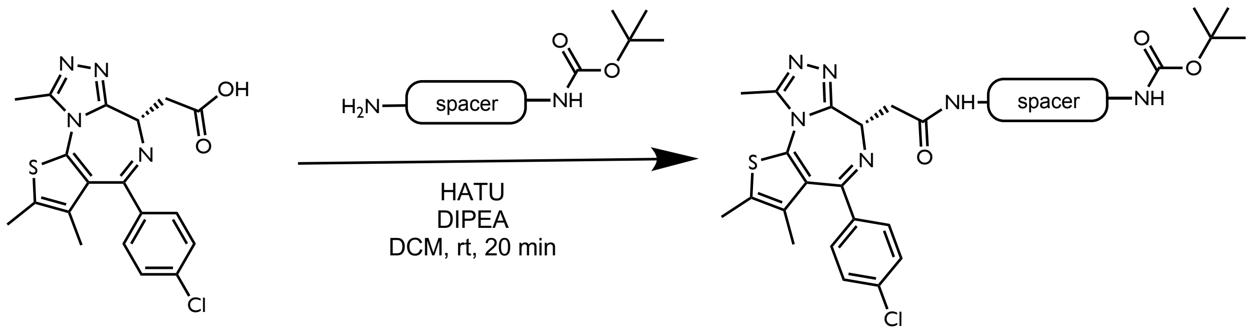


***tert*-butyl (*S*)-(2-(2-(4-(4-chlorophenyl)-2,3,9-trimethyl-6*H*-thieno[3,2-*f*][1,2,4]triazolo[4,3-*a*][1,4]diazepin-6-yl)acetamido)ethyl)carbamate (10)**

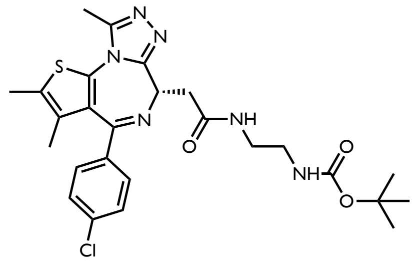

(+)-JQ1 (68 mg, 0.15 mmol) was coupled to N-Boc-ethylenediamine (27 mg, 0.17 mmol) following General Procedure A to yield **10** (92 mg, 0.17 mmol, quant. 113%) as a white solid.
$^{1}$H NMR (400 MHz, $CDCl_3$) δ 7.62 (s, 1H), 7.49 – 7.32 (m, 4H), 5.63 (s, 1H), 4.73 (t, *J* = 7.0 Hz, 1H), 3.60 (dd, *J* = 14.7, 7.4 Hz, 1H), 3.54 – 3.22 (m, 5H), 2.72 (d, *J* = 3.9 Hz, 3H), 2.44 (s, 3H), 1.71 (s, 3H), 1.57 – 1.38 (m, 9H).

***tert*-butyl (*S*)-(3-(2-(4-(4-chlorophenyl)-2,3,9-trimethyl-6*H*-thieno[3,2-*f*][1,2,4]triazolo[4,3-*a*][1,4]diazepin-6-yl)acetamido)propyl)carbamate (11)**

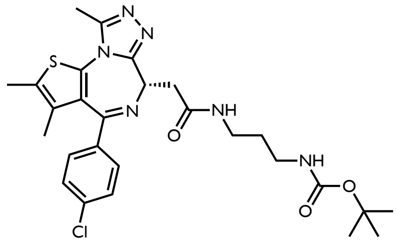

(+)-JQ1 (60 mg, 0.15 mmol) was coupled to N-Boc-1,3-propanediamine (26 mg, 0.15 mmol) following General Procedure A to yield **11** (48 mg, 0.09 mmol, 58%) as a white solid.
$^{1}$H NMR (400 MHz, $CDCl_3$) δ 7.48 – 7.32 (m, 5H), 4.71 (td, *J* = 7.0, 2.4 Hz, 1H), 3.57 (dd, *J* = 14.5, 6.9 Hz, 1H), 3.53 – 3.30 (m, 3H), 3.18 (d, *J* = 6.4 Hz, 3H), 2.72 (d, *J* = 2.4 Hz, 3H), 2.44 (d, *J* = 2.3 Hz, 3H), 1.70 (s, 3H), 1.46 (s, 11H).

***tert*-butyl (*S*)-(4-(2-(4-(4-chlorophenyl)-2,3,9-trimethyl-6*H*-thieno[3,2-*f*][1,2,4]triazolo[4,3-*a*][1,4]diazepin-6-yl)acetamido)butyl)carbamate (12)**

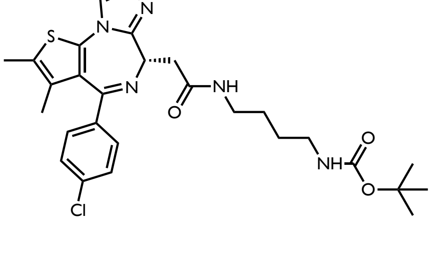

(+)-JQ1 (60 mg, 0.15 mmol) was coupled to N-Boc-1,4-butanediamine (28 mg, 0.15 mmol) following General Procedure A to yield **12** (21.8 mg, 0.04 mmol, 25%) as a white solid.
$^1$H NMR (400 MHz, $CDCl_3$) δ 7.48 – 7.33 (m, 4H), 6.87 (s, 1H), 4.80 (s, 1H), 4.67 (dd, *J* = 7.8, 6.2 Hz, 1H), 3.59 (dd, *J* = 14.4, 7.8 Hz, 1H), 3.43 – 3.28 (m, 3H), 3.15 (s, 2H), 2.71 (d, *J* = 1.5 Hz, 3H), 2.44 (s, 3H), 1.70 (s, 3H), 1.66 – 1.50 (m, 5H), 1.47 (s, 9H).

**_tert_-butyl (_S_)-(5-(2-(4-(4-chlorophenyl)-2,3,9-trimethyl-6_H_-thieno[3,2-_f_][1,2,4]triazolo[4,3-_a_][1,4]diazepin-6-yl)acetamido)pentyl)carbamate (13)**

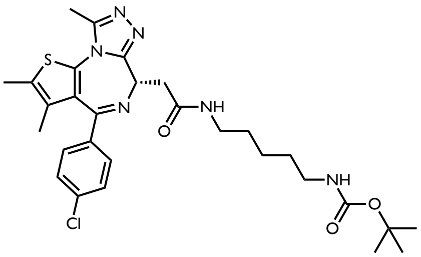

(+)-JQ1 (60 mg, 0.15 mmol) was coupled to N-Boc-1,5-pentanediamine (32 mg, 0.15 mmol) following General Procedure A to yield **13** (60.0 mg, 0.12 mmol, 80%) as a white solid.
$^1$H NMR (400 MHz, $CDCl_3$) δ 7.48 – 7.28 (m, 4H), 6.80 (t, *J* = 5.6 Hz, 1H), 4.75 (s, 1H), 4.61 (dd, *J* = 7.7, 6.0 Hz, 1H), 3.59 – 3.42 (m, 2H), 3.39 – 3.17 (m, 3H), 3.17 – 2.97 (m, 2H), 2.66 (d, *J* = 1.4 Hz, 3H), 2.38 (s, 3H), 1.65 (s, 3H), 1.53 (q, *J* = 7.4 Hz, 3H), 1.42 (d, *J* = 4.4 Hz, 9H), 1.32 (qd, *J* = 8.9, 5.9 Hz, 3H).

**_tert_-butyl (_S_)-(6-(2-(4-(4-chlorophenyl)-2,3,9-trimethyl-6_H_-thieno[3,2-_f_][1,2,4]triazolo[4,3-_a_][1,4]diazepin-6-yl)acetamido)hexyl)carbamate (14)**

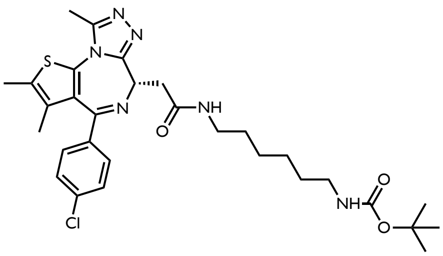

(+)-JQ1 (60 mg, 0.15 mmol) was coupled to N-Boc-1,6-hexanediamine (32 mg, 0.15 mmol) following General Procedure A to yield **14** (93.7 mg, 0.15 mmol, quant. 100%) as a white solid.
$^1$H NMR (400 MHz, $CDCl_3$) δ 7.44 (dd, *J* = 8.4, 1.9 Hz, 2H), 7.40 – 7.33 (m, 2H), 6.73 (d, *J* = 5.9 Hz, 1H), 4.82 – 4.61 (m, 2H), 3.59 (dd, *J* = 14.4, 7.8 Hz, 1H), 3.41 – 3.24 (m, 3H), 3.20 – 3.07 (m, 2H), 2.71 (d, *J* = 1.7 Hz, 3H), 2.44 (s, 2H), 1.70 (s, 3H), 1.61 – 1.41 (m, 15H), 1.35 (dd, *J* = 6.7, 3.0 Hz, 4H).

**_tert_-butyl (_S_)-(7-(2-(4-(4-chlorophenyl)-2,3,9-trimethyl-6_H_-thieno[3,2-_f_][1,2,4]triazolo[4,3-_a_][1,4]diazepin-6-yl)acetamido)heptyl)carbamate (15)**

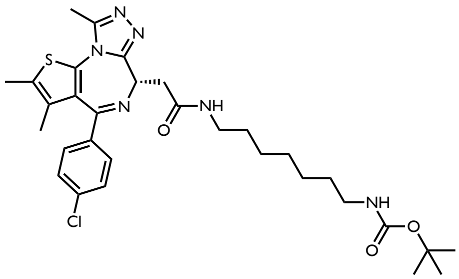

(+)-JQ1 (60 mg, 0.15 mmol) was coupled to N-Boc-1,7-heptanediamine (35 mg, 0.15 mmol) following General Procedure A to yield **15** (64 mg, 0.1 mmol, 76%) as a white solid.
$^1$H NMR (400 MHz, $CDCl_3$) δ 7.44 (d, *J* = 8.1 Hz, 2H), 7.39 – 7.28 (m, 2H), 6.68 (s, 1H), 4.70 – 4.62 (m, 2H), 3.58 (dd, *J* = 14.2, 7.7 Hz, 1H), 3.41 – 3.20 (m, 3H), 3.12 (d, *J* = 7.2 Hz, 2H), 2.70 (s, 3H), 2.43 (s, 3H), 1.70 (s, 3H), 1.59 – 1.51 (m, 2H), 1.47 (s, 9H), 1.33 (s, 7H).

**General Procedure B: Preparation of the Boc protected DDB1 recruiter linked to (+)-JQ1-linker derivatives**

*i.* **9** (1 eq) was charged to a 20 mL scintillation vial and dissolved in THF (3 mL) and methanol (0.3 mL). Aqueous LiOH (1M, 5 eq) was added dropwise and the solution was stirred at room temperature overnight. The next day $NH_4Cl$ (satd., 1mL) was added and the solution was neutralized with HCl (12M) added dropwise. The neutralization was stirred for 1 hr before the solvent was removed *in vacuo*. The resulting solid was dissolved in $CHCl_3$, and the insoluble salts were filtered out. The crude product was concentrated again and then used in the following reaction without further purification.

*ii.* In a 20 mL scintillation vial, the boc protected amine (**10-15**) was dissolved in DCM. TFA was added dropwise at room temperature and the deprotection reaction was stirred for 30 minutes. Nitrogen was blown on the solution to evaporate the solvent and washed with DCM 3 times. The DCM was removed in between each wash. The crude product was carried through to the next step without further purification.

*iii.* To a new reaction vial was charged the crude amine (1 eq), **16** (1 eq), and DCM (0.5M). HATU (2 eq) and DIPEA (5 eq) were added stepwise and the resulting reaction mixture was stirred for 20 mins at room temperature. The reaction was concentrated *in vacuo* and purified via column chromatography (gradient elution, 0-10% Methanol in DCM).

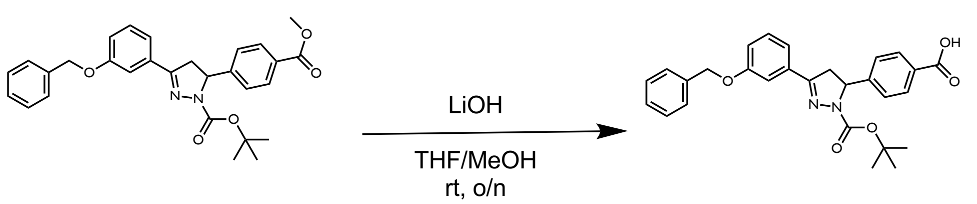


**(16)**

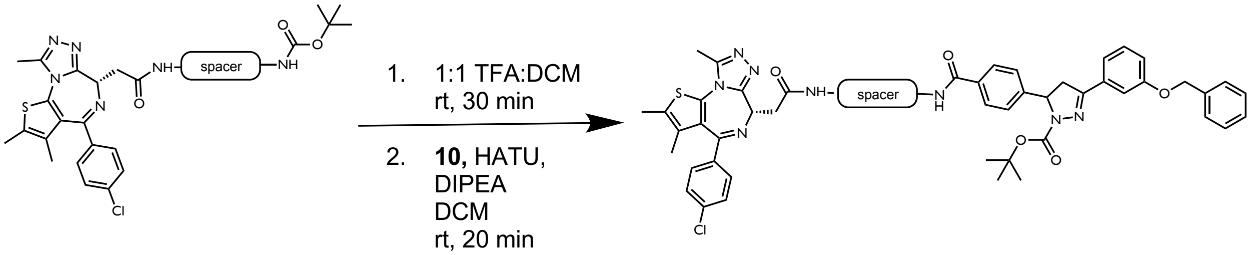


***tert*-butyl 3-(3-(benzyloxy)phenyl)-5-(4-((2-(2-((*S*)-4-(4-chlorophenyl)-2,3,9-trimethyl-6*H*-thieno[3,2-*f*][1,2,4]triazolo[4,3-*a*][1,4]diazepin-6-yl)acetamido)ethyl)carbamoyl)phenyl)-4,5-dihydro-1*H*-pyrazole-1-carboxylate (17)**

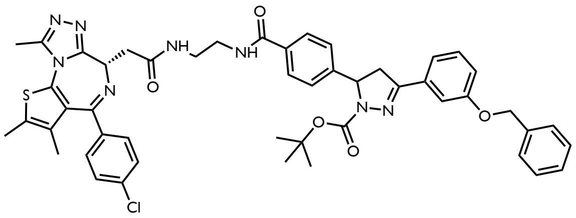

**10** (92 mg, 0.17 mmol) was coupled to **9** (80 mg, 0.17 mmol) following General Procedure B to yield **17** (136 mg, 0.15 mmol, 100%) as a white solid.
$^1$H NMR (400 MHz, $CDCl_3$) δ 7.78 (s, 3H), 7.48 – 7.19 (m, 20H), 7.00 (d, *J* = 8.0 Hz, 1H), 6.85 (s, 1H), 5.35 (s, 1H), 5.08 (s, 2H), 4.61 (s, 2H), 3.80 – 3.68 (m, 2H), 3.60 – 3.04 (m, 9H), 2.80 (s, 1H), 2.64 (s, 4H), 2.37 (s, 3H), 1.64 (s, 3H), 1.48 – 1.35 (m, 1H), 1.31 (s, 17H).

***tert*-butyl 3-(3-(benzyloxy)phenyl)-5-(4-((3-(2-((*S*)-4-(4-chlorophenyl)-2,3,9-trimethyl-6*H*-thieno[3,2-*f*][1,2,4]triazolo[4,3-*a*][1,4]diazepin-6-yl)acetamido)propyl)carbamoyl)phenyl)-4,5-dihydro-1*H*-pyrazole-1-carboxylate (18)**

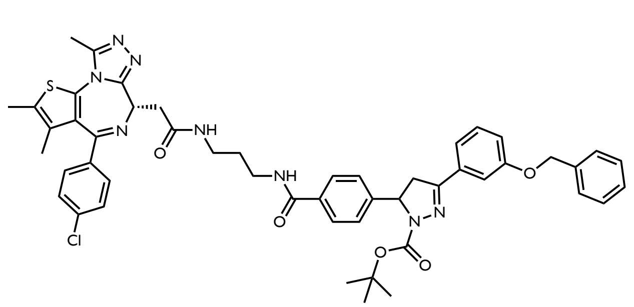

**11** (48 mg, 0.09 mmol) was coupled to **9** (37 mg, 0.09 mmol) following General Procedure B to yield **18** (62 mg, 0.07 mmol, 88%) as a white solid.
$^1$H NMR (400 MHz, $CDCl_3$) δ 7.88 – 7.80 (m, 2H), 7.75 (s, 1H), 7.57 – 7.11 (m, 10H), 6.99 (dt, *J* = 7.8, 2.1 Hz, 1H), 5.33 (dd, *J* = 12.3, 5.4 Hz, 1H), 5.07 (s, 2H), 4.64 (t, *J* = 6.9 Hz, 1H), 3.72 (dd, *J* = 17.6, 12.2 Hz, 1H), 3.54 (ddd, *J* = 14.6, 7.7, 2.5 Hz, 1H), 3.47 – 3.30 (m, 5H), 3.06 (dd, *J* = 17.6, 5.4 Hz, 1H), 2.64 (s, 3H), 2.36 (d, *J* = 5.1 Hz, 3H), 1.75 (d, *J* = 7.0 Hz, 2H), 1.62 (d, *J* = 5.6 Hz, 3H), 1.48 – 1.34 (m, 1H), 1.28 (s, 9H).

***tert*-butyl 3-(3-(benzyloxy)phenyl)-5-(4-((4-(2-((*S*)-4-(4-chlorophenyl)-2,3,9-trimethyl-6*H*-thieno[3,2-*f*][1,2,4]triazolo[4,3-*a*][1,4]diazepin-6-yl)acetamido)butyl)carbamoyl)phenyl)-4,5-dihydro-1*H*-pyrazole-1-carboxylate (19)**

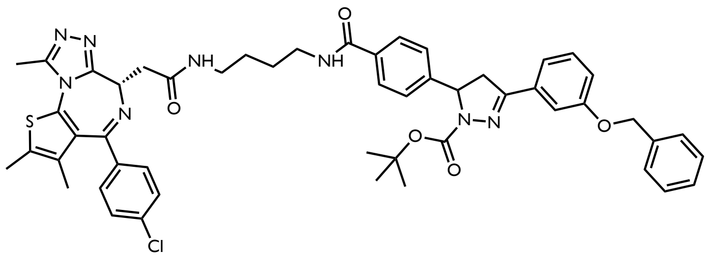

**12** (22 mg, 0.04 mmol) was coupled to **9** (17 mg, 0.04 mmol) following General Procedure B to yield **19** (27 mg, 0.03 mmol, 75%) as a white solid.
$^1$H NMR (400 MHz, $CDCl_3$) δ 7.88 – 7.80 (m, 2H), 7.75 (s, 1H), 7.57 – 7.11 (m, 10H), 6.99 (dt, *J* = 7.8, 2.1 Hz, 1H), 5.33 (dd, *J* = 12.3, 5.4 Hz, 1H), 5.07 (s, 2H), 4.64 (t, *J* = 6.9 Hz, 1H), 3.72 (dd, *J* = 17.6, 12.2 Hz, 1H), 3.54 (ddd, *J* = 14.6, 7.7, 2.5 Hz, 1H), 3.47 – 3.30 (m, 5H), 3.06 (dd, *J* = 17.6, 5.4 Hz, 1H), 2.64 (s, 3H), 2.36 (d, *J* = 5.1 Hz, 3H), 1.75 (d, *J* = 7.0 Hz, 2H), 1.62 (d, *J* = 5.6 Hz, 3H), 1.48 – 1.34 (m, 1H), 1.28 (s, 9H).

***tert*-butyl 3-(3-(benzyloxy)phenyl)-5-(4-((5-(2-((*S*)-4-(4-chlorophenyl)-2,3,9-trimethyl-6*H*-thieno[3,2-*f*][1,2,4]triazolo[4,3-*a*][1,4]diazepin-6-yl)acetamido)pentyl)carbamoyl)phenyl)-4,5-dihydro-1*H*-pyrazole-1-carboxylate (20)**

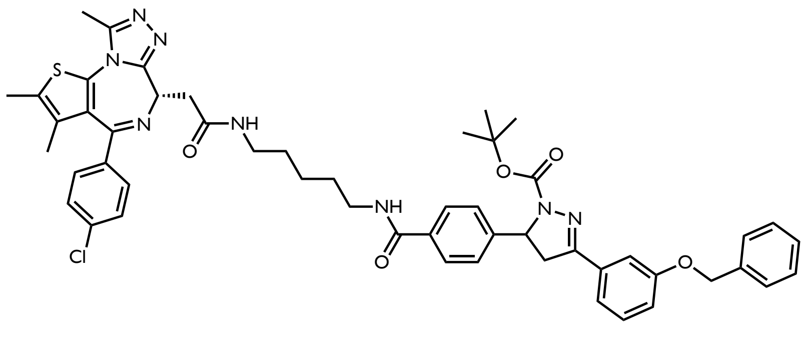

**13** (60.0 mg, 0.12 mmol) was coupled to **9** (59 mmol, 0.12 mmol) following General Procedure B to yield **20** (17 mg, 0.02 mmol, 15%) as a white solid.
$^1$H NMR (400 MHz, $CDCl_3$) δ 7.84 (d, *J* = 5.8 Hz, 2H), 7.56 – 7.28 (m, 12H), 7.23 (dd, *J* = 8.3, 4.8 Hz, 3H), 7.16 – 6.95 (m, 2H), 6.80 (s, 1H), 5.34 (d, *J* = 11.7 Hz, 1H), 5.07 (d, *J* = 4.9 Hz, 2H), 4.60 (d, *J* = 8.7 Hz, 1H), 3.73 (t, *J* = 14.9 Hz, 1H), 3.64 – 2.98 (m, 8H), 2.63 (d, *J* = 4.8 Hz, 3H), 2.38 (d, *J* = 5.0 Hz, 3H), 1.70 – 1.36 (m, 8H), 1.26 (d, *J* = 11.3 Hz, 11H).

***tert*-butyl 3-(3-(benzyloxy)phenyl)-5-(4-((6-(2-((*S*)-4-(4-chlorophenyl)-2,3,9-trimethyl-6*H*-thieno[3,2-*f*][1,2,4]triazolo[4,3-*a*][1,4]diazepin-6-yl)acetamido)hexyl)carbamoyl)phenyl)-4,5-dihydro-1*H*-pyrazole-1-carboxylate (21)**

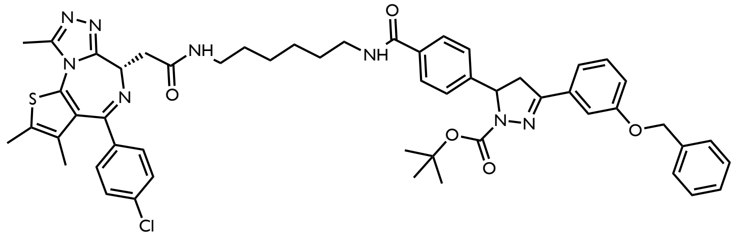

**14** (94 mg, 0.16 mmol) was coupled to **9** (74 mg, 0.16 mmol) following General Procedure B to yield **21** (33 mg, 0.03 mmol, 20%) as a white solid.
$^1$H NMR (400 MHz, $CDCl_3$) δ 5.39 – 5.26 (m, 1H), 5.07 (d, *J* = 4.1 Hz, 2H), 4.71 – 4.62 (m, 1H), 4.23 (s, 3H), 3.85 – 3.19 (m, 5H), 3.17 – 2.94 (m, 1H), 2.52 (d, *J* = 16.1 Hz, 3H), 2.34 (d, *J* = 5.8 Hz, 3H), 1.68 – 1.53 (m, 3H), 1.52 – 0.72 (m, 11H).

***tert*-butyl 3-(3-(benzyloxy)phenyl)-5-(4-((7-(2-((*S*)-4-(4-chlorophenyl)-2,3,9-trimethyl-6*H*-thieno[3,2-*f*][1,2,4]triazolo[4,3-*a*][1,4]diazepin-6-yl)acetamido)heptyl)carbamoyl)phenyl)-4,5-dihydro-1*H*-pyrazole-1-carboxylate (22)**

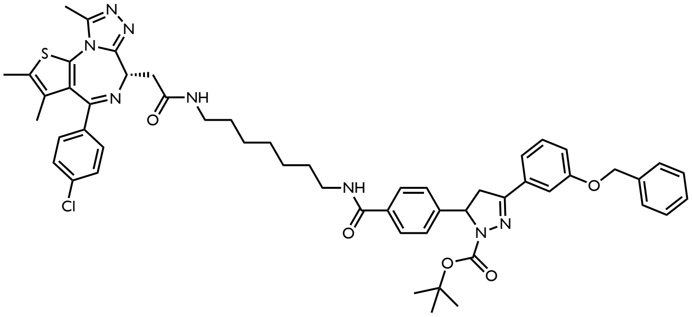

**15** (64 mg, 0.1 mmol) was coupled to **9** (47 mg, 0.1 mmol) following General Procedure B to yield **22** (5.6 mg, 0.01 mmol, 7%) as a white solid.
$^1$H NMR (400 MHz, $CDCl_3$) δ 8.66 – 8.60 (m, 1H), 8.30 (dd, *J* = 8.4, 1.4 Hz, 1H), 7.77 (d, *J* = 7.8 Hz, 2H), 7.52 – 7.18 (m, 18H), 7.03 – 6.97 (m, 1H), 6.95 (s, 1H), 5.35 (s, 1H), 5.08 (s, 2H), 4.67 (dd, *J* = 7.8, 5.8 Hz, 1H), 3.72 (td, *J* = 17.0, 9.3 Hz, 2H), 3.58 – 3.45 (m, 1H), 3.42 – 3.19 (m, 5H), 3.19 – 3.04 (m, 2H), 2.80 (s, 5H), 2.66 (s, 3H), 2.39 (s, 3H), 1.66 (s, 4H), 1.60 – 1.46 (m, 1H), 1.46 – 1.37 (m, 5H), 1.30 (s, 9H).

**General Procedure C: Preparation of chloroacetamide warhead containing JQ1 based bifunctionals.**

i. In a 20 mL scintillation vial, the boc protected amine (**17-22**) was dissolved in DCM. TFA was added dropwise at room temperature and the deprotection reaction was stirred for 30 minutes. The solvent was removed under a stream of nitrogen and washed with DCM 3 times. The DCM was removed by evaporation in between each wash. The crude product was carried through to the next step without further purification.
ii. To a new reaction vial was charged the crude amine and dissolved in DCM. The solution was cooled to -10$^o$C. Chloroacetyl chloride (1.5 eq) was added dropwise before TEA (3 eq) was added and the reaction mixture was warmed to room temperature. The reaction was stirred for 30 minutes before it was concentrated *in vacuo* and purified via column chromatography (gradient elution, 0-10% methanol in DCM).

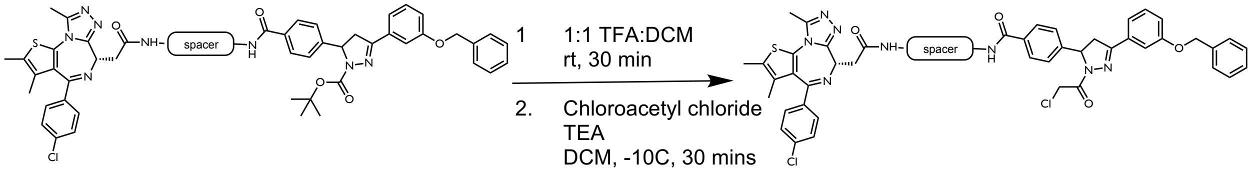


**4-(3-(3-(benzyloxy)phenyl)-1-(2-chloroacetyl)-4,5-dihydro-1*H*-pyrazol-5-yl)-*N*-(2-(2-((*S*)-4-(4-chlorophenyl)-2,3,9-trimethyl-6*H*-thieno[3,2-*f*][1,2,4]triazolo[4,3-*a*][1,4]diazepin-6-yl)acetamido)ethyl)benzamide (22)**

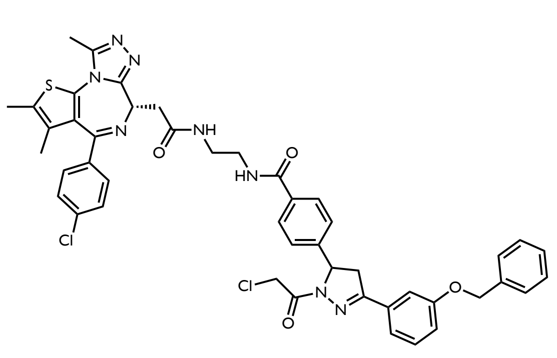

The warhead was added to **17** (136 mg, 0.15 mmol) following General Procedure C to yield **22** (3.3 mg, 0.004 mmol, 2.5%) as a white solid.
$^{1}$H NMR (600 MHz, Acetone) δ 7.84 (dd, *J* = 11.2, 8.1 Hz, 2H), 7.54 – 7.27 (m, 14H), 7.15 (dddd, *J* = 7.9, 3.8, 2.6, 1.2 Hz, 1H), 5.67 (dd, *J* = 11.9, 5.0 Hz, 1H), 5.19 (d, *J* = 2.7 Hz, 2H), 4.72 (dd, *J* = 13.7, 1.2 Hz, 1H), 4.67 – 4.61 (m, 2H), 4.03 – 3.93 (m, 1H), 3.48 – 3.36 (m, 3H), 3.35 – 3.18 (m, 4H), 2.57 (d, *J* = 2.6 Hz, 3H), 2.50 – 2.30 (m, 3H), 1.70 – 1.56 (m, 3H).
$^{13}$C NMR (151 MHz, Acetone) δ 163.57, 163.41, 159.11, 155.22, 144.84, 137.30, 137.22, 135.78, 135.75, 134.10, 132.66, 132.54, 130.78, 130.45, 130.20, 130.18, 129.90, 128.45, 128.35, 128.33, 127.88, 127.74, 127.67, 125.62, 119.62, 117.21, 113.10, 69.77, 60.26, 54.42, 53.67, 45.65, 42.10, 42.08, 41.89, 40.21, 40.15, 38.74, 38.44, 29.68, 29.56, 29.44, 13.64, 12.12, 12.09, 10.88, 7.93; HRMS (ES+) calcd for $C_{46}H_{42}Cl_2N_8O_4S$ [M+H]$^+$ 873.2427, found 873.2501

**4-(3-(3-(benzyloxy)phenyl)-1-(2-chloroacetyl)-4,5-dihydro-1*H*-pyrazol-5-yl)-*N*-(3-(2-((*S*)-4-(4-chlorophenyl)-2,3,9-trimethyl-6*H*-thieno[3,2-*f*][1,2,4]triazolo[4,3-*a*][1,4]diazepin-6-yl)acetamido)propyl)benzamide (23)**

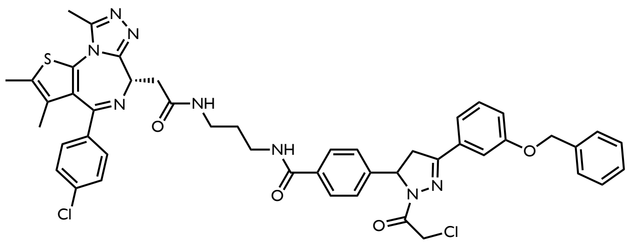

The warhead was added to **18** (62 mg, 0.07 mmol) following General Procedure C to yield **23** (12.2 mg, 0.013 mmol, 36%) as a white solid.
$^{1}$H NMR (500 MHz, Acetone) δ 7.99 (q, *J* = 6.7 Hz, 1H), 7.84 (dd, *J* = 8.2, 6.3 Hz, 2H), 7.74 (t, *J* = 6.3 Hz, 1H), 7.54 – 7.29 (m, 19H), 7.15 (d, *J* = 8.0 Hz, 1H), 5.67 (dd, *J* = 11.9, 4.9 Hz, 1H), 5.19 (s, 2H), 4.72 (d, *J* = 13.7 Hz, 1H), 4.69 – 4.60 (m, 2H), 4.02 – 3.92 (m, 1H), 3.59 – 3.48 (m, 1H), 3.39 (qd, *J* = 12.3, 6.6 Hz, 6H), 3.31 – 3.21 (m, 1H), 2.59 (d, *J* = 2.6 Hz, 4H), 2.41 (d, *J* = 13.7 Hz, 3H), 1.75 (p, *J* = 6.5 Hz, 2H), 1.63 (d, *J* = 15.0 Hz, 3H), 1.44-1.40 (m, 2H). $^{13}$C NMR (126 MHz, Acetone) δ 170.66, 163.45, 163.38, 159.10, 155.58, 155.24, 149.67, 144.79, 137.34, 137.23, 135.78, 134.48, 132.73, 132.57, 130.75, 130.38, 130.18, 129.89, 128.46, 128.38, 127.88, 127.69, 127.57, 125.67, 125.65, 119.61, 117.20, 113.07, 69.76, 60.26, 54.37, 42.10, 38.37, 36.01, 35.96, 35.77, 29.70, 29.51, 29.35, 29.20, 13.60, 12.11, 12.09, 10.87; HRMS (ES+) calcd for $C_{47}H_{44}Cl_2N_8O_4S$ [M+H]$^+$ 887.2583, found 887.2662

**4-(3-(3-(benzyloxy)phenyl)-1-(2-chloroacetyl)-4,5-dihydro-1*H*-pyrazol-5-yl)-*N*-(4-(2-((*S*)-4-(4-chlorophenyl)-2,3,9-trimethyl-6*H*-thieno[3,2-*f*][1,2,4]triazolo[4,3-*a*][1,4]diazepin-6-yl)acetamido)butyl)benzamide (24)**

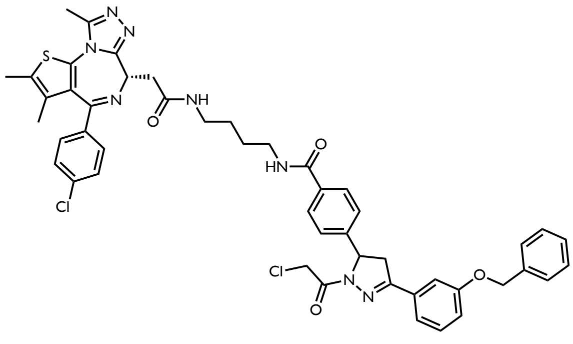

The warhead was added to **19** (27 mg, 0.03 mmol) following General Procedure C to yield **24** (9.0 mg, 0.01 mmol, 34%) as a white solid.

¹H NMR (600 MHz, Acetone) δ 8.08 – 7.69 (m, 3H), 7.53 – 7.47 (m, 6H), 7.46 – 7.36 (m, 6H), 7.33 (t, *J* = 7.4 Hz, 1H), 7.30 – 7.25 (m, 2H), 7.17 – 7.12 (m, 1H), 5.64 (d, *J* = 10.9 Hz, 1H), 5.19 (d, *J* = 2.3 Hz, 2H), 4.74 – 4.59 (m, 3H), 3.96 (t, *J* = 15.2 Hz, 1H), 3.48 – 3.14 (m, 7H), 2.60 (s, 3H), 2.41 (s, 3H), 2.09 – 2.05 (m, 1H), 1.67 (s, 3H), 1.42 (d, *J* = 12.5 Hz, 2H). ¹³C NMR (151 MHz, Acetone) δ 163.45, 163.37, 159.11, 155.32, 144.76, 137.32, 137.23, 135.81, 132.54, 130.81, 130.35, 130.22, 129.88, 129.68, 129.66, 128.45, 128.36, 127.90, 127.87, 127.68, 125.53, 119.62, 117.24, 117.22, 113.09, 69.78, 60.30, 54.50, 42.12, 42.07, 42.03, 39.52, 38.50, 28.40, 23.62, 22.43, 13.67, 13.45, 12.12, 10.90; HRMS (ES+) calcd for $C_{48}H_{46}Cl_2N_8O_4S$ $[M+H]^+$ 901.2740, found 901.2833

**4-(3-(3-(benzyloxy)phenyl)-1-(2-chloroacetyl)-4,5-dihydro-1*H*-pyrazol-5-yl)-*N*-(5-(2-((*S*)-4-(4-chlorophenyl)-2,3,9-trimethyl-6*H*-thieno[3,2-*f*][1,2,4]triazolo[4,3-*a*][1,4]diazepin-6-yl)acetamido)pentyl)benzamide (25)**

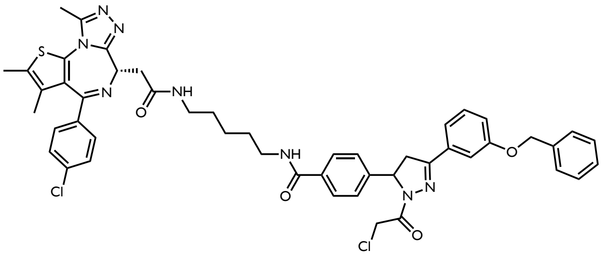

The warhead was added to **20** (17 mg, 0.02 mmol) following General Procedure C to yield **25** (1.4 mg, 0.002 mmol, 11%) as a white solid.
¹H NMR (600 MHz, Acetone) δ 7.89 – 7.11 (m, 17H), 7.01 (ddd, *J* = 7.8, 2.6, 1.3 Hz, 1H), 5.50 (dt, *J* = 11.9, 4.4 Hz, 1H), 5.06 (t, *J* = 1.9 Hz, 2H), 4.62 – 4.44 (m, 3H), 3.82 (ddd, *J* = 17.2, 11.8, 3.6 Hz, 1H), 3.32 – 3.05 (m, 9H), 2.46 (d, *J* = 4.5 Hz, 3H), 2.30 – 2.26 (m, 3H), 1.95 (d, *J* = 1.6 Hz, 2H), 1.94 – 1.93 (m, 3H), 1.54 (d, *J* = 4.8 Hz, 3H), 1.19 (td, *J* = 7.3, 1.5 Hz, 3H).
¹³C NMR (151 MHz, Acetone) δ 163.44, 159.11, 155.48, 155.31, 155.26, 144.88, 144.82, 137.29, 137.22, 135.87, 132.52, 130.85, 130.35, 130.32, 130.21, 130.14, 129.88, 128.45, 128.38, 127.87, 127.85, 127.79, 127.67, 125.57, 125.54, 119.61, 117.67, 117.22, 117.19, 115.73, 113.09, 69.78, 60.28, 54.46, 54.43, 45.77, 42.09, 42.07, 42.04, 42.02, 39.44, 38.47, 23.67, 23.61, 13.63, 12.10, 10.88, 7.95; HRMS (ES+) calcd for $C_{49}H_{48}Cl_2N_8O_4S$ $[M+H]^+$ 915.2896, found 915.2969

**4-(3-(3-(benzyloxy)phenyl)-1-(2-chloroacetyl)-4,5-dihydro-1*H*-pyrazol-5-yl)-*N*-(6-(2-((*S*)-4-(4-chlorophenyl)-2,3,9-trimethyl-6*H*-thieno[3,2-*f*][1,2,4]triazolo[4,3-*a*][1,4]diazepin-6-yl)acetamido)hexyl)benzamide (26)**

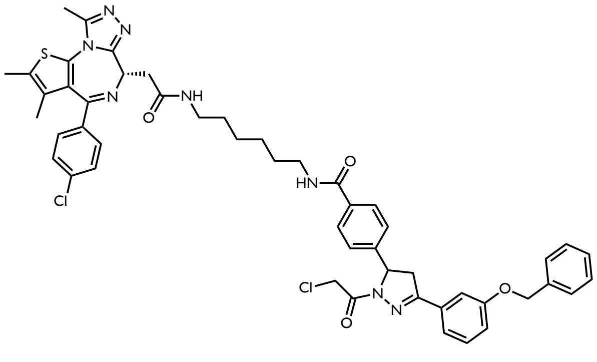

The warhead was added to **21** (33 mg, 0.03 mmol) following General Procedure C to yield **26** (6.0 mg, 0.006 mmol, 20%) as a white solid.
¹H NMR (600 MHz, Acetone) δ 7.87 – 7.83 (m, 3H), 7.53 – 7.46 (m, 5H), 7.44 – 7.38 (m, 6H), 7.38 – 7.27 (m, 4H), 7.14 (ddt, *J* = 7.9, 2.6, 1.3 Hz, 1H), 5.65 (dd, *J* = 10.6, 5.6 Hz, 1H), 5.18 (d, *J* = 1.9 Hz, 2H), 4.71 (dd, *J* = 13.6, 2.0 Hz, 1H), 4.68 – 4.59 (m, 2H), 4.01 – 3.91 (m, 1H), 3.43 – 3.37 (m, 1H), 3.37 – 3.20 (m, 6H), 2.93 (s, 1H), 2.86 – 2.79 (m, 1H), 2.78 (d, *J* = 0.6 Hz, 1H), 2.58 (s, 3H), 2.37 (dd, *J* = 2.8, 1.1 Hz, 3H), 1.65 (d, *J* = 2.5 Hz, 3H), 1.60 – 1.49 (m, 5H). ¹³C NMR (151 MHz, Acetone) δ 170.81, 166.83, 166.76, 164.28, 164.23, 162.72, 159.98, 156.50, 156.14, 150.54, 145.48, 138.21, 138.10, 136.71, 135.50, 135.47, 133.57, 133.43, 131.69, 131.16, 131.14, 131.05, 131.00, 130.76, 129.34, 129.26, 128.76, 128.56, 128.53, 126.44, 120.49, 118.08, 113.96, 70.66, 61.16, 55.34, 42.99, 39.67, 39.55, 39.27, 39.22, 39.09, 36.13, 26.69, 26.67, 26.53, 14.52, 12.98, 11.76, 11.74; HRMS (ES+) calcd for $C_{50}H_{50}Cl_2N_8O_4S$ $[M+H]^+$ 929.3053, found 929.3131

**4-(3-(3-(benzyloxy)phenyl)-1-(2-chloroacetyl)-4,5-dihydro-1*H*-pyrazol-5-yl)-*N*-(7-(2-((*S*)-4-(4-chlorophenyl)-2,3,9-trimethyl-6*H*-thieno[3,2-*f*][1,2,4]triazolo[4,3-*a*][1,4]diazepin-6-yl)acetamido)heptyl)benzamide (28)**

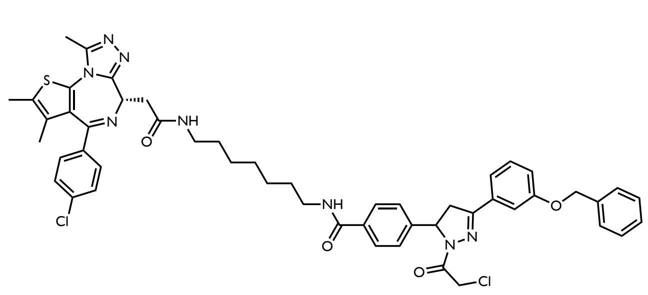

The warhead was added to **22** (5.6 mg, 0.006 mmol) following General Procedure C to yield **28** (3.8 mg, 0.004 mmol, 67%) as a white solid.
$^{1}$H NMR (600 MHz, Acetone) δ 7.86 – 7.81 (m, 2H), 7.78 (s, 1H), 7.54 – 7.46 (m, 6H), 7.45 – 7.30 (m, 11H), 7.18 – 7.11 (m, 1H), 5.66 (dt, *J* = 11.9, 5.2 Hz, 1H), 5.19 (d, *J* = 1.9 Hz, 2H), 4.71 (dd, *J* = 13.6, 1.8 Hz, 1H), 4.65 – 4.58 (m, 2H), 4.01 – 3.93 (m, 1H), 3.42 – 3.32 (m, 4H), 3.31 – 3.22 (m, 4H), 2.58 (d, *J* = 1.0 Hz, 3H), 2.44 (s, 3H), 2.08 (t, *J* = 0.9 Hz, 1H), 2.09 – 2.03 (m, 10H), 1.70 (t, *J* = 0.9 Hz, 4H), 1.58 (q, *J* = 6.8 Hz, 4H), 1.52 (q, *J* = 6.2 Hz, 1H), 1.36 (d, *J* = 5.2 Hz, 8H). $^{13}$C NMR (151 MHz, Acetone) δ 169.97, 166.39, 163.79, 163.64, 159.51, 156.01, 155.63, 150.02, 145.05, 137.78, 137.63, 136.21, 135.10, 133.13, 132.97, 131.16, 130.71, 130.58, 130.55, 130.27, 128.85, 128.76, 128.27, 128.08, 128.00, 126.03, 120.00, 117.61, 113.47, 70.17, 60.68, 54.86, 42.48, 42.46, 39.75, 39.06, 38.82, 29.87, 29.84, 27.01, 26.80, 26.79, 14.03, 13.85, 12.49, 11.24; HRMS (ES+) calcd for $C_{51}H_{52}Cl_2N_8O_4S$ $[M+H]^+$ 943.32093, found 943.32898

**4-(3-(3-(benzyloxy)phenyl)-1-propionyl-4,5-dihydro-1*H*-pyrazol-5-yl)-*N*-(5-(2-((*S*)-4-(4-chlorophenyl)-2,3,9-trimethyl-6*H*-thieno[3,2-*f*][1,2,4]triazolo[4,3-*a*][1,4]diazepin-6-yl)acetamido)pentyl)benzamide (27)**

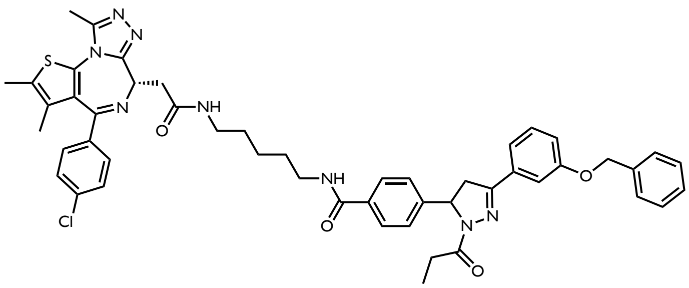

In a 20 mL scintillation vial, the boc protected amine (**20**) (42 mg, 0.04 mmol) was dissolved in DCM. TFA was added dropwise at room temperature and the deprotection reaction was stirred for 30 minutes. The solvent was removed under a stream of nitrogen and washed with DCM 3 times. The DCM was evaporated in between each wash. The crude product was carried through to the next step without further purification. To a new reaction vial was charged the crude amine and dissolved in DCM. The solution was cooled to -10ºC and propionyl chloride (1.5 eq) was added dropwise before TEA (3 eq) was added and the reaction mixture was warmed to room temperature. The reaction was stirred for 30 minutes before it was concentrated *in vacuo* and purified via column chromatography (gradient elution, 0-10% methanol in DCM) to yield **27** (17 mg, 0.02 mmol, 50%).
$^{1}$H NMR (600 MHz, Acetone) δ 7.88 – 7.83 (m, 2H), 7.80 (s, 1H), 7.55 – 7.45 (m, 6H), 7.44 – 7.29 (m, 7H), 7.27 – 7.21 (m, 2H), 7.11 (ddd, *J* = 7.5, 2.6, 1.6 Hz, 1H), 5.59 (ddd, *J* = 12.0, 5.1, 3.0 Hz, 1H), 5.19 (d, *J* = 2.6 Hz, 2H), 4.60 (ddd, *J* = 7.5, 6.5, 2.0 Hz, 1H), 3.87 (ddd, *J* = 17.9, 12.1, 1.0 Hz, 1H), 3.44 – 3.20 (m, 5H), 3.14 (ddd, *J* = 17.9, 5.1, 2.8 Hz, 1H), 2.79 (s, 1H), 2.82 – 2.70 (m, 2H), 2.59 (d, *J* = 4.2 Hz, 3H), 2.42 (dd, *J* = 2.0, 0.9 Hz, 3H), 2.09 (s, 1H), 1.67 (t, *J* = 1.0 Hz, 3H), 1.68 – 1.58 (m, 1H), 1.61 – 1.50 (m, 2H), 1.50 – 1.39 (m, 2H), 1.30 (s, 2H), 1.29 (s, 1H), 1.10 (td, *J* = 7.5, 1.9 Hz, 3H). $^{13}$C NMR (151 MHz, Acetone) δ 171.91, 170.83 (d, *J* = 2.4 Hz), 167.00, 164.19, 159.99, 156.44 (d, *J* = 2.5 Hz), 154.27, 150.57 (d, *J* = 2.2 Hz), 146.58, 138.24, 138.19, 136.71, 136.70, 135.15, 135.10, 134.04, 133.61 (d, *J* = 3.4 Hz), 131.66, 131.63, 131.22, 131.19, 131.07, 131.03, 131.01, 130.69, 129.33, 129.28, 129.26, 128.74, 128.54, 128.51, 126.33, 126.30, 120.19, 117.62, 117.60, 113.70, 70.63, 60.73, 55.41, 42.75, 40.35, 40.32, 39.42, 39.40, 27.98, 27.97, 24.82, 14.52, 12.99, 11.76, 9.36; HRMS (ES+) calcd for $C_{50}H_{51}ClN_8O_4S$ $[M+H]^+$ 895.34425, found 895.35260

**4-(1-acryloyl-3-(3-(benzyloxy)phenyl)-4,5-dihydro-1*H*-pyrazol-5-yl)-*N*-(5-(2-((*S*)-4-(4-chlorophenyl)-2,3,9-trimethyl-6*H*-thieno[3,2-*f*][1,2,4]triazolo[4,3-*a*][1,4]diazepin-6-yl)acetamido)pentyl)benzamide (41)**

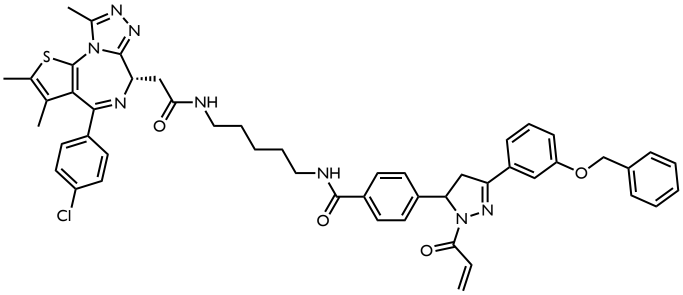

In a 20 mL scintillation vial, the boc protected amine (**20**) (10 mg, 0.01 mmol) was dissolved in DCM. TFA was added dropwise at room temperature and the deprotection reaction was stirred for 30 minutes. The solvent was removed under a stream of nitrogen and washed with DCM 3 times. The DCM was evaporated in between each wash. The crude product was carried through to the next step without further purification. To a new reaction vial was charged the crude amine and dissolved in DCM. The solution was cooled to -10ºC and propionyl chloride (1.5 eq) was added dropwise before TEA (3 eq) was added and the reaction mixture was warmed to room temperature. The reaction was stirred for 30 minutes before it was concentrated *in vacuo* and purified via column chromatography (gradient elution, 0-10% methanol in DCM) to yield **41** (2.4 mg, 0.003 mmol, 30%).
$^1$H NMR (600 MHz, $CDCl_3$) δ 7.88 – 7.67 (m, 2H), 7.50 – 7.29 (m, 11H), 7.22 – 7.14 (m, 2H), 7.08 (d, *J* = 7.1 Hz, 1H), 6.82 – 6.70 (m, 1H), 6.34 (t, *J* = 15.0 Hz, 1H), 5.78 (d, *J* = 10.6 Hz, 1H), 5.64 – 5.26 (m, 1H), 5.14 (d, *J* = 2.3 Hz, 2H), 4.70 – 4.58 (m, 1H), 3.83 – 2.85 (m, 13H), 2.73 (d, *J* = 2.3 Hz, 3H), 2.44 (d, *J* = 4.2 Hz, 3H), 1.75 – 1.66 (m, 3H), 1.52 – 1.25 (m, 4H); Carbon was unable to be acquired due to low signal. HRMS (ES+) calcd for $C_{50}H_{49}ClN_8O_4S$ $[M+H]^+$ 893.3286, found 893.3342

***tert*-butyl 4-(6-((4-(3-chloro-4-cyanophenoxy)cyclohexyl)carbamoyl)pyridazin-3-yl)piperazine-1-carboxylate (28)**

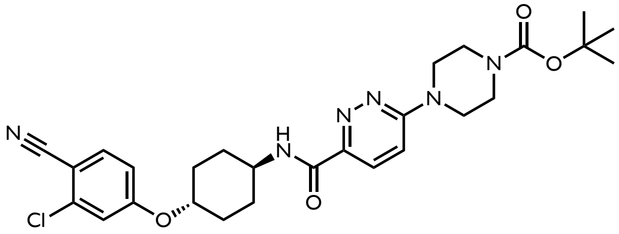

**28** was synthesized following the procedures reported by Forte and coworkers (ACS Chem. Bio., **2023**, 18, 897-904).

***tert*-butyl 2-(4-(6-((4-(3-chloro-4-cyanophenoxy)cyclohexyl)carbamoyl)pyridazin-3-yl)piperazin-1-yl)acetate (29)**

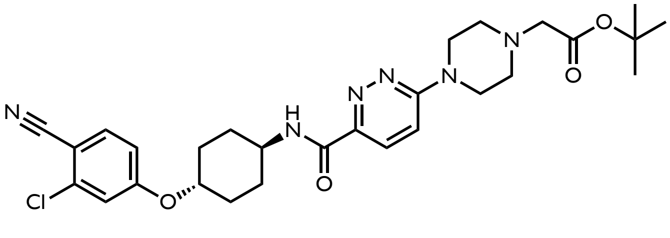

**29** was synthesized following the procedures reported by Forte and coworkers (ACS Chem. Bio., **2023**, 18, 897-904).

**General Procedure D: Preparation of linkers coupled to androgen receptor ligand**

*i.* To a 20 mL scintillation vial, **29** (1 eq) was dissolved in DCM, before a N-boc diamino alkane (1 eq), HATU (2 eq) and DIPEA (5 eq) were added stepwise. The resulting reaction mixture was stirred for 20 mins at room temperature. The reaction was concentrated *in vacuo* and purified via column chromatography (gradient elution, 0-10% Methanol in DCM).

*ii.*

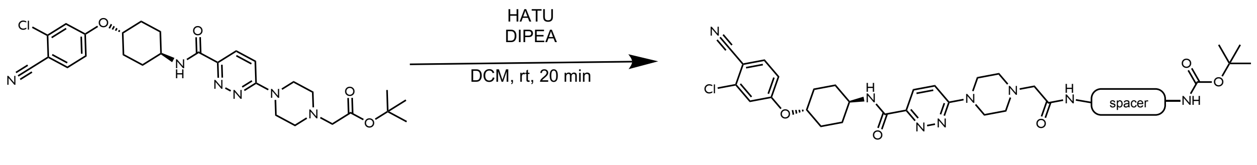

**tert-butyl (2-(2-(4-(6-((4-(3-chloro-4-cyanophenoxy)cyclohexyl)carbamoyl)pyridazin-3-yl)piperazin-1-yl)acetamido)ethyl)carbamate (30)**

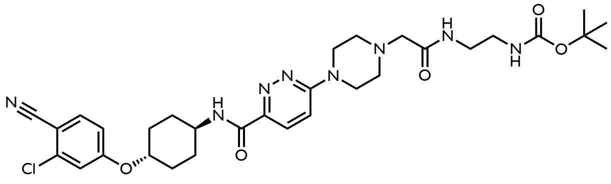

**29** (25 mg, 0.04 mmol) was coupled to N-Boc-ethylenediamine ( following General Procedure D to yield **30** (25 mg, 0.04 mmol, 80%) as a yellow solid.
$^1$H NMR (400 MHz, $CDCl_3$) δ 8.06 (d, *J* = 9.2 Hz, 1H), 7.90 (d, *J* = 8.1 Hz, 1H), 7.65 – 7.56 (m, 1H), 7.03 (dd, *J* = 5.7, 3.3 Hz, 2H), 6.89 (dd, *J* = 8.6, 2.5 Hz, 1H), 4.37 (d, *J* = 10.1 Hz, 1H), 4.09 (d, *J* = 9.5 Hz, 1H), 3.94 – 3.81 (m, 4H), 3.45 (q, *J* = 4.7 Hz, 4H), 3.20 (s, 2H), 3.05 (s, 4H), 2.80 (s, 4H), 1.78 – 1.65 (m, 2H), 1.47 (s, 9H).

**tert-butyl (4-(2-(4-(6-((4-(3-chloro-4-cyanophenoxy)cyclohexyl)carbamoyl)pyridazin-3-yl)piperazin-1-yl)acetamido)butyl)carbamate (31)**

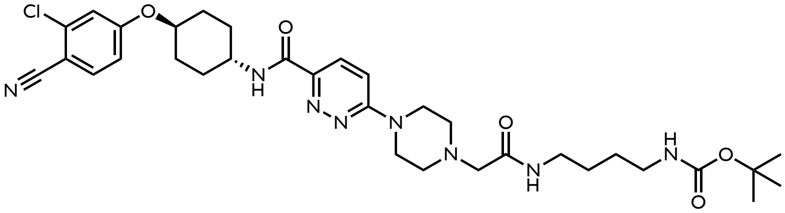

**29** was coupled to N-Boc-1,3-pentanediamine following General Procedure D to yield **31** (10.3 mg, 0.02 mmol, 47%) as a yellow solid.
$^1$H NMR (400 MHz, $CDCl_3$) δ 8.06 (d, *J* = 9.5 Hz, 1H), 7.90 (d, *J* = 8.2 Hz, 1H), 7.59 (d, *J* = 8.6 Hz, 1H), 7.25 (s, 1H), 7.05 (d, *J* = 8.9 Hz, 2H), 6.89 (d, *J* = 8.8 Hz, 1H), 4.37 (d, *J* = 10.3 Hz, 1H), 4.09 (s, 1H), 3.87 (s, 4H), 3.36 (d, *J* = 6.7 Hz, 2H), 3.18 (d, *J* = 9.2 Hz, 4H), 3.03 (s, 4H), 2.80 (s, 4H), 1.75 – 1.53 (m, 8H), 1.46 (s, 9H).

**tert-butyl (5-(2-(4-(6-((4-(3-chloro-4-cyanophenoxy)cyclohexyl)carbamoyl)pyridazin-3-yl)piperazin-1-yl)acetamido)pentyl)carbamate (32)**

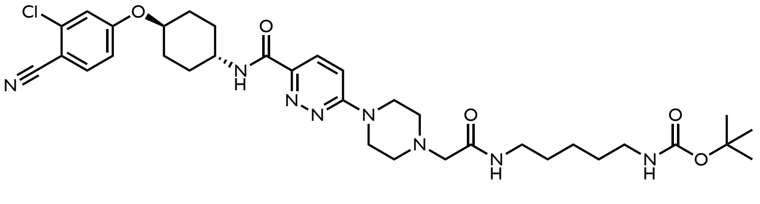

**29** was coupled to N-Boc-1,5-pentanediamine following General Procedure D to yield **32** (60 mg, 0.09 mmol, 92%) as a yellow solid.
$^1$H NMR (400 MHz, $CDCl_3$) δ 8.06 (d, *J* = 9.5 Hz, 1H), 7.90 (d, *J* = 8.2 Hz, 1H), 7.59 (d, *J* = 8.7 Hz, 1H), 7.10 – 7.01 (m, 2H), 6.89 (dd, *J* = 8.7, 2.4 Hz, 1H), 4.36 (dt, *J* = 10.7, 6.4 Hz, 1H), 4.09 (d, *J* = 10.1 Hz, 1H), 3.89 (d, *J* = 6.4 Hz, 4H), 3.38 – 3.20 (m, 4H), 3.17 – 3.09 (m, 4H), 3.03 (s, 5H), 2.84 (s, 3H), 1.70 (dd, *J* = 13.1, 9.0 Hz, 3H), 1.59 (s, 1H), 1.46 (d, *J* = 3.4 Hz, 9H).

**General Procedure E: Preparation of the Boc protected DDB1 recruiter linked to AR ligand-linker derivatives**

*i.* **9** (1 eq) was charged to a 20 mL scintillation vial and dissolved in THF (3 mL) and methanol (0.3 mL). LiOH (1M aq, 5 eq) was added dropwise and the solution was stirred at room temperature overnight. The next day $NH_4Cl$ (satd., 1mL) was added and the solution was neutralized with HCl (12M) added dropwise. The neutralization was stirred for 1hr before the solvent was removed *in vacuo*. The resulting solid was dissolved in $CHCl_3$, and the insoluble salts were filtered out. The crude product was concentrated again and then used in the following reaction without further purification.

*ii.* In a 20 mL scintillation vial, the boc protected amine (**30-32**) was dissolved in DCM. TFA was added dropwise at room temperature and the deprotection reaction was stirred for 30 minutes. Nitrogen was blown on the solution to evaporate the solvent and washed with DCM 3 times. The DCM was removed in between each wash. The crude product was carried through to the next step without further purification.

*iii.* To a new reaction vial was charged the crude amine (1 eq), **16** (1 eq), and DCM (0.5M). HATU (2 eq) and DIPEA (5 eq) were added stepwise and the resulting reaction mixture was stirred for 20 mins at room temperature. The reaction was concentrated *in vacuo* and purified via column chromatography (gradient elution, 0-10% Methanol in DCM).

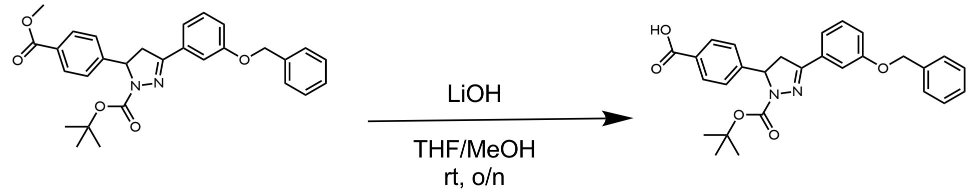


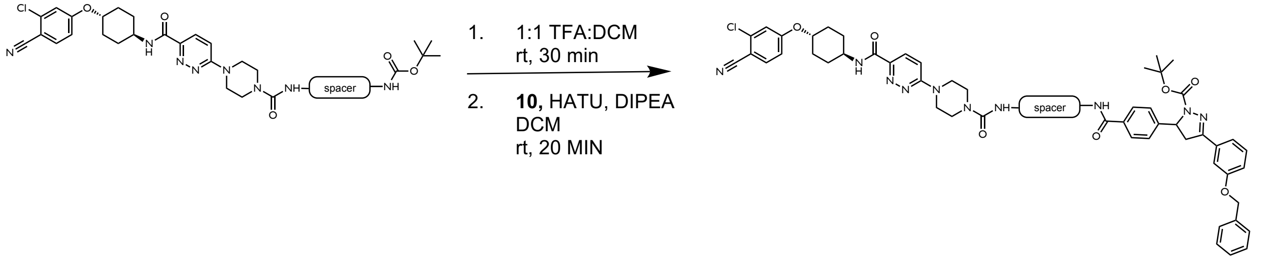


**tert-butyl 3-(3-(benzyloxy)phenyl)-5-(4-((2-(2-(4-(6-((4-(3-chloro-4-cyanophenoxy)cyclohexyl)carbamoyl)pyridazin-3-yl)piperazin-1-yl)acetamido)ethyl)carbamoyl)phenyl)-4,5-dihydro-1*H*-pyrazole-1-carboxylate (33)**

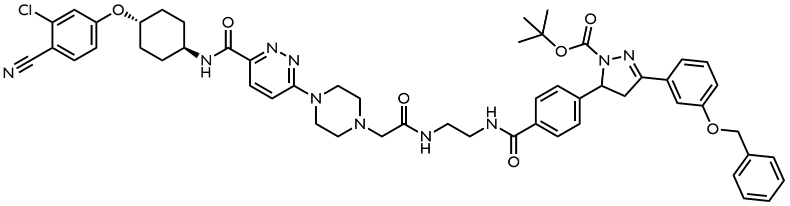

**30** was coupled to **9** following General Procedure E to yield **33** (37.8 mg, 0.04 mmol, 95%) as a yellow solid.
$^{1}$H NMR (600 MHz, $CDCl_3$) δ 8.01 (d, *J* = 9.5 Hz, 1H), 7.86 – 7.78 (m, 3H), 7.62 – 7.13 (m, 12H), 7.01 – 6.94 (m, 2H), 6.85 (dd, *J* = 8.8, 2.4 Hz, 1H), 5.38 (d, *J* = 7.1 Hz, 0H), 5.09 (s, 2H), 4.31 (tt, *J* = 9.9, 4.0 Hz, 1H), 4.05 (dtd, *J* = 10.8, 7.4, 3.9 Hz, 1H), 3.92 – 3.45 (m, 9H), 3.10 (q, *J* = 7.1 Hz, 3H), 2.69 (d, *J* = 10.5 Hz, 4H), 2.22 – 2.13 (m, 4H), 1.83 – 1.55 (m, 7H), 1.32 (s, 12H).

**tert-butyl 3-(3-(benzyloxy)phenyl)-5-(4-((4-(2-(4-(6-((4-(3-chloro-4-cyanophenoxy)cyclohexyl)carbamoyl)pyridazin-3-yl)piperazin-1-yl)acetamido)butyl)carbamoyl)phenyl)-4,5-dihydro-1*H*-pyrazole-1-carboxylate (34)**

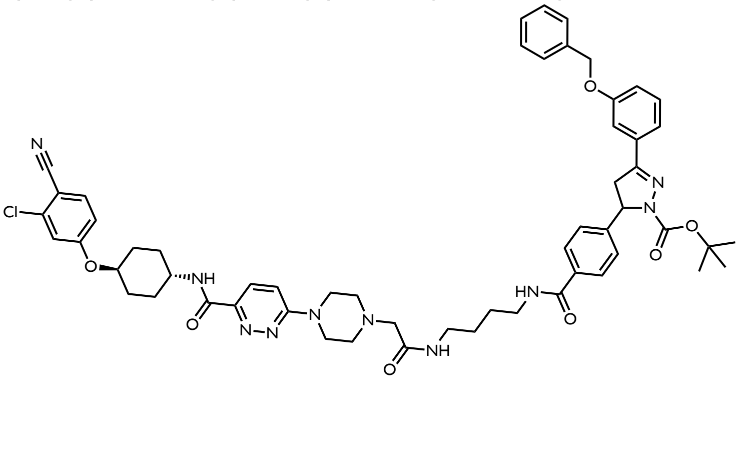

**31** was coupled to **9** following General Procedure E to yield **34** (21.6 mg, 0.021 mmol, 52.5%) as a yellow solid.
$^1$H NMR (600 MHz, $CDCl_3$) δ 8.01 (d, *J* = 9.5 Hz, 1H), 7.86 – 7.78 (m, 3H), 7.62 – 7.13 (m, 16H), 7.01 – 6.94 (m, 3H), 6.85 (dd, *J* = 8.8, 2.4 Hz, 1H), 5.38 (d, *J* = 7.1 Hz, 1H), 5.09 (s, 2H), 4.31 (tt, *J* = 9.9, 4.0 Hz, 1H), 4.05 (dtd, *J* = 10.8, 7.4, 3.9 Hz, 1H), 3.92 – 3.45 (m, 9H), 3.10 (q, *J* = 7.1 Hz, 3H), 2.69 (d, *J* = 10.5 Hz, 4H), 2.22 – 2.13 (m, 4H), 1.83 – 1.55 (m, 4H), 1.32 (s, 9H).

***tert*-butyl 3-(3-(benzyloxy)phenyl)-5-(4-((5-(2-(4-(6-((4-(3-chloro-4-cyanophenoxy)cyclohexyl)carbamoyl)pyridazin-3-yl)piperazin-1-yl)acetamido)pentyl)carbamoyl)phenyl)-4,5-dihydro-1*H*-pyrazole-1-carboxylate (35)**

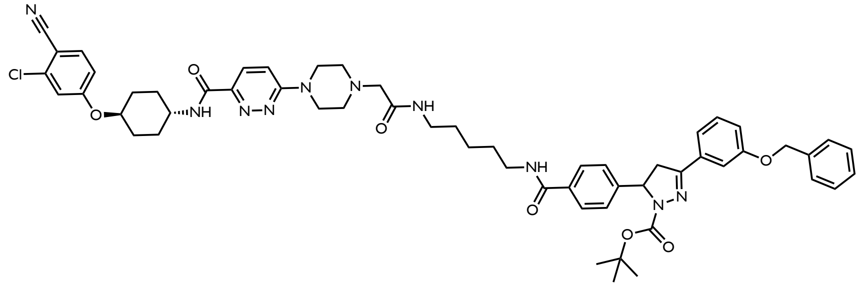

**32** was coupled to **9** following General Procedure E to yield **35** (46.3 mg, 0.045 mmol, 50%) as a yellow solid.
$^1$H NMR (600 MHz, $CDCl_3$) δ 8.06 – 7.99 (m, 1H), 7.82 – 7.72 (m, 3H), 7.49 (dd, *J* = 8.7, 1.7 Hz, 1H), 7.40 (d, *J* = 10.7 Hz, 1H), 7.19 (s, 16H), 7.05 – 6.84 (m, 3H), 6.78 (dd, *J* = 8.7, 2.3 Hz, 1H), 6.68 (s, 1H), 5.33 – 5.28 (m, 1H), 5.02 (d, *J* = 7.4 Hz, 2H), 4.25 (s, 1H), 4.02 – 3.05 (m, 14H), 2.12 (t, *J* = 14.9 Hz, 4H), 1.41 (d, *J* = 11.4 Hz, 8H), 1.27 (d, *J* = 8.4 Hz, 10H), 0.87 – 0.62 (m, 9H).

**General Procedure F: Preparation of chloroacetamide warhead containing AR ligand based bifunctionals.**

i. In a 20 mL scintillation vial, the boc protected amine (**33-35**) was dissolved in DCM. TFA was added dropwise at room temperature and the deprotection reaction was stirred for 30 minutes. The solvent was removed and washed with DCM 3 times. The DCM was removed in between each wash. The crude product was carried through to the next step without further purification.
ii. To a new reaction vial was charged the crude amine and dissolved in DCM. The solution was cooled to -10ºC. Chloroacetyl chloride (1.5 eq) was added dropwise before TEA (3 eq) was added and the reaction mixture was warmed to room temperature. The reaction was stirred for 30 minutes before it was concentrated *in vacuo* and purified via column chromatography (gradient elution, 0-10% methanol in DCM).

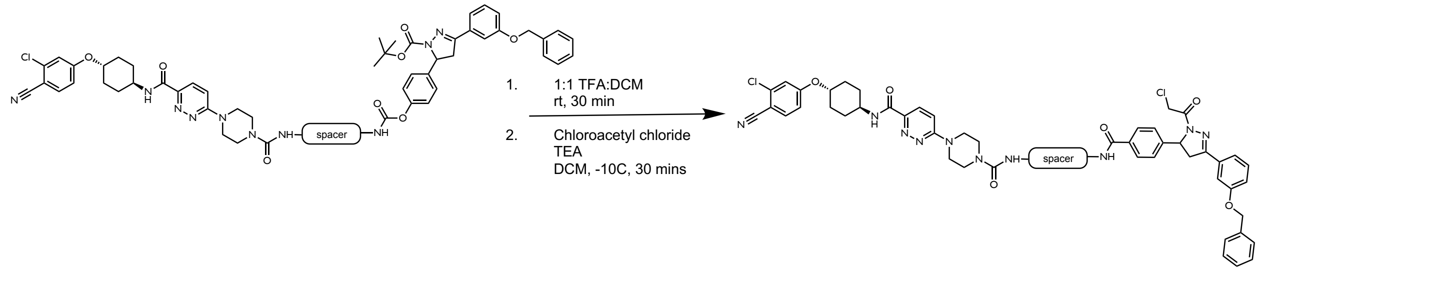


**6-(4-(2-((2-(4-(3-(3-(benzyloxy)phenyl)-1-(2-chloroacetyl)-4,5-dihydro-1*H*-pyrazol-5-yl)benzamido)ethyl)amino)-2-oxoethyl)piperazin-1-yl)-*N*-(4-(3-chloro-4-cyanophenoxy)cyclohexyl)pyridazine-3-carboxamide (36)**

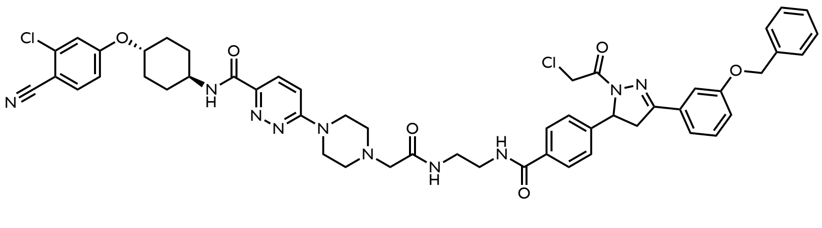

The warhead was added to **33** following General Procedure C to yield **36** (3.0 mg, 0.003 mmol, 7.9%) as a white solid.
$^1$H NMR (600 MHz, Acetone) δ 8.02 – 7.96 (m, 2H), 7.89 (s, 0H), 7.90 – 7.81 (m, 2H), 7.76 (d, *J* = 8.7 Hz, 1H), 7.54 – 7.45 (m, 3H), 7.42 – 7.28 (m, 7H), 7.26 (d, *J* = 2.5 Hz, 1H), 7.20 (d, *J* = 9.5 Hz, 1H), 7.18 – 7.09 (m, 2H), 5.67 (dd, *J* = 11.8, 5.1 Hz, 1H), 5.18 (s, 2H), 4.73 (d, *J* = 13.6 Hz, 1H), 4.65 – 4.56 (m, 2H), 4.03 – 3.93 (m, 2H), 3.74 (q, *J* = 4.5 Hz, 4H), 3.55 (q,

$J$ = 5.6 Hz, 2H), 3.52 – 3.41 (m, 2H), 3.30 – 3.23 (m, 1H), 3.00 (d, $J$ = 1.5 Hz, 2H), 2.61 (t, $J$ = 5.1 Hz, 4H), 2.25 – 2.18 (m, 2H), 1.75 – 1.59 (m, 4H).
$^{13}$C NMR (151 MHz, Acetone) δ 170.99, 167.44, 164.39, 163.39, 163.05, 161.42, 159.99, 156.19, 145.88, 145.85, 138.28, 138.09, 136.35, 135.02, 133.40, 130.79, 129.34, 128.77, 128.57, 128.55, 127.07, 126.68, 120.52, 118.10, 117.77, 116.86, 116.17, 113.96, 113.09, 105.07, 76.78, 70.65, 62.28, 61.20, 53.66, 48.14, 45.50, 43.00, 40.80, 39.98, 30.81; HRMS (ES+) calcd for $C_{51}H_{52}Cl_2N_{10}O_6Na$ [H+Na]$^+$ 993.034483, found 993.33215

**6-(4-(2-((4-(4-(3-(3-(benzyloxy)phenyl)-1-(2-chloroacetyl)-4,5-dihydro-1*H*-pyrazol-5-yl)benzamido)butyl)amino)-2-oxoethyl)piperazin-1-yl)-*N*-(4-(3-chloro-4-cyanophenoxy)cyclohexyl)pyridazine-3-carboxamide (37)**

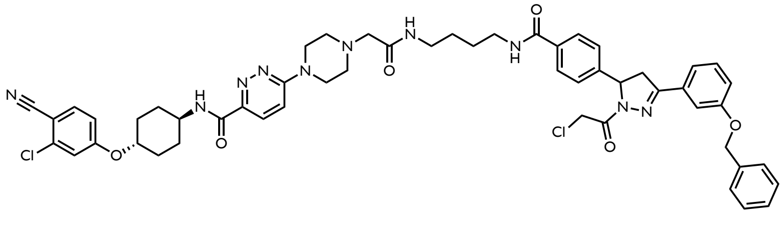

The warhead was added to **34** following General Procedure C to yield **37** (3.7 mg, 0.004 mmol, 18%) as a white solid. $^1$H NMR (600 MHz, Acetone) δ 7.98 (d, $J$ = 8.1 Hz, 1H), 7.93 – 7.87 (m, 1H), 7.87 – 7.79 (m, 3H), 7.77 (dd, $J$ = 8.7, 1.9 Hz, 1H), 7.57 – 7.21 (m, 13H), 7.17 – 7.11 (m, 2H), 5.67 (dd, $J$ = 11.8, 5.0 Hz, 1H), 5.19 (s, 2H), 4.72 (d, $J$ = 13.6 Hz, 1H), 4.68 – 4.56 (m, 2H), 4.06 – 3.94 (m, 2H), 3.86 – 3.74 (m, 5H), 3.41 (q, $J$ = 6.4 Hz, 2H), 3.32 – 3.21 (m, 3H), 3.00 (s, 2H), 2.66 (q, $J$ = 5.8 Hz, 5H), 2.29 – 2.21 (m, 3H), 1.77 – 1.55 (m, 9H), 1.28 (d, $J$ = 9.6 Hz, 4H). $^{13}$C NMR (151 MHz, Acetone) δ 168.92, 168.86, 166.07, 163.43, 162.47, 162.18, 160.59, 159.11, 155.24, 145.02, 144.77, 137.40, 137.22, 135.47, 134.57, 132.55, 129.89, 128.45, 127.88, 127.68, 127.62, 126.13, 125.70, 119.61, 117.21, 116.88, 115.95, 115.30, 113.07, 112.18, 104.19, 75.90, 69.77, 61.53, 60.30, 52.81, 52.40, 47.26, 47.15, 44.65, 42.10, 42.08, 39.18, 38.04, 37.92, 29.94, 29.75, 29.72, 29.44, 27.19, 26.77; HRMS (ES+) calcd for $C_{53}H_{56}Cl_2N_{10}O_6$ [M+H]$^+$ 999.37614, found 999.38324

**6-(4-(2-((5-(4-(3-(3-(benzyloxy)phenyl)-1-(2-chloroacetyl)-4,5-dihydro-1*H*-pyrazol-5-yl)benzamido)pentyl)amino)-2-oxoethyl)piperazin-1-yl)-*N*-(4-(3-chloro-4-cyanophenoxy)cyclohexyl)pyridazine-3-carboxamide (38)**

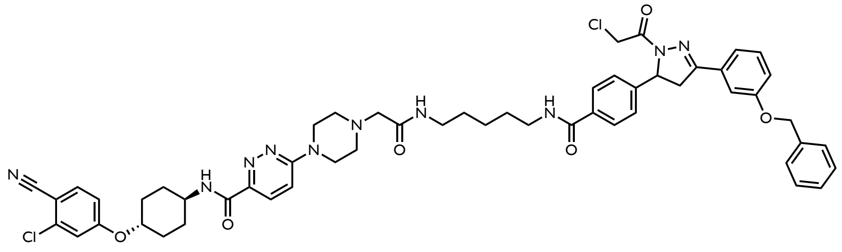

The warhead was added to **35** following General Procedure C to yield **38** (8.6 mg, 0.008 mmol, 19%) as a white solid. $^1$H NMR (600 MHz, Acetone) δ 7.91 – 7.82 (m, 3H), 7.75 (d, $J$ = 8.7 Hz, 1H), 7.54 – 7.46 (m, 3H), 7.43 – 7.29 (m, 3H), 7.28 – 7.20 (m, 2H), 7.17 – 7.09 (m, 2H), 5.66 (dd, $J$ = 11.9, 5.1 Hz, 1H), 5.18 (s, 2H), 4.72 (d, $J$ = 13.6 Hz, 1H), 4.65 – 4.56 (m, 2H), 4.22 (s, 1H), 4.02 – 3.92 (m, 2H), 3.77 (t, $J$ = 5.0 Hz, 4H), 3.38 (t, $J$ = 6.8 Hz, 2H), 3.30 – 3.20 (m, 3H), 3.00 (s, 2H), 2.62 (t, $J$ = 5.0 Hz, 4H), 2.25 – 2.18 (m, 2H), 1.75 – 1.49 (m, 9H), 1.47 – 1.33 (m, 2H). $^{13}$C NMR (151 MHz, Acetone) δ 169.02, 166.11, 163.49, 162.47, 162.16, 160.54, 159.10, 155.30, 144.98, 144.74, 137.40, 137.21, 135.48, 134.52, 132.51, 129.91, 128.46, 127.89, 127.69, 127.67, 126.17, 125.70, 119.63, 117.19, 116.89, 115.99, 115.28, 113.09, 112.23, 104.18, 75.88, 69.78, 61.41, 60.32, 52.77, 47.18, 44.57, 42.15, 42.12, 39.27, 38.17, 23.99; HRMS (ES+) calcd for $C_{54}H_{58}Cl_2N_{10}O_6$ [M+H]$^+$ 1013. 39179, found 1013. 40186

***tert*-butyl 3-(3-(benzyloxy)phenyl)-5-(4-(4-(6-((4-(3-chloro-4-cyanophenoxy)cyclohexyl)carbamoyl)pyridazin-3-yl)piperazine-1-carbonyl)phenyl)-4,5-dihydro-1*H*-pyrazole-1-carboxylate (39)**

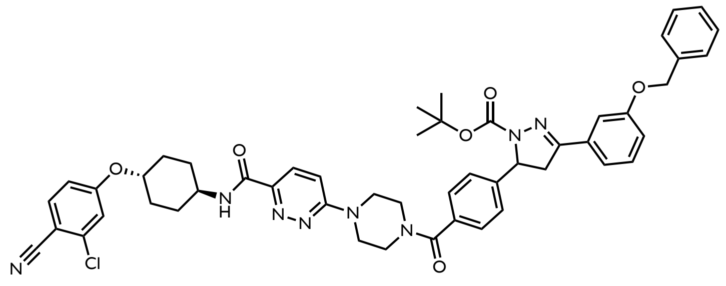

In a 20 mL scintillation vial, the boc protected amine (**28**) was dissolved in DCM. TFA was added dropwise at room temperature and the deprotection reaction was stirred for 30 minutes. Nitrogen was blown on the solution to evaporate the solvent and washed with DCM 3 times. The DCM was removed in between each wash. The crude product was carried through to the next step without further purification.
To a new reaction vial was charged the crude amine (1 eq), **16** (1 eq), and DCM (0.5M). HATU (2 eq) and DIPEA (5 eq) were added stepwise and the resulting reaction mixture was stirred for 20 mins at room temperature. The reaction was concentrated *in vacuo* and purified via column chromatography (gradient elution, 0-10% Methanol in DCM) to yield **39** (63 mg, 0.07 mmol, 74%) as a yellow solid.
$^1$H NMR (600 MHz, $CDCl_3$) δ 8.09 (d, *J* = 9.5 Hz, 1H), 7.87 (d, *J* = 8.1 Hz, 1H), 7.61 – 7.15 (m, 17H), 7.09 – 7.02 (m, 2H), 7.02 – 6.92 (m, 2H), 6.85 (dd, *J* = 8.7, 2.4 Hz, 1H), 5.43 – 5.39 (m, 1H), 5.10 (s, 2H), 4.31 (ddt, *J* = 9.9, 7.2, 3.9 Hz, 1H), 4.20 – 3.40 (m, 12H), 3.13 (d, *J* = 17.9 Hz, 1H), 2.22 – 2.13 (m, 4H), 1.52 – 1.42 (m, 4H), 1.37 (s, 9H).

**6-(4-(4-(3-(3-(benzyloxy)phenyl)-1-(2-chloroacetyl)-4,5-dihydro-1*H*-pyrazol-5-yl)benzoyl)piperazin-1-yl)-*N*-(4-(3-chloro-4-cyanophenoxy)cyclohexyl)pyridazine-3-carboxamide (40)**

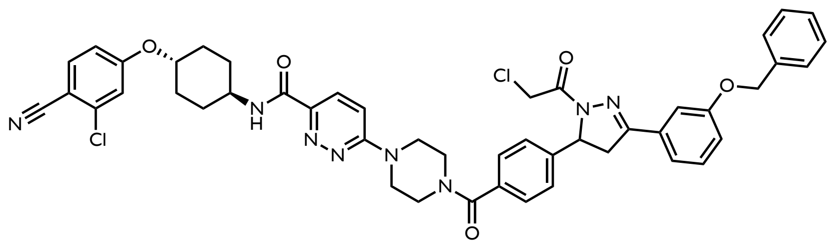

In a 20 mL scintillation vial, the boc protected amine (**39**) was dissolved in DCM. TFA was added dropwise at room temperature and the deprotection reaction was stirred for 30 minutes. The solvent was removed and washed with DCM 3 times. The DCM was removed in between each wash. The crude product was carried through to the next step without further purification. To a new reaction vial was charged the crude amine and dissolved in DCM. The solution was cooled to -10°C. Chloroacetyl chloride (1.5 eq) was added dropwise before TEA (3 eq) was added and the reaction mixture was warmed to room temperature. The reaction was stirred for 30 minutes before it was concentrated *in vacuo* and purified via column chromatography (gradient elution, 0-10% methanol in DCM) to yield **40** (2.4 mg, 0.003 mmol, 43%) as a yellow solid.
$^1$H NMR (600 MHz, Acetone) δ 7.99 (d, *J* = 8.2 Hz, 1H), 7.93 (d, *J* = 9.5 Hz, 1H), 7.77 (dd, *J* = 8.8, 0.8 Hz, 1H), 7.56 – 7.26 (m, 15H), 7.19 – 7.12 (m, 2H), 5.71 (dd, *J* = 11.8, 4.9 Hz, 1H), 5.20 (s, 2H), 4.74 (d, *J* = 13.7 Hz, 1H), 4.68 – 4.60 (m, 2H), 4.06 – 3.96 (m, 2H), 3.34 – 3.28 (m, 1H), 2.77 (s, 2H), 2.25 (dt, *J* = 11.9, 3.8 Hz, 2H), 2.15 – 2.07 (m, 2H), 1.78 – 1.62 (m, 4H). $^{13}$C NMR (151 MHz, Acetone) δ 169.29, 163.45, 162.37, 162.31, 162.19, 160.55, 159.12, 155.31, 145.30, 143.51, 137.40, 137.23, 135.46, 132.57, 131.13, 129.89, 128.75, 128.46, 127.88, 127.79, 127.68, 126.23, 125.78, 119.61, 117.22, 116.87, 115.94, 115.31, 113.10, 112.49, 104.19, 75.90, 69.77, 67.44, 60.27, 47.28, 47.18, 42.16, 42.13, 38.79, 31.74, 30.29, 29.95, 29.74, 29.71, 29.44, 23.62, 22.75, 22.43, 13.42, 10.42; HRMS (ES+) calcd for $C_{47}H_{44}Cl_2N_8O_5$ [M+Na]$^+$ 893.28117, found 893.27014

AP-01-98R-H.4.fid
1H starting parameters - HC 06/17/2019
No decoupling

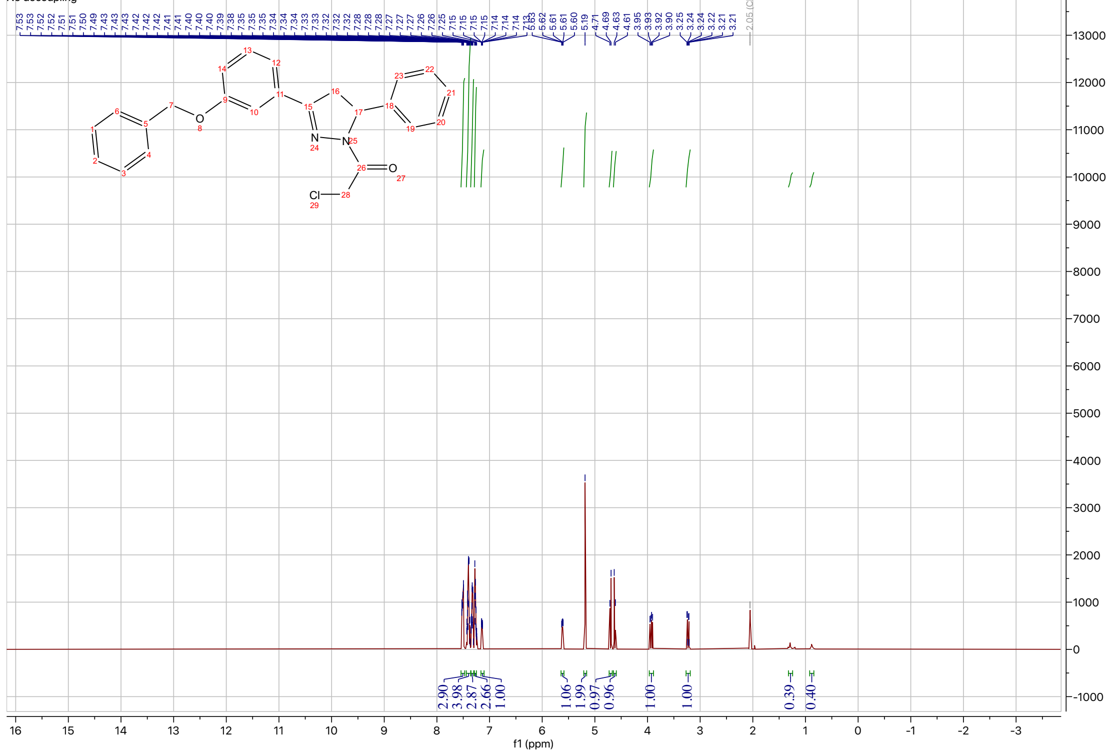

AP-01-98R-C.5.fid
13C{1H} starting parameters - HC 06/17/2019

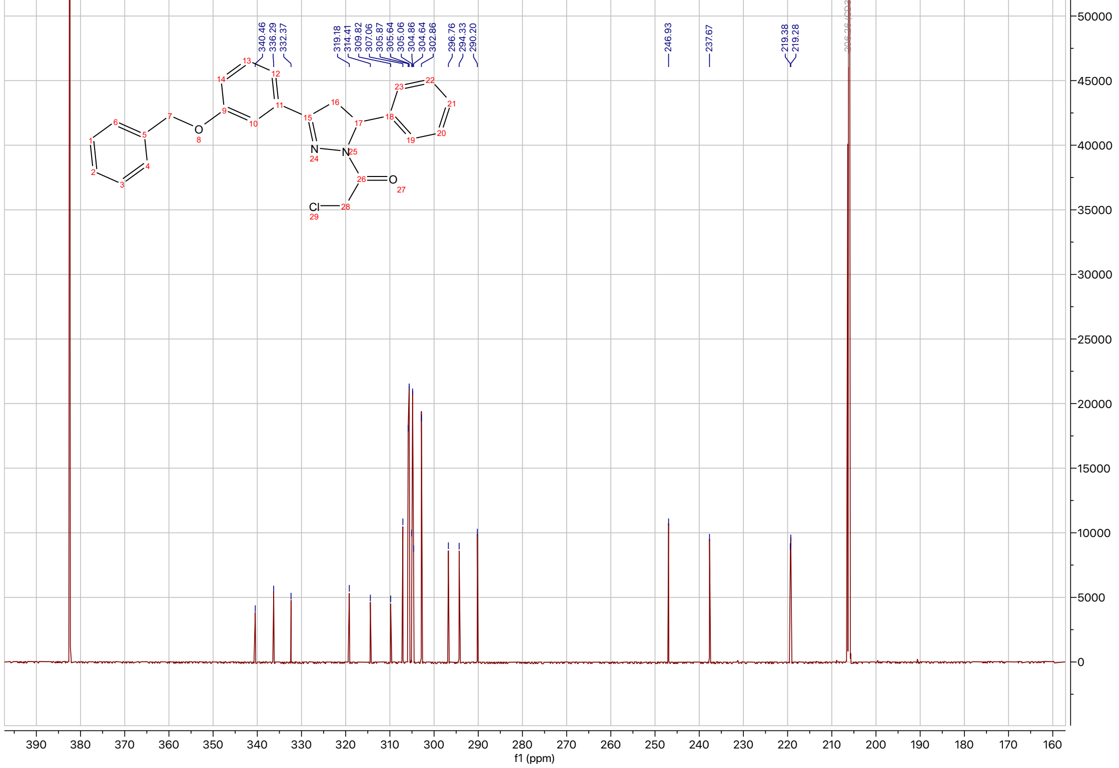

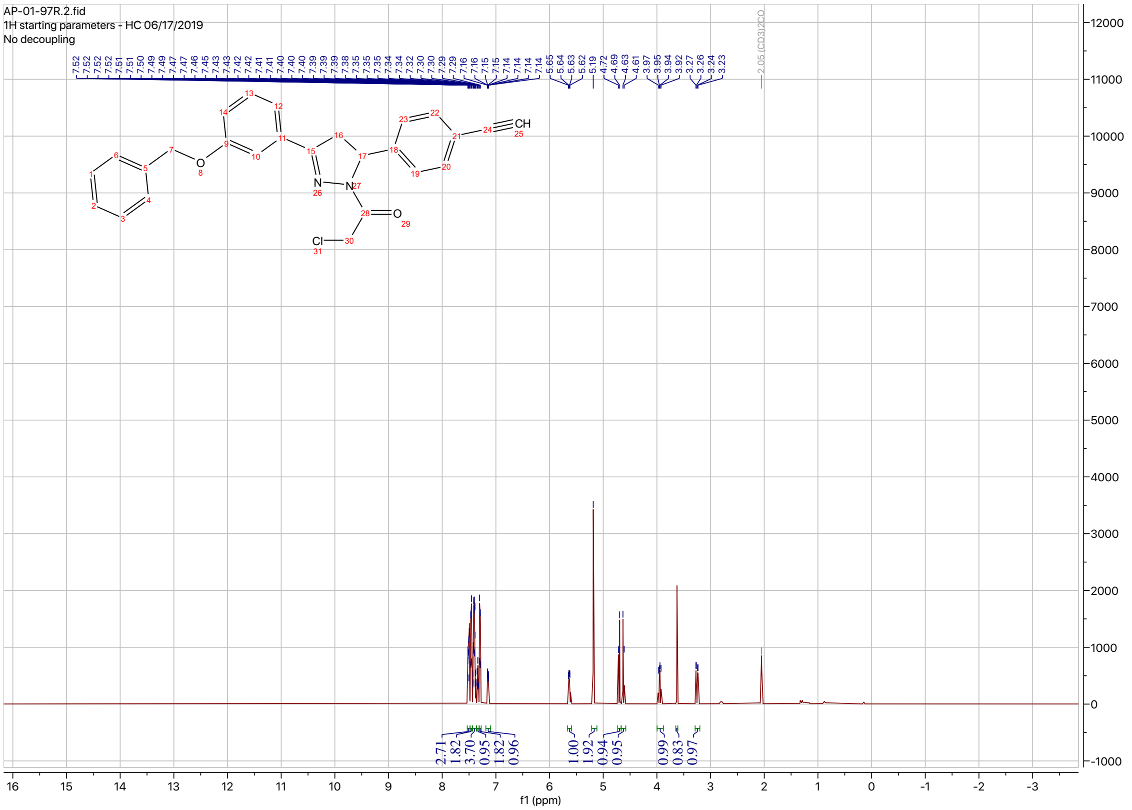
AP-01-97R.2.fid
1H starting parameters - HC 06/17/2019
No decoupling
2.05 (CD3)2CO
f1 (ppm)

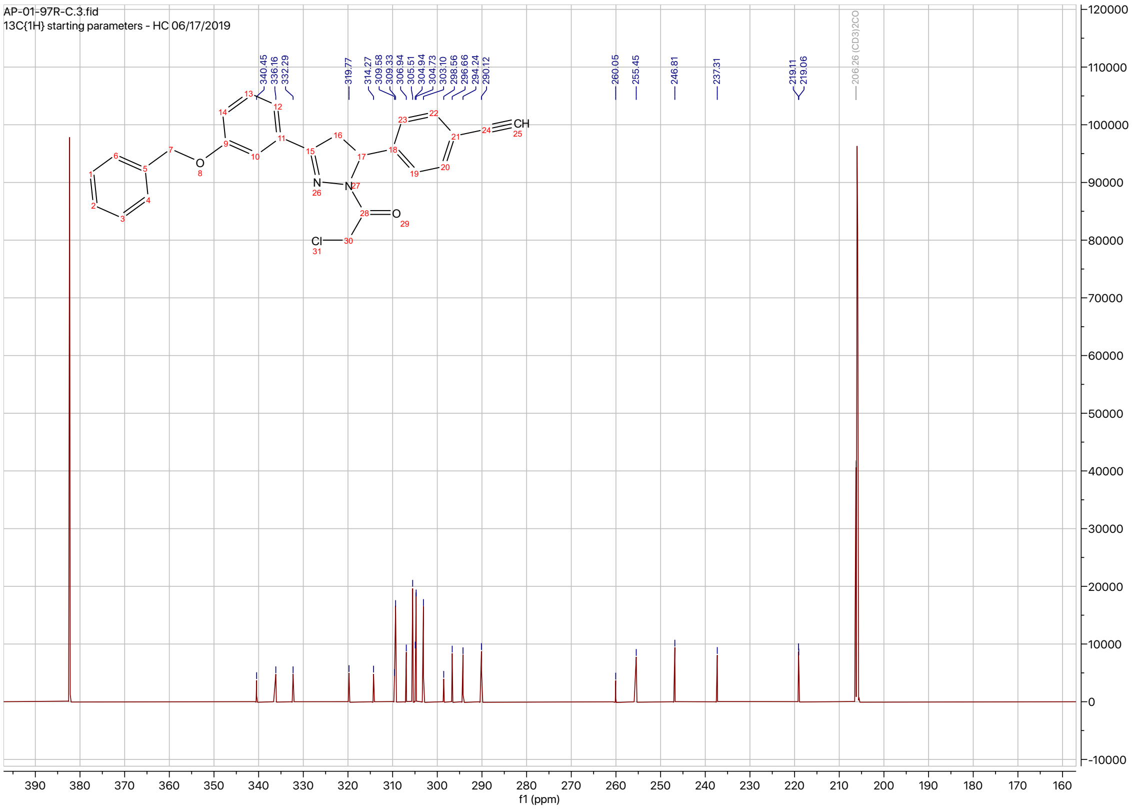
AP-01-97R-C.3.fid
13C{1H} starting parameters - HC 06/17/2019
206.26 (CD3)2CO
f1 (ppm)

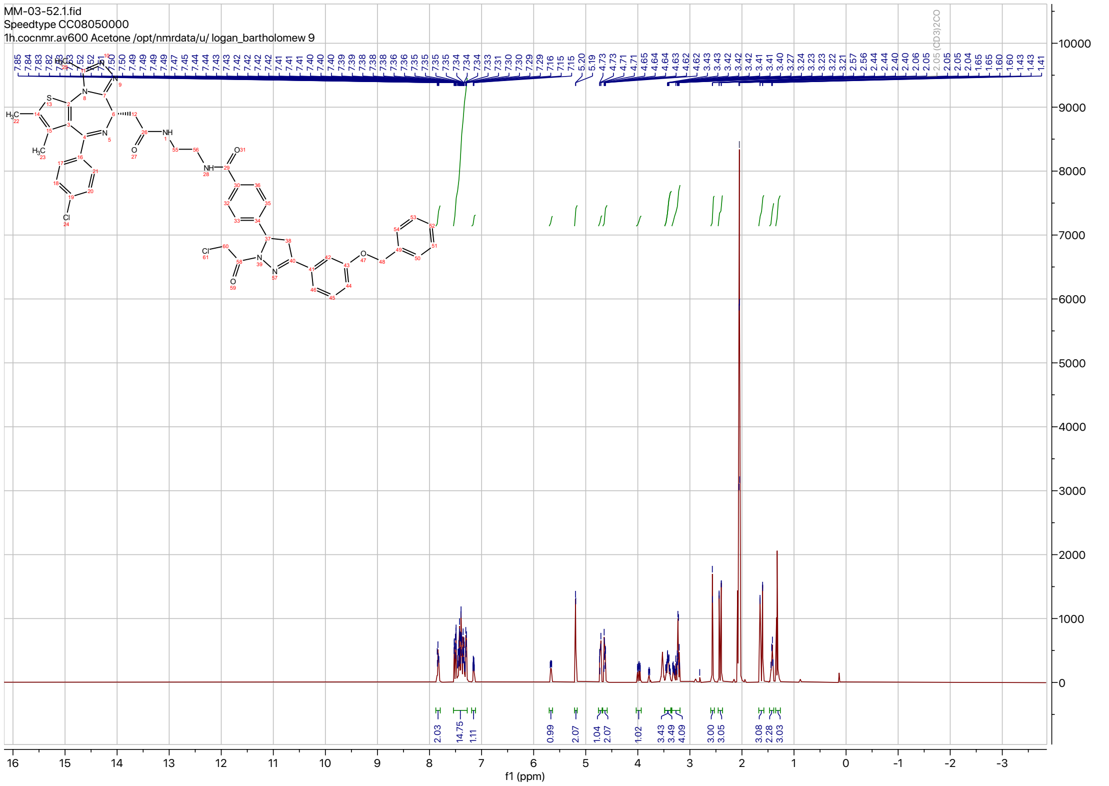

MM-03-52.1.fid
Speedtype CC08050000
1h.cocnmr.av600 Acetone /opt/nmrdata/u/ logan_bartholomew 9
f1 (ppm)


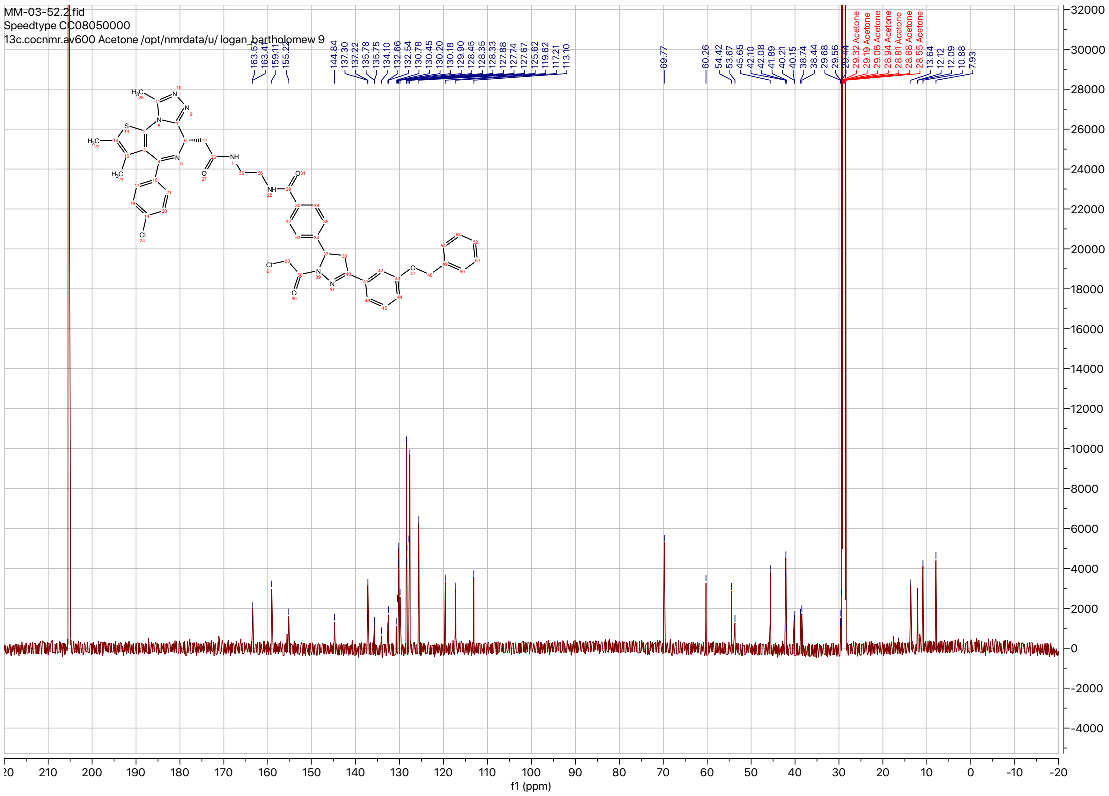

MM-03-52.2.fid
Speedtype CC08050000
13c.cocnmr.av600 Acetone /opt/nmrdata/u/ logan_bartholomew 9
f1 (ppm)

MM-03-41-H.13.fid
Speedtype CC08050010

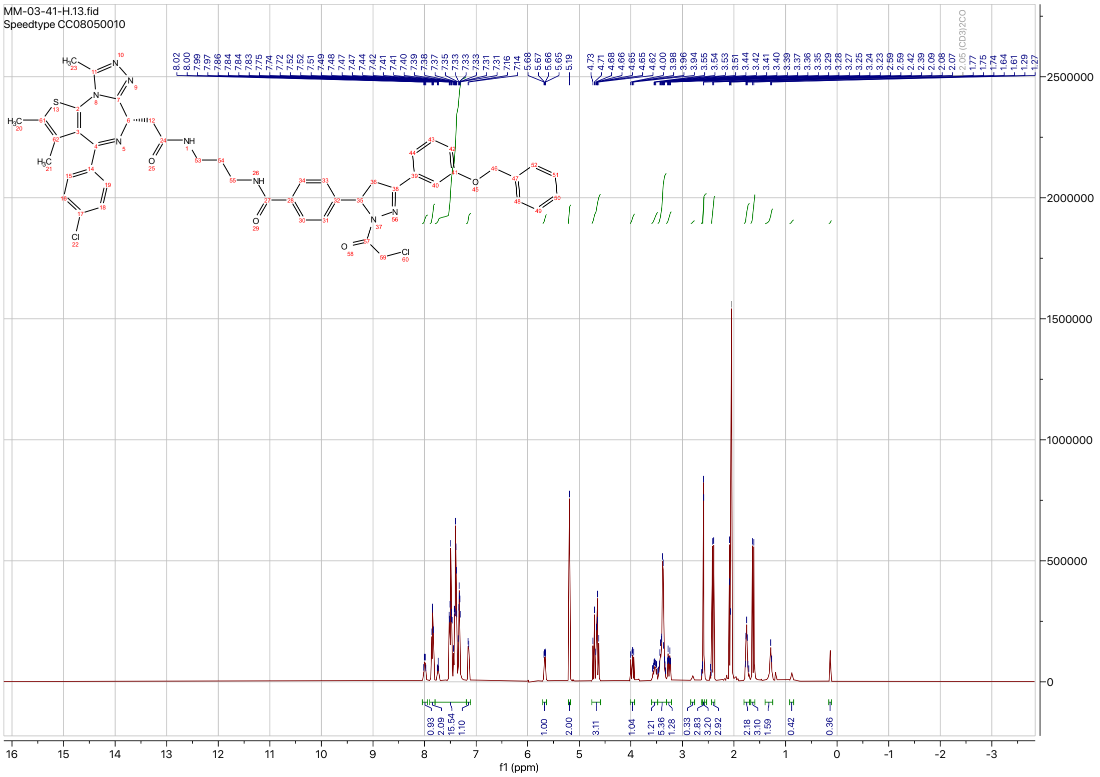

MM-03-41-C.12.fid
Speedtype CC08050010

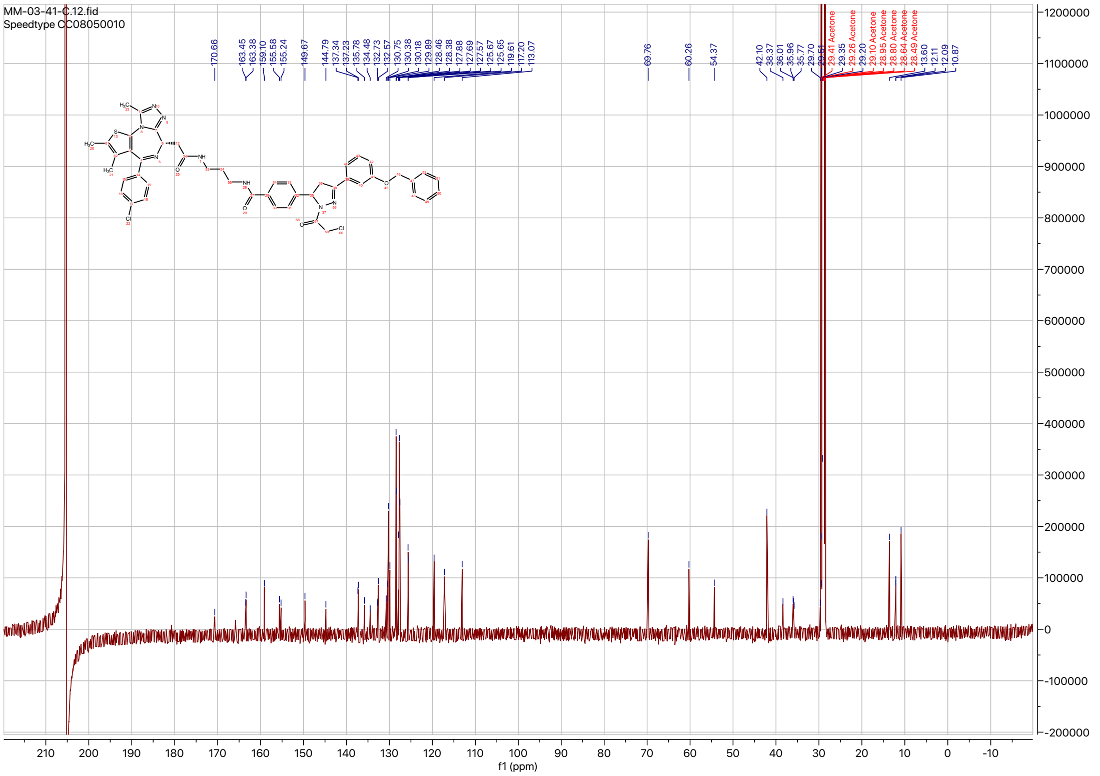

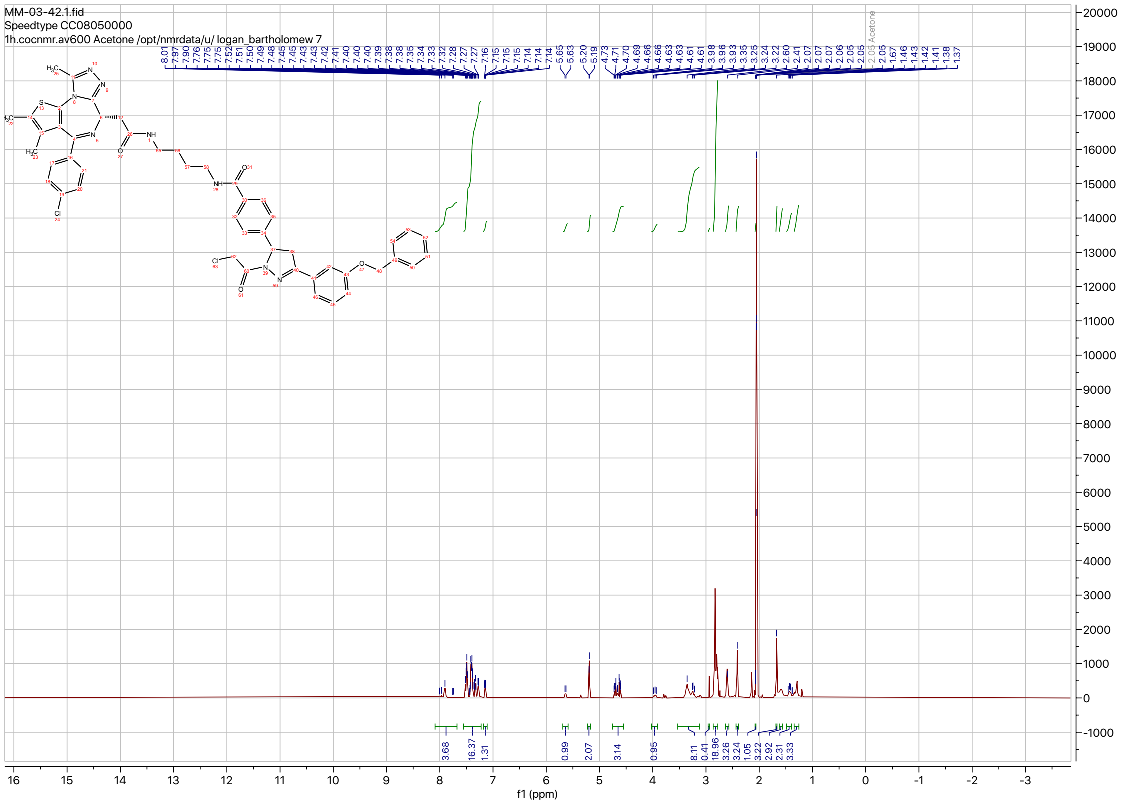
MM-03-42.1.fid
Speedtype CC08050000
1h.cocnmr.av600 Acetone /opt/nmrdata/u/ logan_bartholomew 7
8.01 7.97 7.90 7.76 7.75 7.75 7.52 7.51 7.50 7.49 7.48 7.45 7.43 7.43 7.42 7.41 7.40 7.40 7.39 7.38 7.38 7.35 7.34 7.33 7.32 7.28 7.27 7.27 7.16 7.15 7.15 7.14 7.14 7.14 5.65 5.63 5.20 5.19 4.73 4.71 4.70 4.69 4.66 4.66 4.63 4.63 4.61 4.61 3.98 3.96 3.93 3.35 3.25 3.24 3.22 2.60 2.41 2.07 2.07 2.07 2.06 2.05 2.05 2.05 2.05 Acetone 2.05 1.67 1.46 1.43 1.42 1.41 1.38 1.37
3.68 16.37 1.31 0.99 2.07 3.14 0.95 8.11 0.41 18.96 3.26 3.24 1.05 3.22 2.92 2.31 3.33
f1 (ppm)

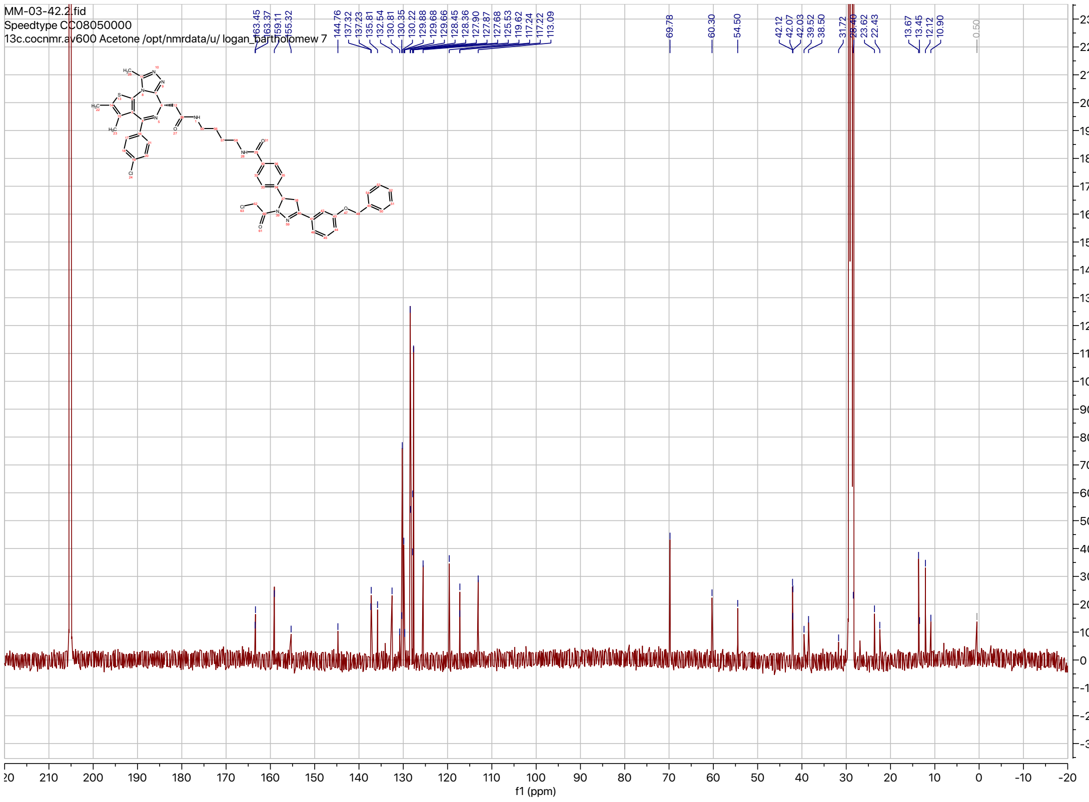
MM-03-42.2.fid
Speedtype CC08050000
13c.cocnmr.av600 Acetone /opt/nmrdata/u/ logan_bartholomew 7
163.45 163.37 159.11 155.32 144.76 137.32 137.23 135.81 132.54 130.81 130.35 130.22 129.88 129.66 129.68 128.45 128.36 127.90 127.87 127.68 125.53 119.62 117.24 117.22 113.09 69.78 60.30 54.50 42.12 42.07 42.03 39.52 38.50 31.72 28.40 23.62 22.43 13.67 13.45 12.12 10.90 0.50
f1 (ppm)

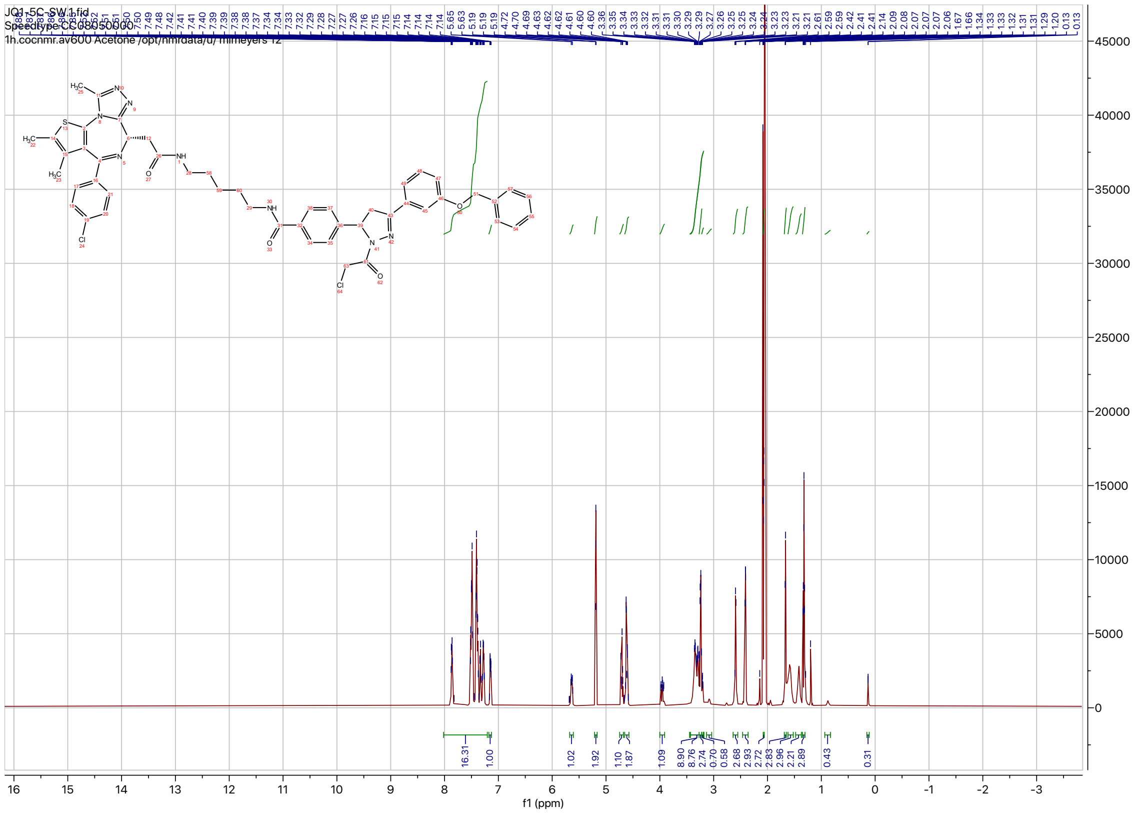
JQ1-5C-SW.1.fid
Speedtype CC08050000
1h.cocnmr.av600 Acetone /opt/nmrdata/u/ mlmeyers 12
f1 (ppm)

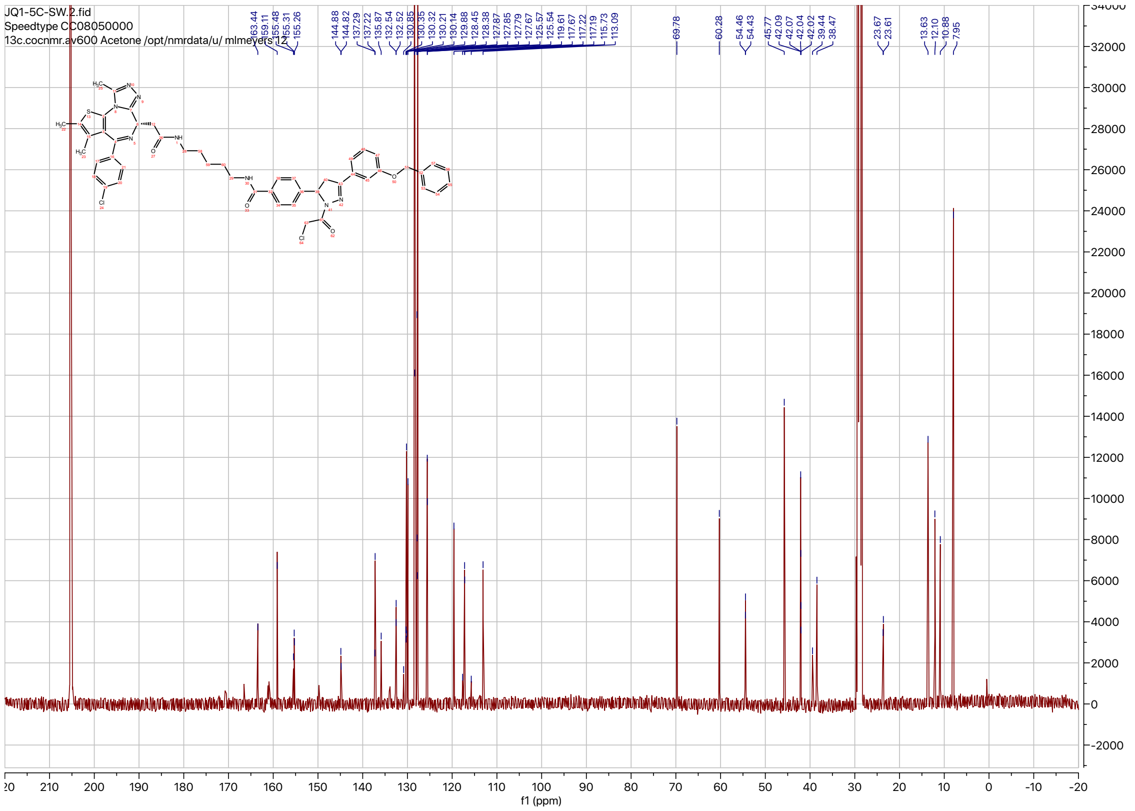
JQ1-5C-SW.2.fid
Speedtype CC08050000
13c.cocnmr.av600 Acetone /opt/nmrdata/u/ mlmeyers 12
f1 (ppm)

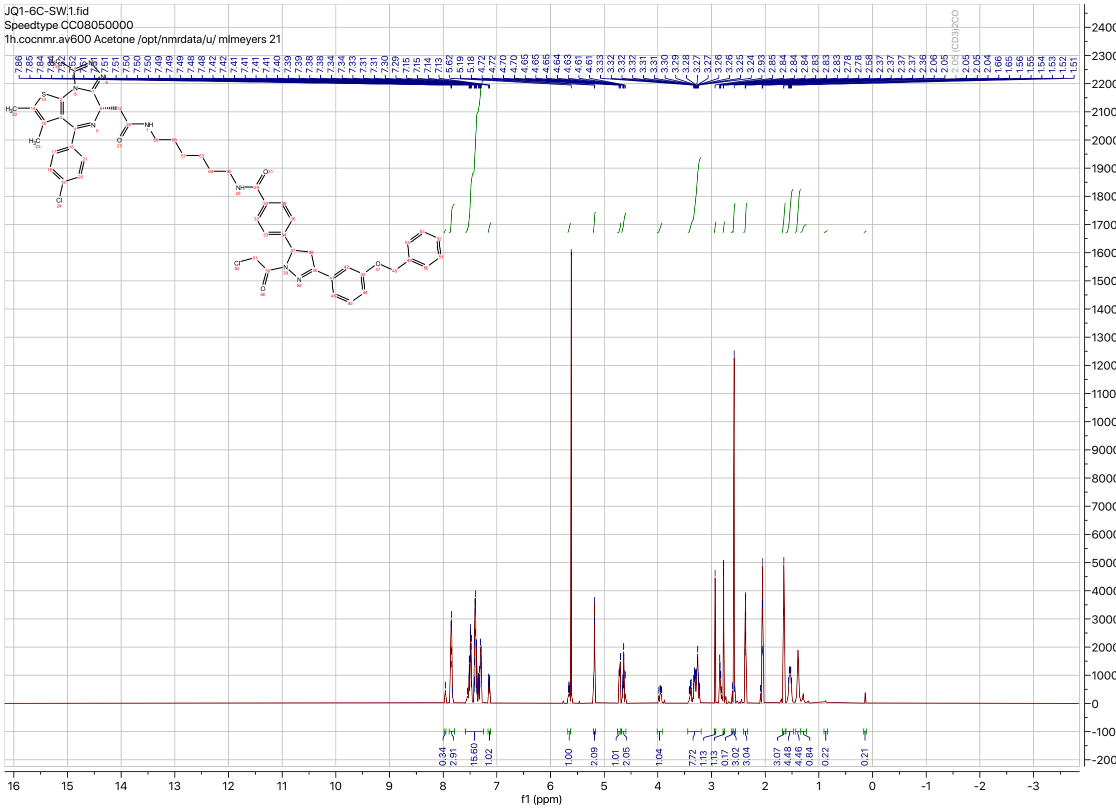
JQ1-6C-SW.1.fid
Speedtype CC08050000
1h.cocnmr.av600 Acetone /opt/nmrdata/u/ mlmeyers 21
0.34
2.91
15.60
1.02
1.00
2.09
1.01
2.05
1.04
7.72
1.13
0.17
3.02
3.04
3.07
4.48
4.46
0.84
0.22
0.21
f1 (ppm)

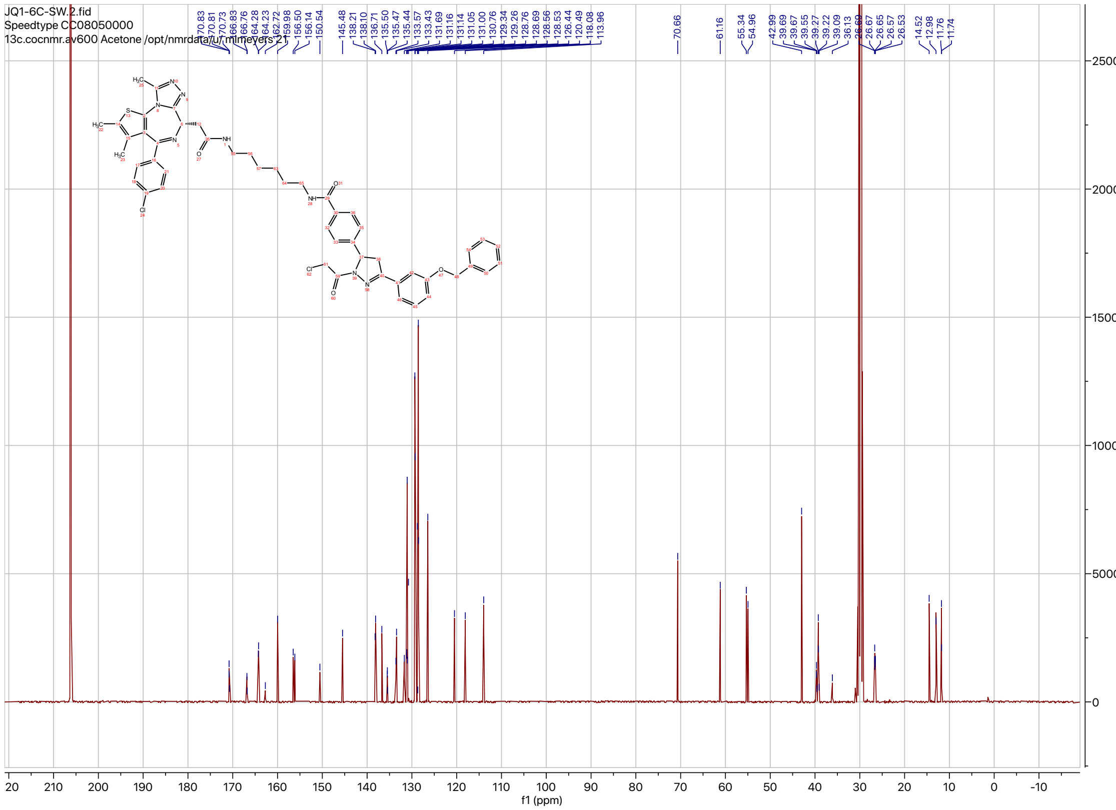
JQ1-6C-SW.2.fid
Speedtype CC08050000
13c.cocnmr.av600 Acetone /opt/nmrdata/u/ mlmeyers 21
170.83
170.81
170.73
166.83
166.76
164.28
164.23
162.72
159.98
156.50
156.14
150.54
145.48
138.21
138.10
136.71
135.50
135.47
135.44
133.57
133.43
131.69
131.16
131.14
131.05
131.00
130.76
129.34
129.26
128.76
128.69
128.56
128.53
126.44
120.49
118.08
113.96
70.66
61.16
55.34
54.96
42.99
39.69
39.67
39.55
39.27
39.22
39.09
36.13
26.67
26.65
26.57
26.53
14.52
12.98
11.76
11.74
f1 (ppm)

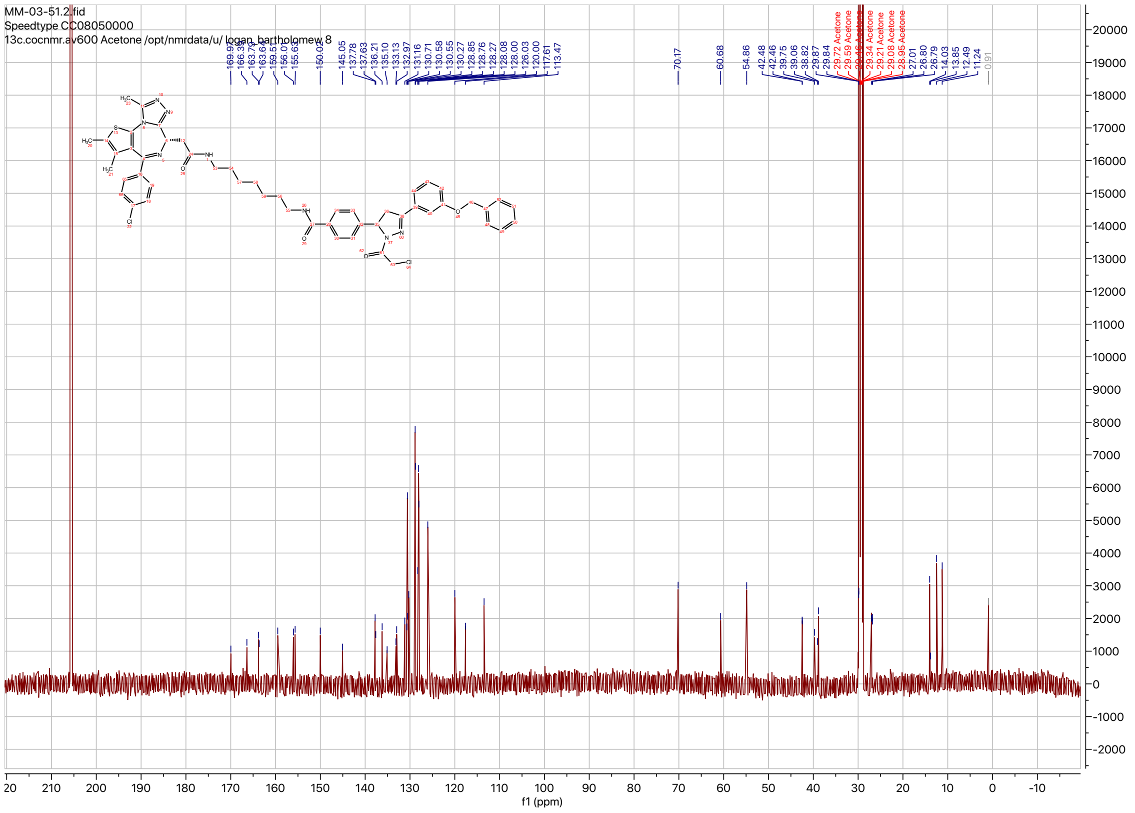
MM-03-51.2.fid
Speedtype CC08050000
13c.cocnmr.av600 Acetone /opt/nmrdata/u/ logan_bartholomew 8
f1 (ppm)

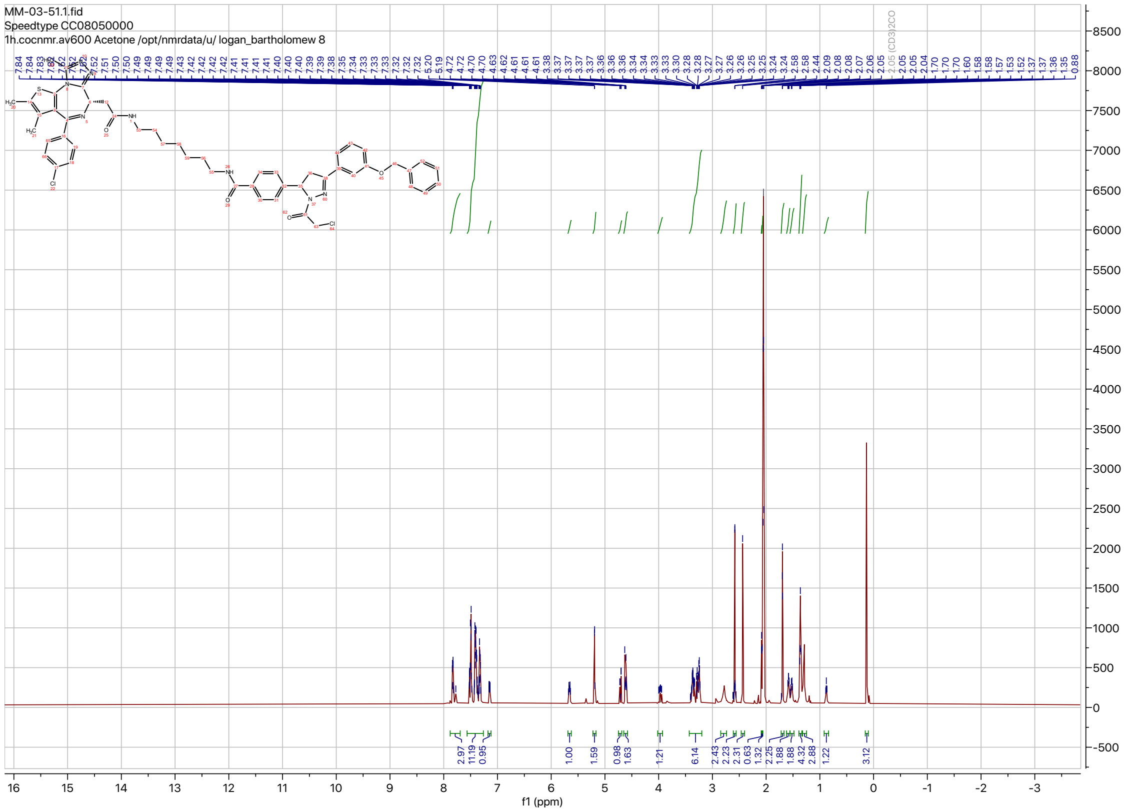
MM-03-51.1.fid
Speedtype CC08050000
1h.cocnmr.av600 Acetone /opt/nmrdata/u/ logan_bartholomew 8
f1 (ppm)

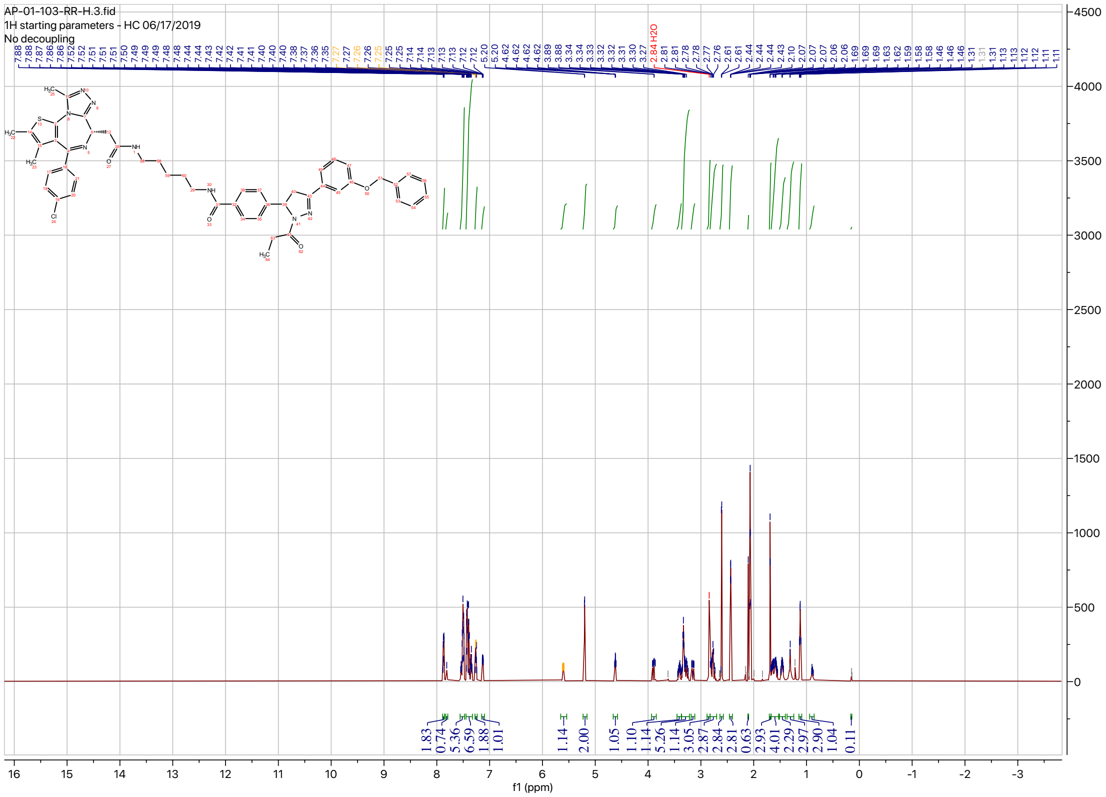
AP-01-103-RR-H.3.fid
1H starting parameters - HC 06/17/2019
No decoupling
f1 (ppm)

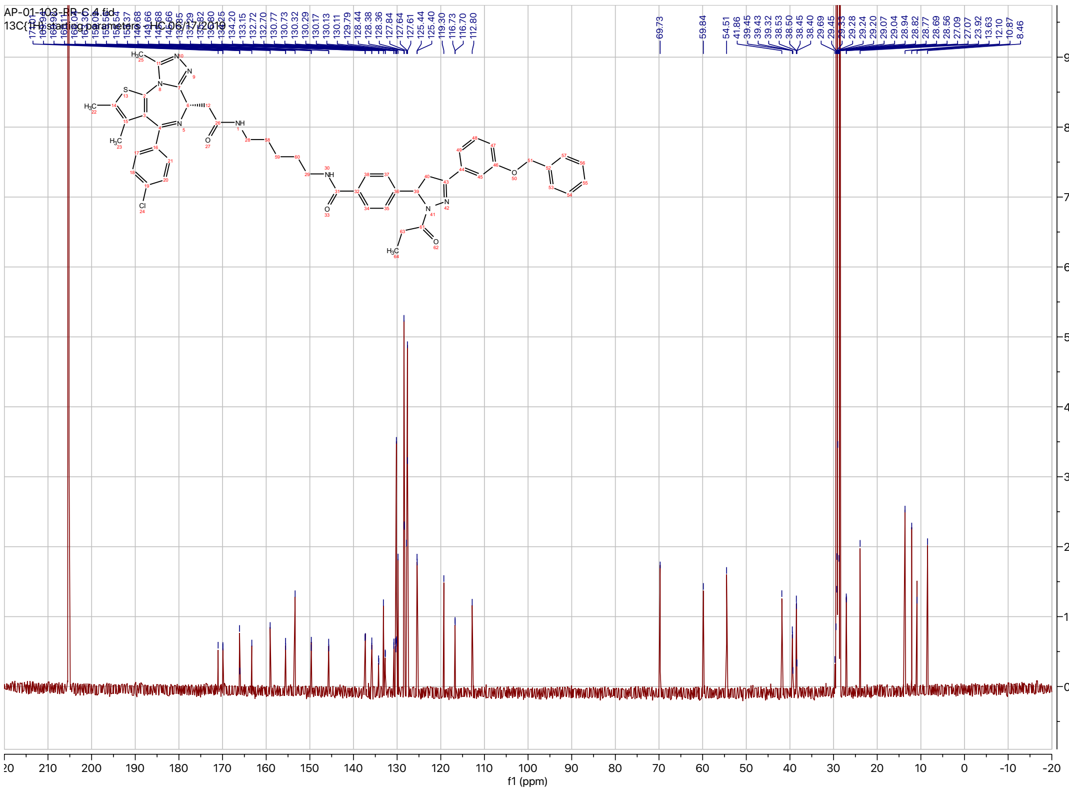
AP-01-103-RR-C.4.fid
13C{1H} starting parameters - HC 06/17/2019
f1 (ppm)

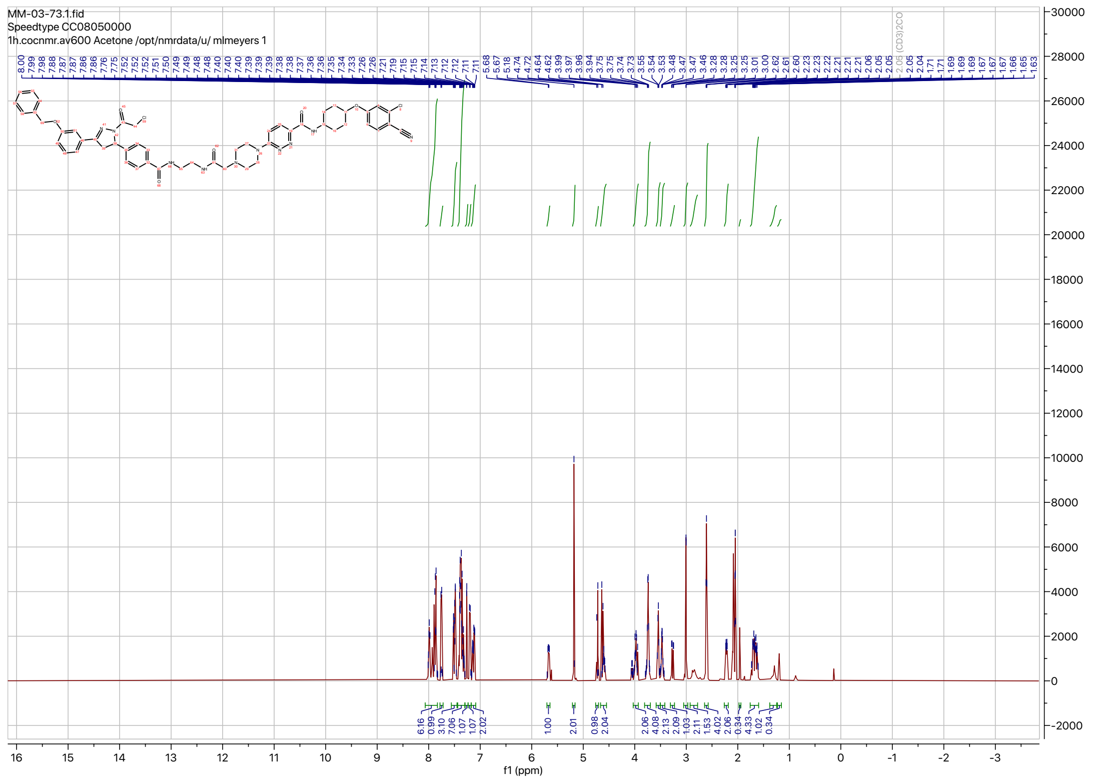
MM-03-73.1.fid
Speedtype CC08050000
1h.cocnmr.av600 Acetone /opt/nmrdata/u/ mlmeyers 1
f1 (ppm)

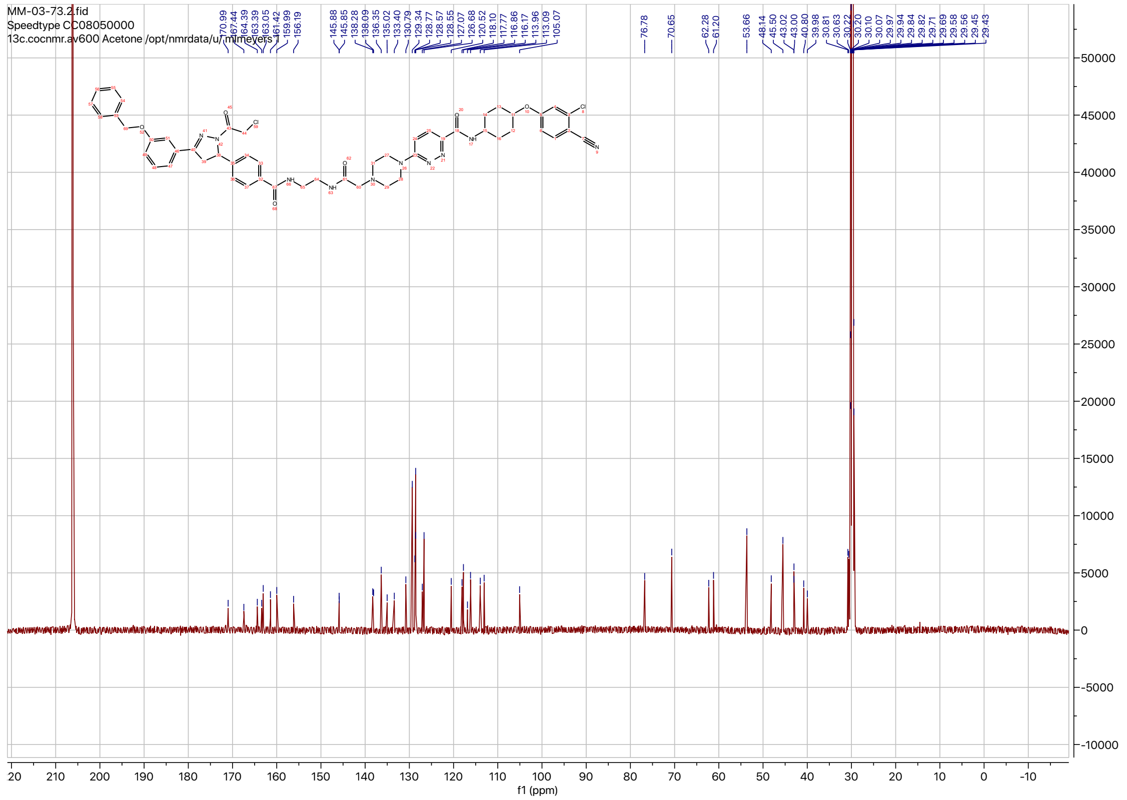
MM-03-73.2.fid
Speedtype CC08050000
13c.cocnmr.av600 Acetone /opt/nmrdata/u/ mlmeyers 1
f1 (ppm)

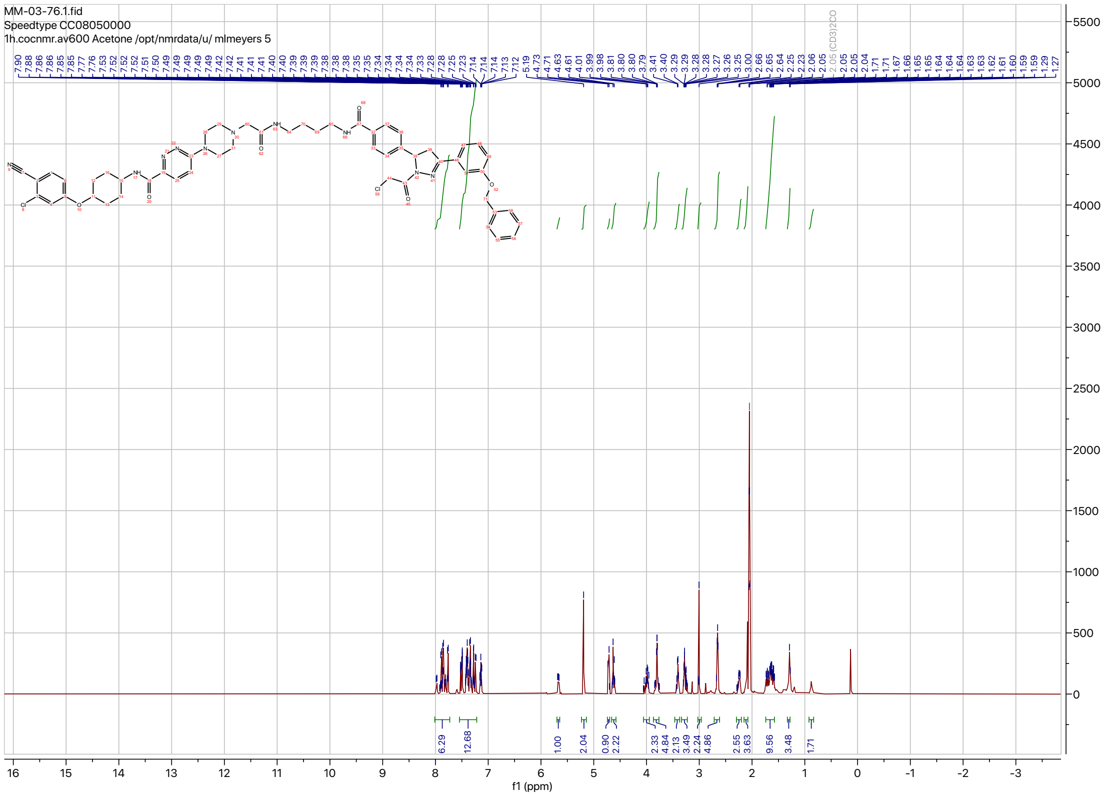
MM-03-76.1.fid
Speedtype CC08050000
1h.cocnmr.av600 Acetone /opt/nmrdata/u/ mlmeyers 5
7.90 7.88 7.86 7.85 7.77 7.76 7.53 7.52 7.52 7.51 7.50 7.49 7.49 7.49 7.49 7.42 7.42 7.41 7.41 7.40 7.40 7.39 7.39 7.39 7.38 7.38 7.38 7.35 7.35 7.34 7.34 7.34 7.34 7.33 7.28 7.28 7.25 7.23 7.14 7.14 7.14 7.14 7.13 7.12 5.19 4.73 4.71 4.63 4.61 4.01 3.99 3.98 3.81 3.80 3.80 3.79 3.41 3.40 3.29 3.29 3.29 3.28 3.28 3.27 3.26 3.25 3.00 2.66 2.65 2.64 2.25 2.23 2.06 2.05 2.05 (CD3)2CO 2.05 2.05 2.04 1.71 1.71 1.67 1.66 1.65 1.65 1.64 1.64 1.63 1.63 1.62 1.61 1.60 1.59 1.59 1.29 1.27
6.29 12.68 1.00 2.04 0.90 2.22 2.33 4.84 2.13 3.49 2.24 4.86 2.55 3.63 9.56 3.48 1.71
f1 (ppm)

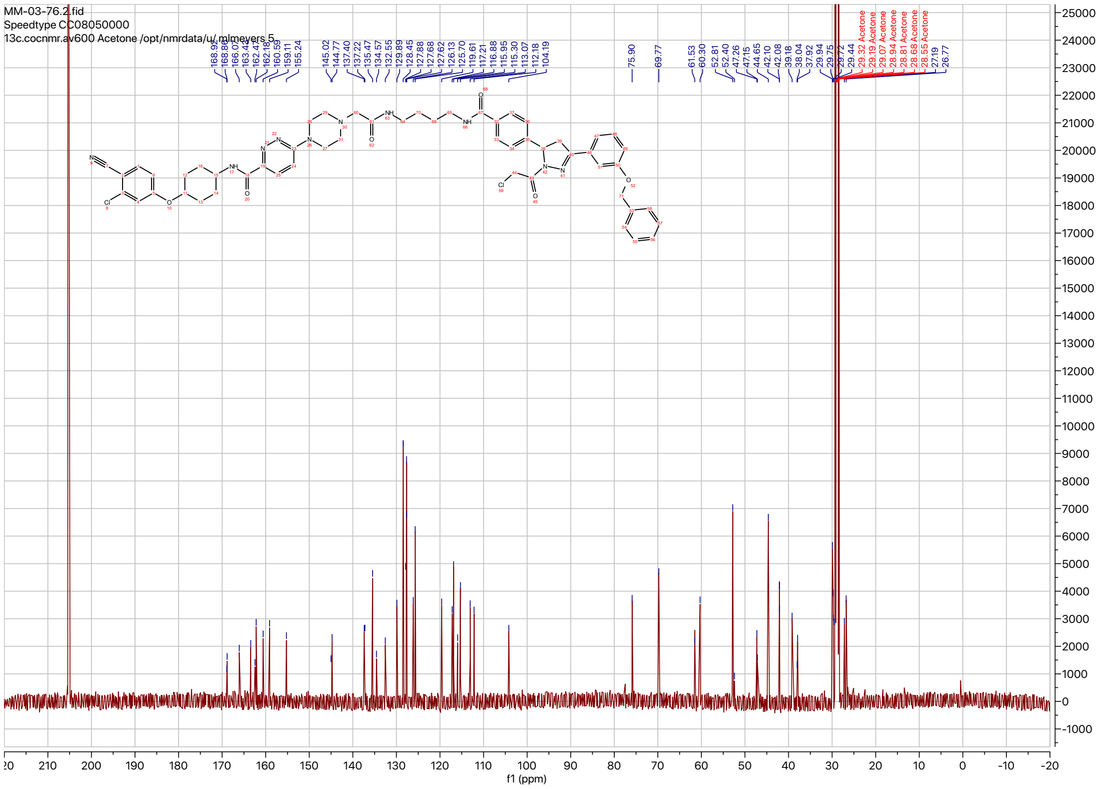
MM-03-76.2.fid
Speedtype CC08050000
13c.cocnmr.av600 Acetone /opt/nmrdata/u/ mlmeyers 5
168.92 168.89 166.07 163.42 162.47 162.16 160.59 159.11 155.24 145.02 144.77 137.40 137.22 135.47 134.57 132.55 129.89 128.45 127.88 127.68 127.62 126.13 125.70 119.61 117.21 116.88 115.95 115.30 113.07 112.18 104.19 75.90 69.77 61.53 60.30 52.81 52.40 47.26 47.15 44.65 42.10 42.08 39.18 38.04 37.92 29.94 29.75 29.72
29.44 Acetone 29.32 Acetone 29.19 Acetone 29.07 Acetone 28.94 Acetone 28.81 Acetone 28.68 Acetone 28.55 Acetone
27.19 26.77
f1 (ppm)

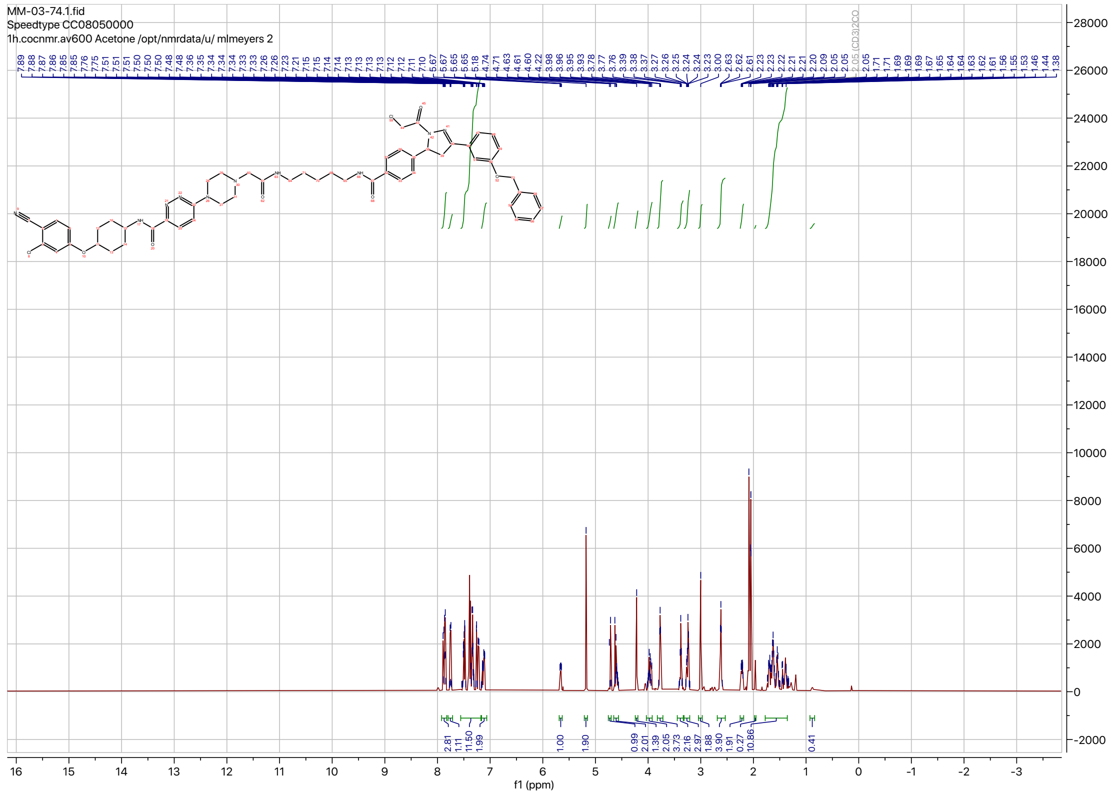
MM-03-74.1.fid
Speedtype CC08050000
1h.cocnmr.av600 Acetone /opt/nmrdata/u/ mlmeyers 2
7.89 7.88 7.86 7.85 7.85 7.76 7.75 7.51 7.51 7.50 7.50 7.48 7.48 7.36 7.35 7.34 7.34 7.33 7.33 7.26 7.26 7.23 7.21 7.15 7.15 7.14 7.14 7.13 7.13 7.13 7.13 7.12 7.11 7.10 5.67 5.67 5.65 5.65 5.18 4.74 4.71 4.63 4.61 4.60 4.22 3.98 3.96 3.95 3.93 3.78 3.77 3.76 3.39 3.38 3.37 3.27 3.25 3.24 3.24 3.23 3.00 2.63 2.62 2.61 2.23 2.23 2.22 2.21 2.21 2.20 2.09 2.05 2.05 (CD3)2CO 2.05 1.71 1.71 1.69 1.69 1.67 1.65 1.64 1.64 1.63 1.62 1.61 1.56 1.53 1.46 1.44 1.38
2.81 1.11 11.50 1.99 1.00 1.90 0.99 2.01 1.39 2.05 3.73 2.16 2.97 1.88 3.90 1.91 0.27 10.86 0.41
f1 (ppm)

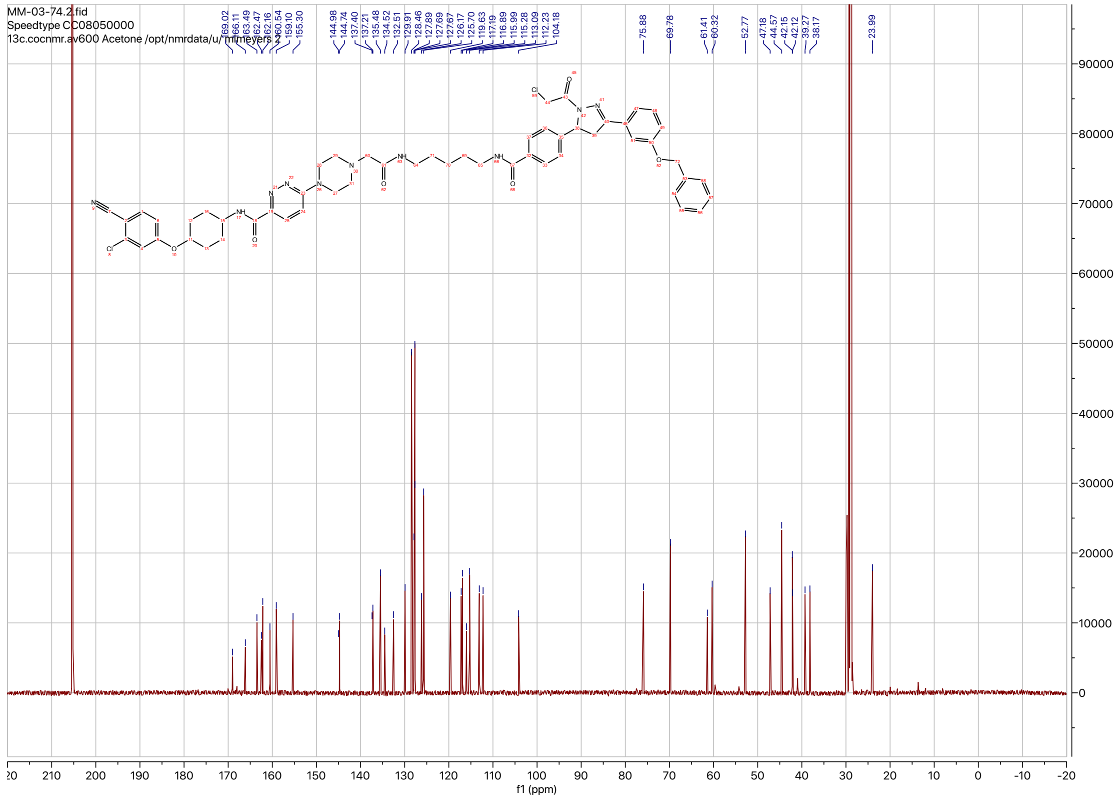
MM-03-74.2.fid
Speedtype CC08050000
13c.cocnmr.av600 Acetone /opt/nmrdata/u/ mlmeyers 2
169.02 166.11 163.49 162.47 162.16 160.54 159.10 155.30 144.98 144.74 137.40 137.21 135.48 134.52 132.51 129.91 128.46 127.89 127.69 127.67 126.17 125.70 119.63 117.19 116.89 115.99 115.28 113.09 112.23 104.18 75.88 69.78 61.41 60.32 52.77 47.18 44.57 42.15 42.12 39.27 38.17 23.99
f1 (ppm)

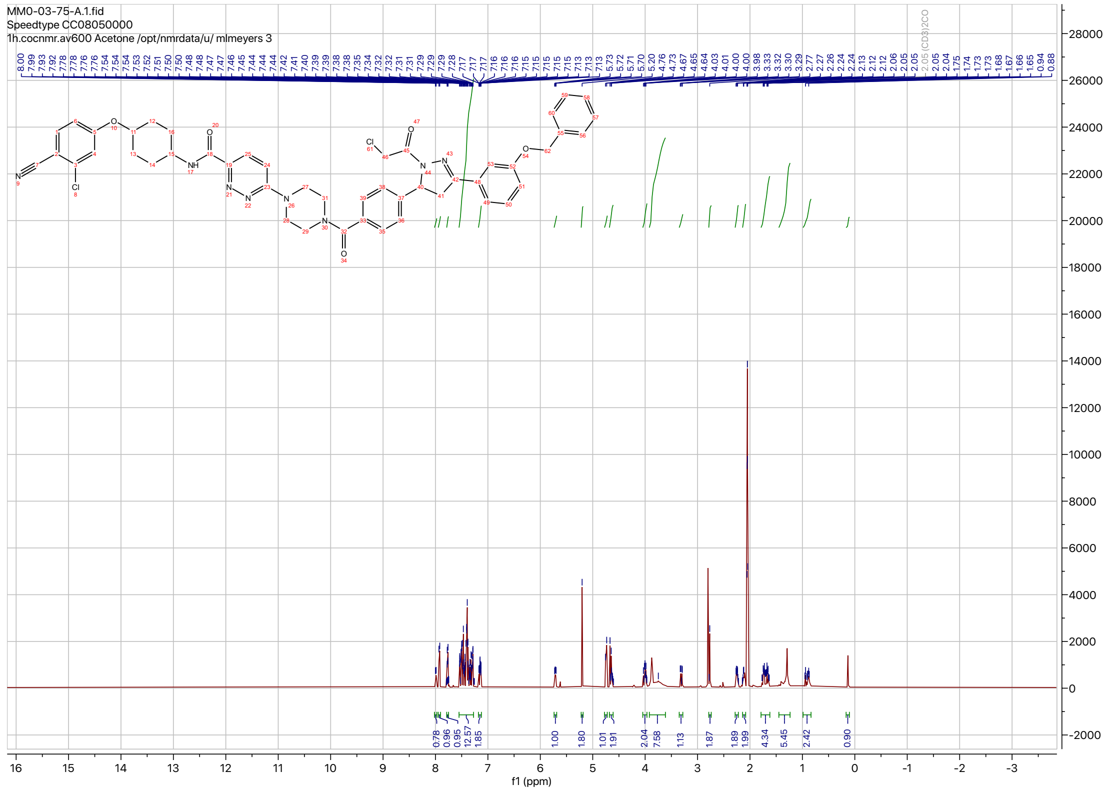
MM0-03-75-A.1.fid
Speedtype CC08050000
1h.cocnmr.av600 Acetone /opt/nmrdata/u/ mlmeyers 3
f1 (ppm)

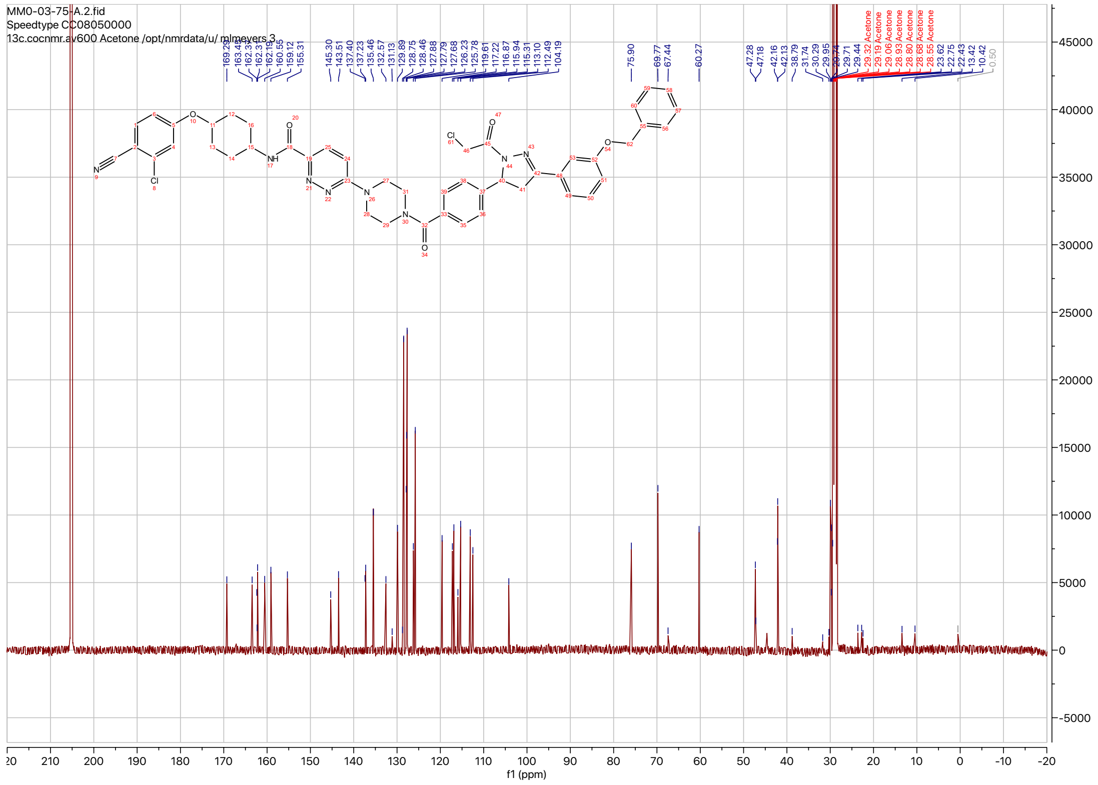
MM0-03-75-A.2.fid
Speedtype CC08050000
13c.cocnmr.av600 Acetone /opt/nmrdata/u/ mlmeyers 3
f1 (ppm)

MM-04-09 post hplc.6.fid
13C{1H} starting parameters - HC 06/17/2019

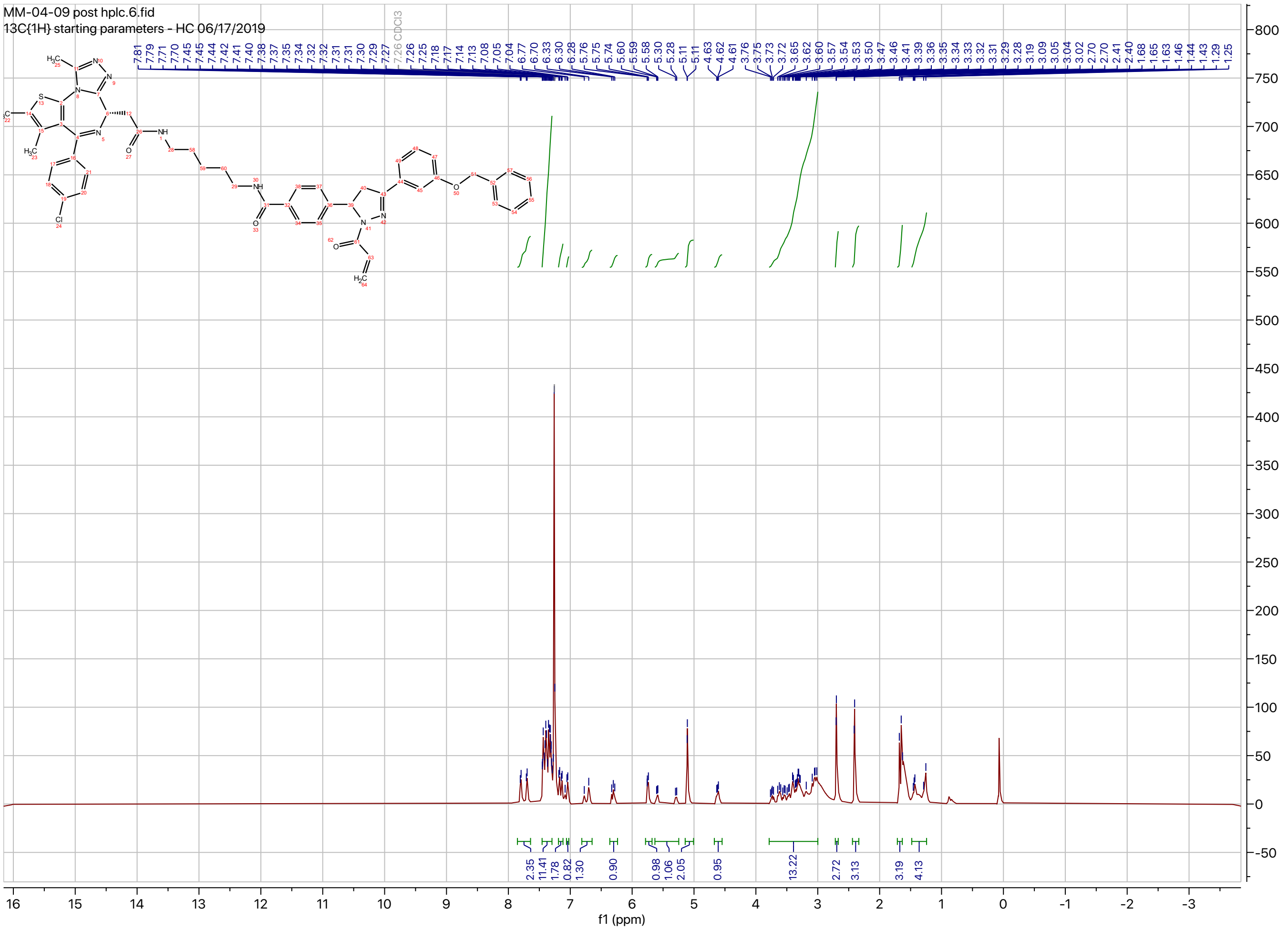
